# Population-based structure modeling reveals key roles of nuclear microenviroment in gene functions

**DOI:** 10.1101/2021.07.11.451976

**Authors:** Asli Yildirim, Nan Hua, Lorenzo Boninsegna, Guido Polles, Ke Gong, Shengli Hao, Wenyuan Li, Xianghong Jasmine Zhou, Frank Alber

## Abstract

The nuclear folding of chromosomes relative to nuclear bodies is an integral part of gene function. Here, we demonstrate that population-based modeling—from ensemble Hi-C data—can provide a detailed description of the nuclear microenvironment of genes and its role on gene function. We define the microenvironment by the subnuclear positions of genomic regions with respect to nuclear bodies, local chromatin compaction, and preferences in chromatin compartmentalization. These structural descriptors are determined in single cell models on a genome-wide scale, thereby revealing the structural variability between cells. We demonstrate that the structural microenvironment of a genomic region is linked to its functional potential in gene transcription, replication and chromatin compartmentalization. Some chromatin regions are distinguished by their strong preferences to a single microenvironment, due to associations to specific nuclear bodies in most cells. Other chromatin shows high structural variability, which is a strong indicator of functional heterogeneity. Moreover, we identify specialized nuclear microenvironments, which distinguish chromatin in different functional states and reveal a key role of nuclear speckles in chromosome organization. We demonstrate that our method produces highly predictive 3-dimensional genome structures, which accurately reproduce data from TSA-seq, DamID, GPSeq and super-resolution imaging. Thus, our method considerably expands the range of Hi-C data analysis and is widely applicable.

## Introduction

The spatial organization of eukaryotic genomes is linked to regulation of gene transcription, DNA replication, cell differentiation and upon malfunction to cancer and other diseases^1,2^. Recent advances have led to a prolific development of improved technologies for probing chromosome interactions and 3D organization^3,4^. Live-cell and super-resolution microscopy^5–11^ as well as mapping technologies based on high-throughput sequencing^12–27^ shed light into the dynamic formation of chromatin loops and topological associating domains (TADs). These structural elements play a role in moderating promoter-enhancer interactions between remote DNA regions for regulating gene expression^28–30^. However, besides local promoter-enhancer interactions, gene expression and other functions are also influenced by their nuclear locations and chromatin compartmentalization, i.e., preferential associations of chromatin with similar functional profiles^31,32^. Chromosome conformation mapping and imaging^8,10,11,33^ show spatial segregation of chromatin into transcriptionally active and inactive A/B compartments^21^, subsequently refined, at high sequencing depth, into 5 primary Hi-C subcompartments^34^. Chromatin compartmentalization is also instigated by associations to nuclear bodies, such as nuclear speckles, PML bodies, Polycomb bodies or lamina associating domains, and other nuclear compartments^32^. Transcriptional permissive regions often locate at nuclear speckles, nuclear pore complexes and PML bodies, while regions of transcriptional repression are associated with the nuclear lamina and perinucleolar chromatin^35^. Thus, gene positions to nuclear bodies can play critical roles in permissiveness of gene expression and other functions^35,36^.

However, mapping the three-dimensional (3D) organization of all genes in single cells remains a major challenge. Several experimental technologies probe the mean distances (TSA-seq^14^) or association frequencies (NAD-seq^37^, DamID^17^) of genes to nuclear speckles, lamina associated domains, and nucleoli. However, collecting this information simultaneously within the same cell, at the same time, is challenging, especially when considering cell-to-cell variability of a gene’s microenvironment within a population of cells. Several super-resolution microscopy techniques have recently provided such information^7–9^. For instance, DNA- and RNA-multiplexed error-robust fluorescence *in situ* hybridization (MERFISH) super-resolution imaging detected, within the same cells, the nuclear locations of 1,137 genes, together with the positions of nuclear speckles, nucleoli, as well as the amount mRNA transcripts^8^. However, at this point, the amount of probed genomic DNA regions is still sparse, containing ~1% of entire genomes.

Here, we introduce an approach for modeling a population of single cell 3D genome structures to describe the nuclear microenvironment of genomic regions on a genome-wide scale. Our aim is to evaluate the roles of the nuclear architecture and its cell-to-cell variability in genome function and identify specialized nuclear microenvironments, which distinguish chromatin in different functional states.

We achieve this goal by using a population-based genome structure modeling approach, which takes *in situ* Hi-C data to generate a population of diploid genome structures statistically consistent with it^38,39^. We demonstrate that our method produces—from Hi-C data alone—highly predictive genome structures, which accurately predict data from SON TSA-seq^14^, lamin B1 TSA-seq^14^, lamin B1 pA-DamID^40^, GPSeq^41^, 3D fluorescence *in situ* hybridization (FISH)^19^ and DNA-MERFISH^8^ experiments. We define the nuclear microenvironment of genomic regions by an array of structural descriptors, including radial positions, association frequencies and mean distances to nuclear speckles, lamin B1, and nucleoli, the local chromatin fiber compaction, and local compartmentalization in form of the trans A/B ratio^8^ (**Fig. 1a,b**). These structural descriptors are determined in single cell models, thereby revealing cell-to-cell variability of structures across the population of models.

Our analysis provides several key findings. Firstly, genomic regions with stable structural properties, thus a strong preference in their nuclear microenvironment are most homogenous in their functional properties across cells in a population. For instance, genes with high cell-to-cell heterogeneity in expression^42^ often show increased structural variability, indicating a contribution of extrinsic noise to gene expression heterogeneity^43^. Chromatin with low structural variability are associated with either nuclear speckles or constitutive lamina associated domains (LADs) in the majority of cells. These regions provide structural anchor points for other chromatin and thus are a major factor in genome organization. We also observe nuclear zones around speckles to be hubs of inter-chromosomal interactions in the active compartment. Secondly, our analysis shows that the subnuclear microenvironment of a genomic region reflects its transcriptional potential. Genes with highest expression levels can be distinguished from those with lowest based on their structural microenvironment. Among all structural descriptors, the speckle association frequency and trans A/B ratio have the highest predictive value for its gene expression potential. Thirdly, the nuclear microenvironment of a genomic region is a good indicator of its replication timing. Moreover, our observations also confirm that Hi-C subcompartments^34^ define physically distinct chromatin environments, some of which (like A1) linked to associations with nuclear bodies. Interestingly, the A2 subcompartment stands out by its high structural variability between cells, a feature distinctly different from the A1 subcompartment.

Although other computational approaches also modeled entire chromosomes or even diploid genomes from Hi-C data^19,39,44–60^ so far, none documented the predictive accuracy in reproducing multimodal experimental data as presented here. Our findings demonstrate that our approach, from Hi-C data alone, produces exceedingly predictive models, providing a detailed description of the subnuclear locations, folding and compartmentalization of chromatin in diploid genomes. Therefore, our approach considerably expands the scope of information retrieved from Hi-C data and is widely applicable to any cell type for which Hi-C data is available.

## Results

### Assessment of 3D genome structures

Here, we study 3D structures of diploid lymphoblastoid genomes (GM12878) from *in situ* Hi-C data^34^ at 200 kb (kilobase) resolution. Our method generates a population of 10,000 genome structures, in which all accumulated chromatin contacts are statistically consistent with contact probabilities from Hi-C experiments^38,39,61^. The structure optimization is achieved by solving a maximum likelihood estimation problem utilizing an iterative optimization algorithm with a series of optimization strategies for efficient and scalable model estimation^38,39,52,60^. The resulting genome structure population accurately reproduces Hi-C contact probabilities (Pearson’s *r*=0.98, genome-wide; 0.99 and 0.83 for cis and trans contacts, p=~0, average chromosome SCC^62^=0.87, **Extended Data Fig. 1a,b**, *Supplementary Information*). More than 99.91% of all contact constraints are fully satisfied and predicted contact frequencies show very small residuals (**Extended Data Fig. 1c**).

Model accuracy is assessed by predicting experimental data not used as input in the modeling process. First, models generated from a sparse Hi-C data set, with 50% entries randomly removed, predict the missing Hi-C contact frequencies with high accuracy (Pearson’s *r*=0.93 (cis) and 0.69 (trans) of missing data, p=~0, **Extended Data Fig. 1d,e**, *Methods*). Thus, our method is robust against missing data. Second, our models predict with good accuracy a host of orthogonal data from lamin B1 pA-DamID^40^, lamin B1 TSA-seq^14^, SON-TSA-seq^14^, and GPSeq^41^ experiments (Pearson’s *r*=0.80, 0.78, 0.87, and 0.80, respectively, **Table 1**, *Methods*). We will discuss these data in greater detail throughout this paper. Our models also confirm interior radial preferences of chromatin replicated in the earliest G1b phase (p=2.39e^−77^, Mann-Whitney-Wilcoxon test, two-sided) and predicted a gradual increase in average radial positions for chromatin replicated at progressively later times^63^ (**Extended Data Fig. 1f**). Our models also agree with 3D FISH experiments^19^, namely co-location frequencies of four inter-chromosomal pairs of loci (Pearson’s *r*= 0.99, p=0.014, **Extended Data Fig. 1g**), and distance distributions between three loci on chromosome 6 and relative differences in radial positions of these loci (**Extended Data Fig. 1h**). Finally, all our results are reproduced by technical replicates and converge even with smaller population sizes (*Methods*, *Supplementary Information*).

**Table 1.** Genome-wide correlations between experimental and predicted omics and imaging data. All p-values are ~0. Chromosome X is discarded from genome-wide correlation calculations in TSA-seq, DamID, and GPSeq comparisons.

|  | Pearson's *r* | Spearman's *r* |
| --- | --- | --- |
| SON TSA-seq<sup>14</sup> | 0.87 | 0.89 |
| Lamin B1 TSA-seq<sup>14</sup> | 0.78 | 0.81 |
| Lamin B1 pA-DamID<sup>40</sup> | 0.80 | 0.79 |
| GPSeq<sup>41</sup> | 0.80 | 0.79 |
| SAF<sup>8</sup> | 0.77 | 0.73 |
| LAF<sup>8</sup> | 0.64 | 0.58 |
| NAF<sup>8</sup> | 0.71 | 0.63 |
| Median trans A/B ratio<sup>8</sup> | 0.70 | 0.67 |

After establishing our models’ predictive value, we now determine the nuclear microenvironment of genomic regions by a variety of structural descriptors in each single cell model, such as their nuclear locations, distances to nuclear bodies and spatial compartmentalization (**Fig. 1a,b**). Our aim is to identify specialized nuclear microenvironments that distinguish chromatin in different functional states, and evaluate the roles of the nuclear architecture and its cell-to-cell variability in regulating transcription and replication.

**Fig. 1.**
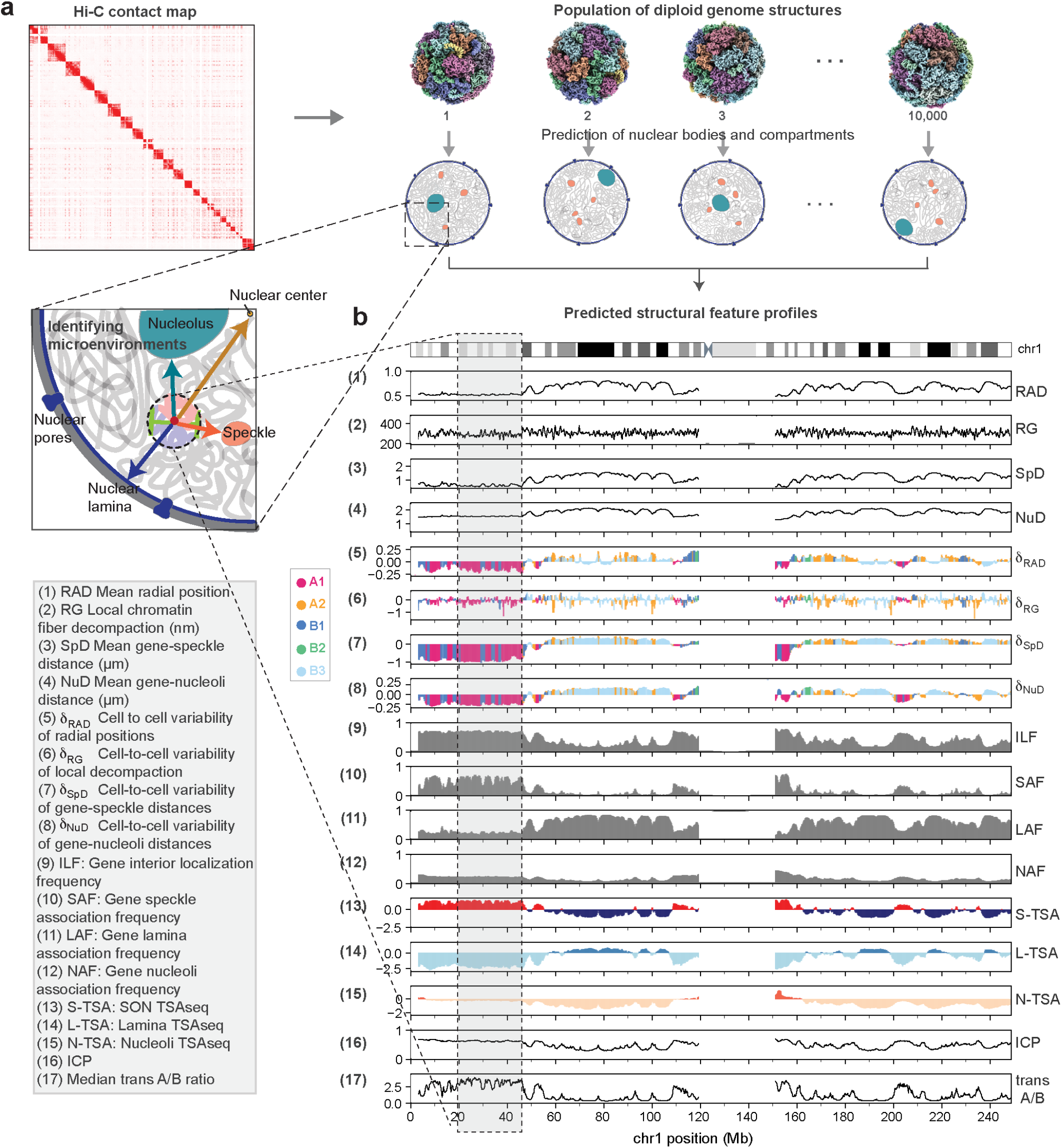
Microenvironment and structural features of genomic regions. **a**, Schematic depiction of our approach. A population of 10,000 genome structures is generated that is statistically consistent with the ensemble Hi-C data. Genome structures predict the locations of nuclear speckles, nucleoli and the lamina associated compartment, which serve as reference points to describe the global genome organization and define structural features. **b**, 17 structural features are calculated from the models that describe the nuclear microenvironment of each genomic region. Structure feature profiles for chromosome 1 are shown. Profiles for other chromosomes are shown in *Supplementary Information* (Fig. S5 – S25).

### Subnuclear positions and cell-to-cell heterogeneity vary by genomic loci

The nuclear positions of genes are of functional relevance. FISH experiments revealed for a number of genes, upon transcriptional activation, a statistical shift of their locations towards the nuclear center^64,65^. Due to their stochastic nature, nuclear positions of a locus can vary between individual cells. For instance, in our models some loci can be observed in an interior position in one structure of the population and close to the periphery in another (**Fig. 2a).** However, when averaged over the population of models it is evident that radial positions of genomic regions show preferred averaged locations, which substantially vary by genomic loci. Most evidently this is seen when plotting average radial positions along a chromosome. Radial profiles reveal pronounced minima and extended maxima, flanked by regions undergoing large radial transitions over relatively short sequence distances (**Fig. 2b,** upper panel). These minima overlap with regions of lowest lamin B1 DamID signals^66^ **(Extended Data Fig. 2a,b**). Our observations reproduce similar positional preferences detected in GPSeq experiments^41^ (Pearson’s *r*=0.80, p=~0, **Extended Data Fig. 2c,d**).

**Fig. 2.**
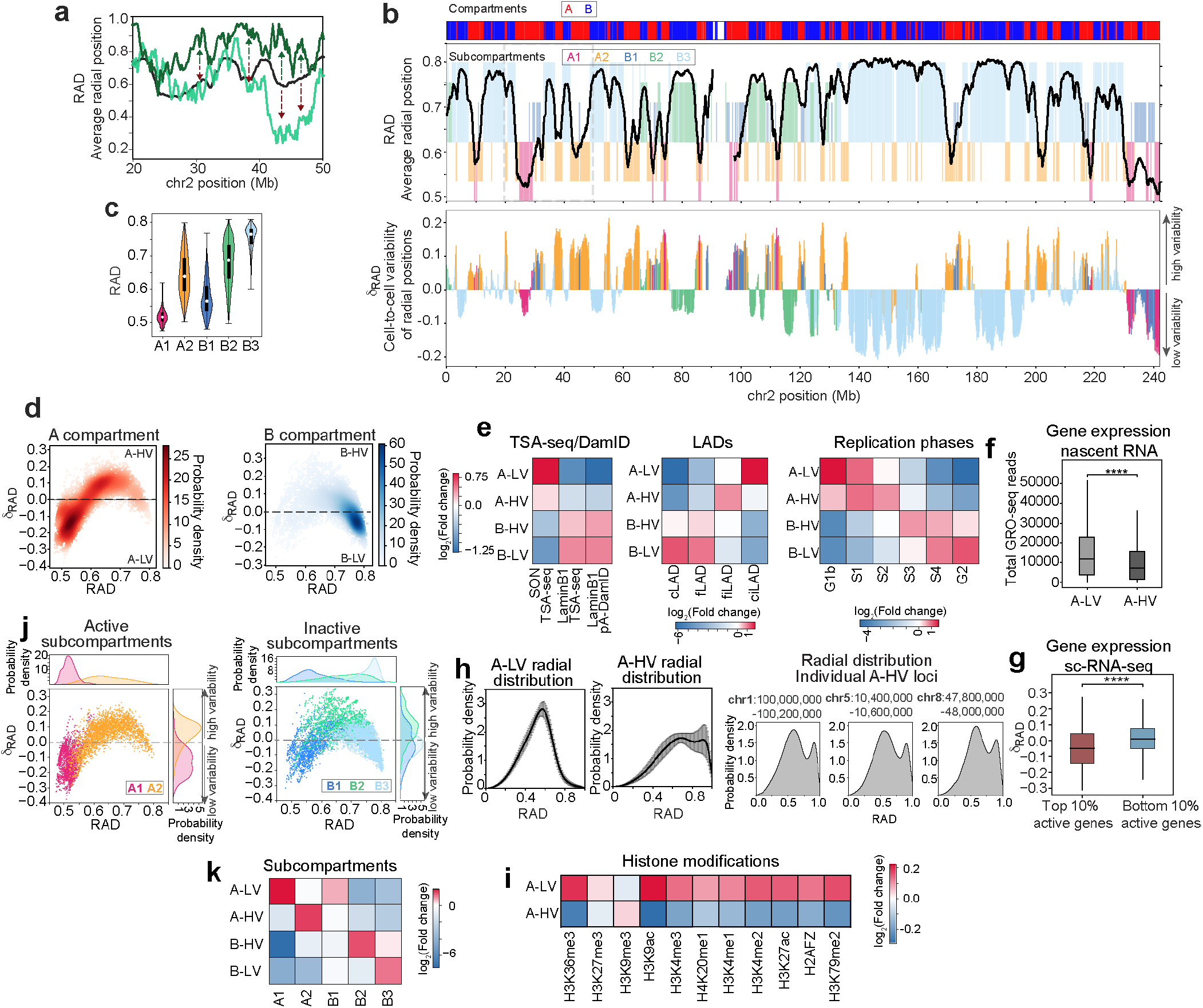
Radial chromatin positions and cell-to-cell variability. **a**, Radial position profiles for a 30 Mb region in chromosome 2. Black line shows the average radial position over the population of structures, and dark and light green lines show the radial positions in two different single structures. Arrows depict regions with high cell-to-cell variability. **b**, (upper panel) Average radial positions (RAD) of chromatin regions in chromosome 2. Background colors indicate subcompartment assignments, (lower panel) Cell-to-cell variability of radial positions (*δ_RAD_*) for each chromatin region in chromosome 2. Color-code for subcompartment annotations as in upper panel. **c**, Violin plots for the distributions of average radial positions for all chromatin regions in a subcompartment. White circles and black bars show the median value and the interquartile range (IQR: Q1 – Q3), respectively. **d**, Scatter density plots of *δ_RAD_* vs RAD for chromatin regions in A (left) and B (right) compartments. Dashed lines separate low (A-LV and B-LV) and high (A-HV and B-HV) levels of variability. **e**, Fold-change enrichment of SON TSA-seq^14^, Lamin B1 TSA-seq^14^ and pA-DamID^40^ signals (left), constitutive (cLAD) and facultative (fLAD) lamina associated regions (LAD) and constitutive and facultative inter-LADs^41,66^ (ciLAD and fiLAD, respectively) (middle), and replication phases^63^ (right) for chromatin regions with low and high cell-to-cell variability (*δ_RAD_*) in A and B compartment. **f**, Box plots of the nascent RNA expression levels (from GRO-seq experiments^67^) for chromatin regions in A compartment with low (A-LV) and high (A-HV) radial cell-to-cell variability (*δ_RAD_*) (Mann-Whitney-Wilcoxon test, two-sided). **g**, Box plots of radial cell-to-cell variability (*δ_RAD_*) distributions for chromatin regions with top 10% highest and bottom 10% lowest transcript numbers of actively transcribed genes according to scRNA-seq data^42^ (Mann-Whitney-Wilcoxon test, two-sided). **h**, Probability density distributions for the radial positions of A chromatin with low (*δ_RAD_* < Q1) and high (*δ_RAD_* > Q3) cell-to-cell variability (two left panels), and radial distributions of three representative A regions with high-cell to cell variability (three right panels). Black lines in the two left panels indicate the average distribution, and gray areas show the standard deviation from all regions within each group. **i**, Fold-change enrichment of histone marks in A-LV and A-HV groups. **j**, Scatter density plots of *δ_RAD_* vs RAD for chromatin regions in A1, A2 (left) and B1, B2, B3 (right) subcompartments. Top and right panels in each plot show the probability density distributions of RAD and *δ_RAD_* values for each subcompartment, respectively. **k**, Fold-change enrichment of subcompartments in A-LV, A-HV, B-LV, and B-HV groups.

Interestingly, regions undergoing large radial transitions often overlap with borders between the 5 primary Hi-C subcompartments identified by Rao *et al*.^34^ (**Fig. 2b**, upper panel, *Methods*), two transcriptional active (A1,A2) and three inactive subcompartments (B1,B2,B3). 76% of regions with high radial gradient are located at subcompartment borders. A1 chromatin, gene dense with relatively high GC content, shows the lowest, most interior average radial positions (**Fig. 2c**), with the highest probability at the most interior radial shell, with sharply decreasing probabilities otherwise, confirming previous observations^26,41^ (**Extended Data Fig. 2e**). B3 chromatin, mostly LADs^34^, show exterior average positions, with high probabilities at the outermost two shells and a relatively narrow distribution of average radial positions (**Fig. 2c, Extended Data Fig. 2e**). The B1 subcompartment, enriched in silencing H3K27me3, shares similar location preferences to A1—with highest probabilities in the interior radial shells (**Fig. 2c, Extended Data Fig. 2e**). However, the A2 subcompartment shows more evenly distributed average location probabilities across all nuclear shells (**Fig. 2c, Extended Data Fig. 2e**). Thus, A2 genomic loci do not share a common preference in their average positions. A similar behavior, with a relatively wide average radial distribution, is seen for all B2 regions, enriched in pericentromeric and nucleoli associated domains (NADs), with a slight increase in probabilities towards the outer half of nuclear shells (**Fig. 2c, Extended Data Fig. 2e**).

However, average positions alone are inferior measures of the dynamic positions of genes. They cannot convey if a region preferentially locates at the average location in most cells or if the region is rarely found at the average location, for instance when locations follow a multimodal or a skewed radial distribution. Our approach reveals the cell-to-cell variability of radial positions. To quantify stochastic variations of radial positions in the population of cells, we calculated *δ_RAD_*, the log-transformed fraction of observed and expected standard deviations of a genomic region’s radial position (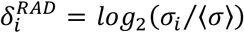, *Methods*). *δ_RAD_* differs distinctly between genomic loci (**Fig. 2b,** lower panel). High variability regions (*δ_RAD_* > 0) alternate, in sharp transition, with regions of low variability (*δ_RAD_* < 0)—transitions between high and low variability regions occur over relatively small sequence distances (**Fig. 2b,** lower panel). The smallest variabilities are observed for regions with the very lowest and highest average radial positions (**Fig. 2d**). Interestingly, intermediate average positions have almost exclusively high structural variability. Local peaks of high variability appear to coincide with local minima in the radial profiles at intermediate radial positions (dips with average radial positions ranging 0.55 – 0.70) (**Fig. 2b,** lower panel). This indicates that regions at local minima do not locate at intermediate radial positions in most cells, but greatly alternate between outer and inner locations between cells.

### Structural variability correlates with functional properties

Interestingly, structural variability of genomic regions is a strong indicator of their functional properties, for both chromatin in active A and inactive B compartments. We first divide chromatin of the active A compartment into a group with high (A-HV) (*δ_RAD_* > 0) and one group with low structural variability (A-LV) (*δ_RAD_* < 0) (**Fig. 2d**). A-LV, with low structural variability, is strongly enriched in signals from SON TSA-seq experiments, which specify short mean distances to nuclear speckles (**Fig. 2e,** left panel). A-LV regions are also strongly depleted in Lamin B1 pA-DamID signals and highly enriched for chromatin considered as constitutive inter LADs (ciLAD)—genomic regions never found to be associated with the lamina compartment across all studied cell types^66^ (**Fig. 2e,** middle panel). A-LV chromatin is dominantly replicated at the earliest G1b phase, which is distinctly different from A-HV regions, which are enriched for chromatin replicated at later S1 and S2 stages (**Fig. 2e,** right panel). A-LV chromatin show significantly higher transcriptional activity than A-HV regions (p=1.35e^−40^, Mann-Whitney-Wilcoxon test, two-sided, **Fig. 2f**). Overall, active genes with the lowest number of transcripts in single cell RNA-seq (scRNA-seq) experiment^42^ show significantly higher structural variability in their radial positions (*δ_RAD_*) (p=3.45e^−18^, Mann-Whitney-Wilcoxon test, two-sided, **Fig. 2g**,).

A-HV regions lack SON-TSA-seq enrichment and are enriched in facultative inter-LADS (fiLADs)— genomic regions that in some cell types can also be found to be lamina associated (**Fig. 2e**). Interestingly, A-HV regions with the largest structural variability often show a bimodal distribution in radial positions, an indication of two distinct favored locations—a nuclear interior and a peripheral location in a fraction of models (**Fig. 2h**). We hypothesize that these genes may exist in two functional states: active in the transcriptionally favorable interior, and silenced at the periphery. Indeed, A-HV regions are more enriched in H3K9me3 (related to heterochromatin) and depleted in H3K9ac (related to gene activation) than A-LV chromatin (**Fig. 2i**). This could point to a higher functional heterogeneity of these regions between cells (**Fig. 2i**).

B compartment chromatin also show functional differences between highly variable (B-HV) (*δ_RAD_* > 0) and low variable (B-LV) (*δ_RAD_* < 0) genomic regions. B-LV, with low structural variability, show higher enrichment in lamina associated features (i.e. lamin B1 pA-DamID and lamin B1 TSA-seq) than B-HV regions (**Fig. 2e**). Moreover, B-LV regions show strong enrichment for chromatin in constitutive LADs (cLADs)—regions that are always found as lamina associated in all studied cell types (**Fig. 2e**). In contrast, B-HV show higher enrichment in facultative LADs (fLAD), genomic regions that are lamina associated in most but not all cell types (**Fig. 2e**). Differences are also seen in replication timing. B-LV chromatin is replicated mostly at the very latest G2 phase (**Fig. 2e**), while B-HV regions are enriched in chromatin replicated at intermediate time points S3 and S4.

The structural variability of chromatin is also a distinguishing factor between Hi-C subcompartments. We found that A1 and A2 subcompartments, both active, can be distinguished by their structural variability alone (**Extended Data Fig. 2f**). A1 chromatin show overall the lowest, and A2 the highest *δ_RAD_* values (**Extended Data Fig. 2f**). 93% of all highly variable regions (*δ_RAD_* > 0) in the active compartment are A2 chromatin. Subsequently, A1 subcompartment chromatin are strongly enriched in A-LV genomic regions, while the A2 subcompartment are dominantly enriched in the A-HV regions (**Fig. 2k)**. Both subcompartments separate in two clusters, when considering average radial positions and radial variability (**Fig. 2j**).

B1, B2 and B3 subcompartments are also well distinguished by their structural variability and separate in three distinct clusters, when plotting average radial positions of chromatin against their radial variability (**Fig. 2j**). B2 chromatin is highly enriched in variable B-HV chromatin, while B3 chromatin is enriched in B-LV. B1 chromatin is structurally associated with A1 (**Fig. 2k**).

Interestingly, continuous genomic regions with similar trends in *δ_RAD_* are often part of the same subcompartment. These blocks with similar high variability (*δ_RAD_* > 0) alternate with blocks of low variability (*δ_RAD_* < 0). Transitions between high and low variability regions align remarkably well with the borders between subcompartments, most prominently between A2 and B3 subcompartments (**Fig. 2b,** lower panel).

### Subcompartments separate into spatial partitions

We now focus on the 3D compartmentalization of Hi-C subcompartments in single cell models. Chromosome folding permits functionally related chromatin to assemble into spatial compartments (**Fig. 3a**). When we calculated the single cell interaction networks (CINs) for chromatin in the same subcompartment, we saw a heterogeneous network organization with clusters of highly connected subgraphs intersected by low connectivity regions (**Fig. 3b**, *Methods*). Thus, subcompartment chromatin is divided spatially into a number of local partitions, which define nuclear territories with the highest concentration of chromatin in a given subcompartment. This organization is reminiscent to microphase fragmentation, instigated by the physical nature of the chromatin polymer preventing the segregation of each subcompartment into a single macrophase^32^. Spatial partitions are identified in single cells as highly connected subgraphs in the chromatin interaction network (**Fig. 3b**, *Methods*) and can be visualized in single genome structures by the occupied volume of the contained chromatin (**Fig. 3b,c**).

**Fig. 3.**
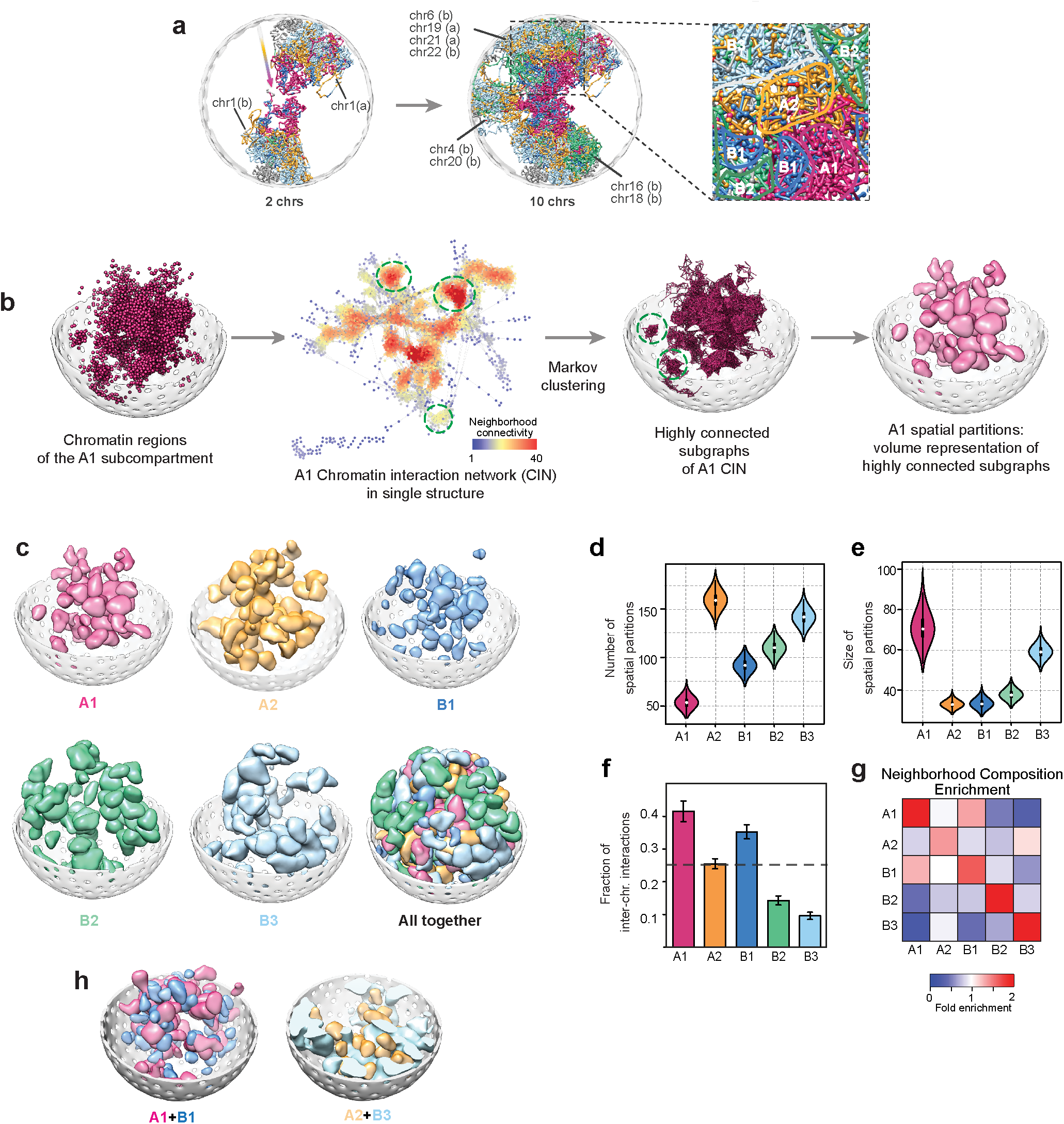
Spatial partitions of subcompartments. **a**, A representative genome structure showing chromosome folding patterns. Both images show the same structure with different numbers of chromosomes. Zoomed inset delineates regions that are primarily occupied by chromatin of the same subcompartment. Color-code indicates subcompartment annotations for each chromatin region (A1: pink, A2: yellow, B1: dark blue, B2: green, B3: light blue). **b**, Procedure to identify spatial partitions of subcompartments: A chromatin interaction network (CIN) is generated from all chromatin regions in a given subcompartment for each structure in the population. Each node in the CIN represents a single chromatin region connected by edges if the two regions are in physical contact in the 3D structure. Nodes are colored by their neighborhood connectivity (i.e. average contacts formed by their neighbor nodes) ranging from low (blue) to high (red). Highly connected subgraphs are then identified by Markov Clustering of CINs (*Methods*) and visualized in the 3D structure (some examples are shown in green dashed circles in 2D plot). The rightmost image illustrates the volume occupied by a spatial partition in a single genome structure. **c**, Spatial partitions of subcompartments, shown by their occupied volume in the 3D structures. For clarity only the 50 largest partitions (i.e. subgraphs with the largest numbers of nodes) are shown per subcompartment. **d**, Distributions of the number of subcompartment partitions per genome structure. **e**, Distributions of the average size (i.e. number of nodes) of subcompartment partitions. In d and e, white circles and black bars show the median value and the interquartile range (IQR: Q1 – Q3), respectively. **f**, Average fraction of inter-chromosomal edges in spatial partitions for each subcompartment. Error bars indicate standard deviations, and the gray dashed line is the average fraction of all partitions. **g,** Neighborhood enrichment of chromatin in each subcompartment, defined as the ratio of (observed/expected) subcompartment chromatin in the immediate neighborhood (within 500 nm) of each chromatin region (*Methods*). The strong diagonal shows that chromatin is preferentially surrounded by their own kind. **h,** A representative structure showing examples of colocalizations of A1-B1 and A2-B3 partitions in the 3D space.

Network structures differ between the subcompartments, and therefore, the size, number, and locations of spatial partitions also vary (**Fig. 3c,d,e, Extended Data Table 1**). For instance, A1 chromatin is fragmented into the fewest number (~50) but largest sized partitions of all subcompartments (**Fig. 3d,e, Extended Data Table 1**). These partitions contain the highest fraction of inter-chromosomal interactions (42%) (**Fig. 3f**). A2 networks are fragmented into larger numbers of smaller partitions, dominantly formed by intra-chromosomal interactions (75%) (**Extended Data Table 1, Fig. 3d,e,f**). While B1 networks also show high fragmentation into small partitions (**Fig. 3d,e**), they are formed by a larger fraction of inter-chromosomal interactions (35%) (**Fig. 3f**). B3 partitions are large and dominantly formed by intra-chromosomal interactions (90%) (**Fig. 3e,f**).

The larger partition sizes of A1, B2 and B3 chromatin lead to a more homogenous compartment organization—these chromatin are preferentially surrounded by their own kind (see high enrichment along the diagonal in **Fig. 3g**). Due to their smaller partition sizes, A2 and B1 chromatin show relatively high neighborhood enrichment with other chromatin (see off diagonal enrichment in **Fig. 3g**). A2 partitions are often closely associated with those of B3 chromatin, while B1 partitions associate with those of the A1 subcompartment^41^ (**Fig. 3g,h**).

We found that spatial partitions of active chromatin are regional territories of highest transcriptional activities. For instance, when we mapped nascent RNA expression from GRO-Seq experiments^67^ onto our genome structures, we found increasing transcriptional activities towards the centers of A1 partitions (**Fig. 4a**). A2 partitions show similar trends, although substantially lower signals (**Fig. 4a**). We also observe that highly expressed genes reside preferably in larger partitions, and expression levels at the centers of large A1 and A2 partitions are significantly higher than those of smaller ones (**Fig. 4b**). These observations indicate a functional relevance of spatial partitions, which we explore further in the following section.

**Fig. 4.**
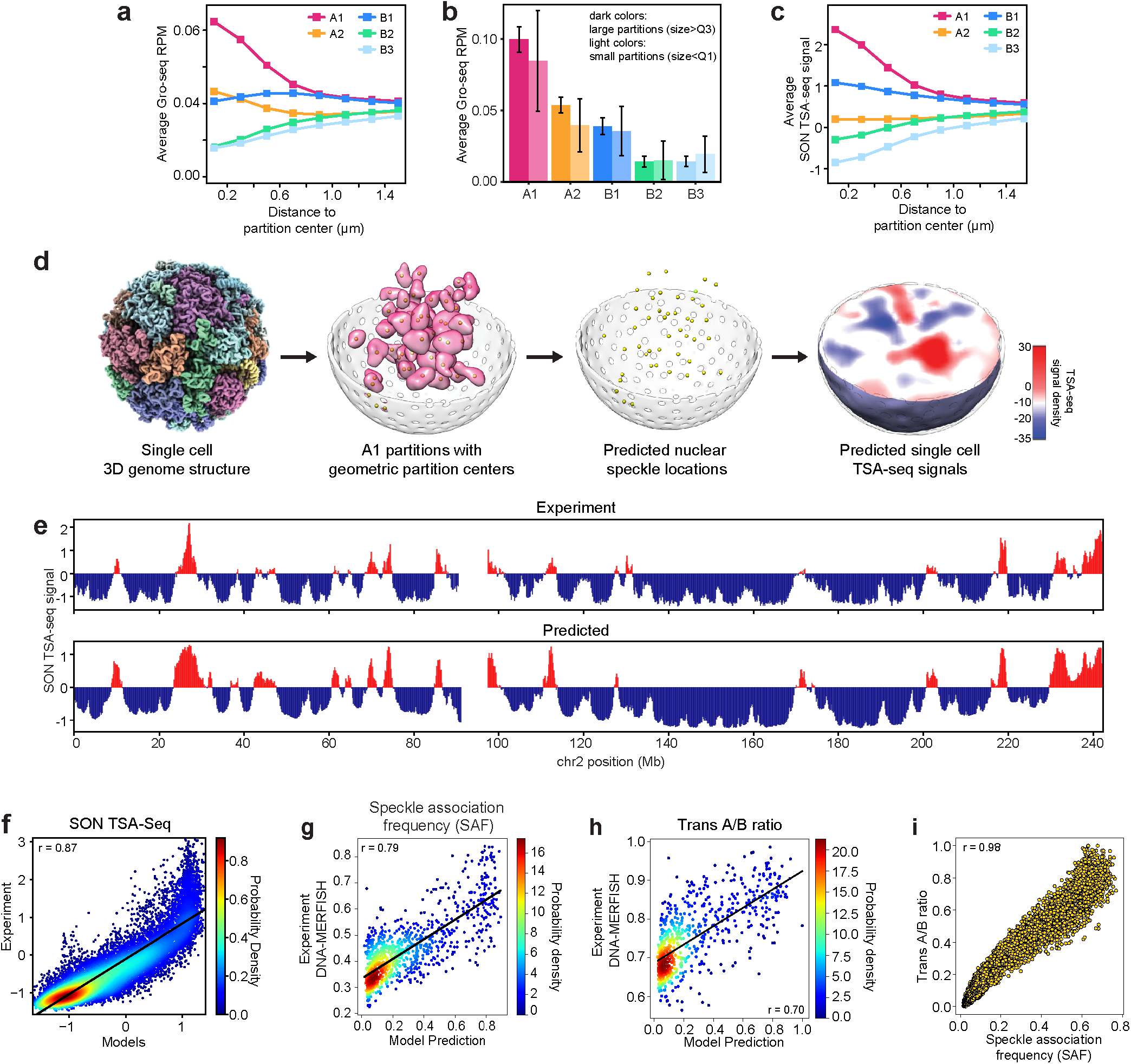
SON TSA-seq predictions using 3D structures. **a,** Average GRO-seq signal^67^ (RPM) of chromatin with respect to their 3D distances to subcompartment partition centers (*Methods*). **b**, Comparison of average GRO-seq signals^67^ for chromatin in large (size>Q3, dark colors) and small (size<Q1, light colors) spatial partitions for different subcompartments. Error bars are standard deviation. **c**, Average SON TSA-seq signals^14^ of chromatin with respect to their 3D distances to subcompartment partition centers (*Methods*). **d**, The procedure for SON TSA-seq signal prediction from 3D models: The geometric centers of identified A1 partitions in each single structure are used as point sources for the simulation of SON-TSA-produced tyramide free-radical diffusion^14^. SON TSA-seq signals are averaged over all structures (*Methods*). The rightmost image shows a cross section of the predicted TSA-seq signal density distribution in a genome structure. **e**, Comparison of the experimental and predicted SON TSA-seq profiles for chromosome 2 (Pearson’s *r*= 0.90, p=~0). **f**, Scatter density plot of the experimental vs predicted SON TSA-seq signals genome-wide (Pearson’s *r*=0.87, p=~0). **g**, Scatter density plot of the predicted speckle association frequencies (SAF) vs SAF determined with DNA-MERFISH experiments^8^ for 1,041 imaged loci (Pearson’s *r*= 0.87, p=~0). **h**, Scatter density plot of the median trans A/B ratios predicted in our models (*Methods*) vs from DNA-MERFISH experiment^8^ for 724 imaged loci that share the same compartment in GM12878 and IMR-90 cells (Pearson’s *r*=0.70, p=~0). **i**, Scatter plot of the predicted median trans A/B ratios vs SAF for each chromatin region in our models (Pearson’s *r*=0.98, p=~0).

### Predicting locations of nuclear speckles and speckle associated structural features

We now infer locations of nuclear bodies in single cell models. When we mapped TSA-seq data to chromatin in our structures, we noticed that TSA-seq signals are strongest—thus have smallest mean speckle distances—for chromatin located towards the central regions of A1 partitions (**Fig. 4c**). A2 partitions are devoid of TSA-seq signals (**Fig. 4c**). These observations suggest that A1 partition centers could represent locations of nuclear speckles in individual cell models.

To test this assumption, we simulated the experimental TSA-seq process by using A1 partition centers as approximate speckle locations (**Fig. 4d**). SON TSA-seq relies on a gradient of diffusible tyramide free-radicals, instigated at speckle locations, to measure distance-dependent labelling of DNA^14^. The steady state concentration of tyramide free-radicals at any given chromatin location can then be modeled in single cells with an exponential decay function using the spatial distances to all predicted speckle locations in a model^14^ (**Fig. 4d**, *Methods*). The simulated SON TSA-seq data, averaged over the population of cells, agrees remarkably well with experiment (Pearson’s *r*=0.87 p=~0), capturing both, peak sizes and signal distributions (**Fig. 4e,f**). For instance, the TSA-seq profile of chromosome 2 is reproduced with high correlation (Pearson’s *r=0*. 90, p=~0) across the entire chromosome profile, even though it contains only few A1 regions (6.4%) (**Fig. 4e**). Chromatin with different predicted TSA-seq signals show characteristic enrichment of histone modifications, identical to those observed in the experiment^14^. This confirms high prediction accuracy across all ranges of TSA-seq values (**Extended Data Fig. 3a**). Moreover, predicted speckle locations confirm the proposed correlation between mean speckle distances of chromatin and its experimental TSA-seq signal (**Extended Data Fig. 3b**).

Interestingly, prediction accuracy is dramatically reduced when simulations are performed on isolated chromosomes (i.e., extracted from the genome model), even when identical chromosome conformations are used (e.g. Pearson’s *r* for chromosome 17 drops from 0.82 to 0.52) (genome-wide Pearson’s *r*=0.73, p=~0, **Extended Data Fig. 3c, Extended Data Table 2**). This points to a substantial contribution of trans interactions. When we assume A2 partition centers as speckle locations, simulations fail entirely (Pearson’s *r*=0.18, p=9.4×10^−98^ **Extended Data Fig. 3d**). Also, random chromosome territories (Pearson’s *r*=0.60, p=~0) or simulations based on A1 sequence positions, rather than 3D structures, do not produce accurate TSA-seq profiles (Pearson’s *r*=0.35, respectively, p=~0) (**Extended Data Fig. 3c, Extended Data Table 2**).

To generalize our approach to other cell types, we devised a prediction method that does not rely on subcompartment annotations. We found that spatial partitions of chromatin with the 10% lowest average radial positions predict speckle locations within 500nm to those derived from A1 partitions in 99% of structures (78% of chromatin with 10% lowest radial positions are part of A1.). Subsequently, the simulated SON TSA-seq data is almost identical, with excellent accuracy (Pearson’s *r*=0.86, p=~0) (**Extended Data Fig. 3d, Extended Data Table 2**). Thus, our approach predicts speckle locations and SON-TSA-seq signals using only Hi-C data. This is important, because subcompartment annotations are available only to a limited number of cell types.

With predicted speckle locations as reference points, we now calculate speckle-associated features for genomic regions, namely the (i) mean distance to the closest speckle (SpD), (ii) cell-to-cell variability of the specke distances (*δ_SpD_*), and (iii) the speckle association frequency (SAF), as the fraction of models in which a genomic region is in close proximity with a speckle (**Fig. 1**, *Methods*). The predicted SAF agrees remarkably well with those in a recent DNA-MERFISH microscopy study^8^ (Pearson’s *r*=0.79, p=8.4×10^−223^, **Fig. 4g**, *Methods*).

We also calculated for each genomic region the trans A/B density ratio, defined as the ratio of A and B compartment chromatin forming inter-chromosomal interactions with the target loci^8^. Trans A/B ratios calculated from our models show good agreement with DNA-MERFISH experiments (Pearson’s *r*=0.70, p=7.6×10^−109^, **Fig. 4h**). Our models also confirm the correlation between a gene’s SAF and its trans A/B ratio from experiment (Pearson’s *r*=0.98, p=~0, **Fig. 4i**)^8^.

#### Defining lamina associated structure features

The lamin compartment at the nuclear periphery is an important component of the nuclear architecture. Lamin B1 associated chromatin features are calculated with the nuclear envelope as reference point (*Methods*). Simulated lamin B1 TSA-seq data (Pearson’s *r*=0.78, p=~0, **Extended Data Fig. 4a, Table 1**) and lamin B1 DamID data (Pearson’s *r*=0.80, p=~0, **Extended Data Fig. 4b, Table 1**) are in good agreement with experiment^14^, thus validating correct mean distances and contact frequencies of genomic regions with lamin B1 at the nuclear periphery. Simulated lamina association frequencies (LAF) show also high correlation with those from DNA-MERFISH imaging^8^ (Pearson’s *r*=0.64, p=~3.6×10^−119^, **Extended Data Fig. 4c**), although the correlation is lower than for SAF predictions, likely due to shape differences between flat IMR-90 and spherical GM12878 cell nuclei. Predicted LAF values are inversely correlated with a gene’s trans A/B ratios and SAF, confirming previous observations from DNA-MERFISH imaging^8^ (**Extended Data Fig. 4d**).

#### Defining nucleolus associated structure features

Nucleoli are major organizing factors in genome structure. To calculate nucleoli related features, we identify approximate nucleoli locations from spatial partitions formed by chromatin known to be nucleolus organizing regions (NOR) (short arms of chromosomes 13,14,15, 21 and 22), and nucleolus associated domains (NADs)^68^ (*Methods*). The centers of these chromatin partitions in single cell models then serve as reference points to calculate for each genomic region the mean nucleoli distance (NuD), the cell-to-cell variability of the NuD (*δ_NuD_*), nucleoli association frequencies (NAF) and nucleoli-TSA-seq data (**Fig. 1**). The NAF calculated in our models shows good agreement with NAF extracted from DNA-MERFISH imaging (Pearson’s *r*=0.71, p=1.2×10^−152^, **Extended Data Fig. 4e**, *Methods*).

#### Defining structural features of the local chromatin fiber

Finally, we also calculate features of the local chromatin fiber structure. The volume occupied by a chromatin region relates to its local compaction and is estimated, for each chromatin region, by the radius of gyration (RG) of a continuous 1Mb window centered at the target locus (**Extended Data Fig. 5a**, upper panel, **Fig. 1**, *Methods*). Average RG profiles show pronounced maxima at locations of TAD boundaries, while minima show domain-like compaction (**Extended Data Fig. 5a,b,c**). RG profiles in single cells show distinct maxima and minima, which can vary between cells (**Extended Data Fig. 5a, lower panel**). The probability for observing a peak is at maximum at TAD border locations, while randomly selected regions show a flat probability distribution (**Extended Data Fig. 4d)**. About 20% of structures show a RG peak (i.e., domain border) at the exact TAD border location (50% show a RG peak within the immediate vicinity). These TAD border frequencies in single cell structures agree with recent observations in oligoSTORM superresolution imaging^5^.

### The spatial microenvironment of a gene mirrors its functional state

Overall, we calculated a total of 17 structural features from single cell genome models (**Fig. 1**). Together, these features define the nuclear microenvironment of each genomic region. The advantage of our approach is that we can determine these features simultaneously in each single cell model, which allows us to analyze correlations between them and assess the role of the nuclear microenvironment to explain functional differences between chromatin, in particular for gene transcription, DNA replication and chromatin compartmentalization.

#### Gene transcription

We now investigate the role of the nuclear microenvironment in gene transcription. We first compare the stochastic variability of gene-speckle distances across single cell models with the variability of single cell gene expression from scRNA-seq experiments^42^. For each chromosome, we plot a heatmap representing all gene-speckle distances in the cell population (**Fig. 5a** top panel). Each column contains the cumulatively ranked distances between a genomic region and the nearest predicted speckle in all models of the population (**Fig. 5a** top panel). Likewise, we plot a heatmap representing the number of gene transcripts found in all single cells of a population. Each column contains the cumulatively ranked transcript numbers of genes from scRNA-seq data^42^ for each genomic region in all cells of the population (**Fig. 5b,** top panel). The two heatmaps show striking similarities. We then compared the gene transcription frequency (TRF), defined as the fraction of cells a transcript is detected in scRNA-seq experiments^42^ with the SAF predicted from the models (**Fig. 5a,b**, lower panels). The TRF and SAF profiles are remarkably similar and show highly significant correlation (**Fig. 5c**, left panel, Spearman’s *r*=0.51, p=~0). Genes with transcripts in a large fraction of cells are located close to speckles in a large fraction of models. This is an interesting finding. It links the nascent transcript frequency of a gene to its local nuclear environment. Thus it is possible that the local nuclear environment of a gene defines its transcription potential if gene expression is initiated. We also validated our finding with transcription frequencies measured in a recent RNA-MERFISH microscopy study for 1,137 genes^8^ (**Fig. 5c,** right panel). Here as well, we observe the identical highly significant correlation between TRF and SAF (Spearman’s *r*=0.51, p=1.6×10^−64^). Interestingly, a gene’s interior location frequency (ILF) (i.e., the fraction of cells a genomic region is located in the interior of the nucleus) shows substantially smaller correlation with the TRF than the SAF, for both from scRNA-e and RNA-MERFISH experiments (Spearman’s *r*=0.42, p=~0 (scRNA-seq) and 0.45, p=4.1×10^−50^ (RNA-MERFISH)) (**Fig. 5c**).

**Fig. 5.**
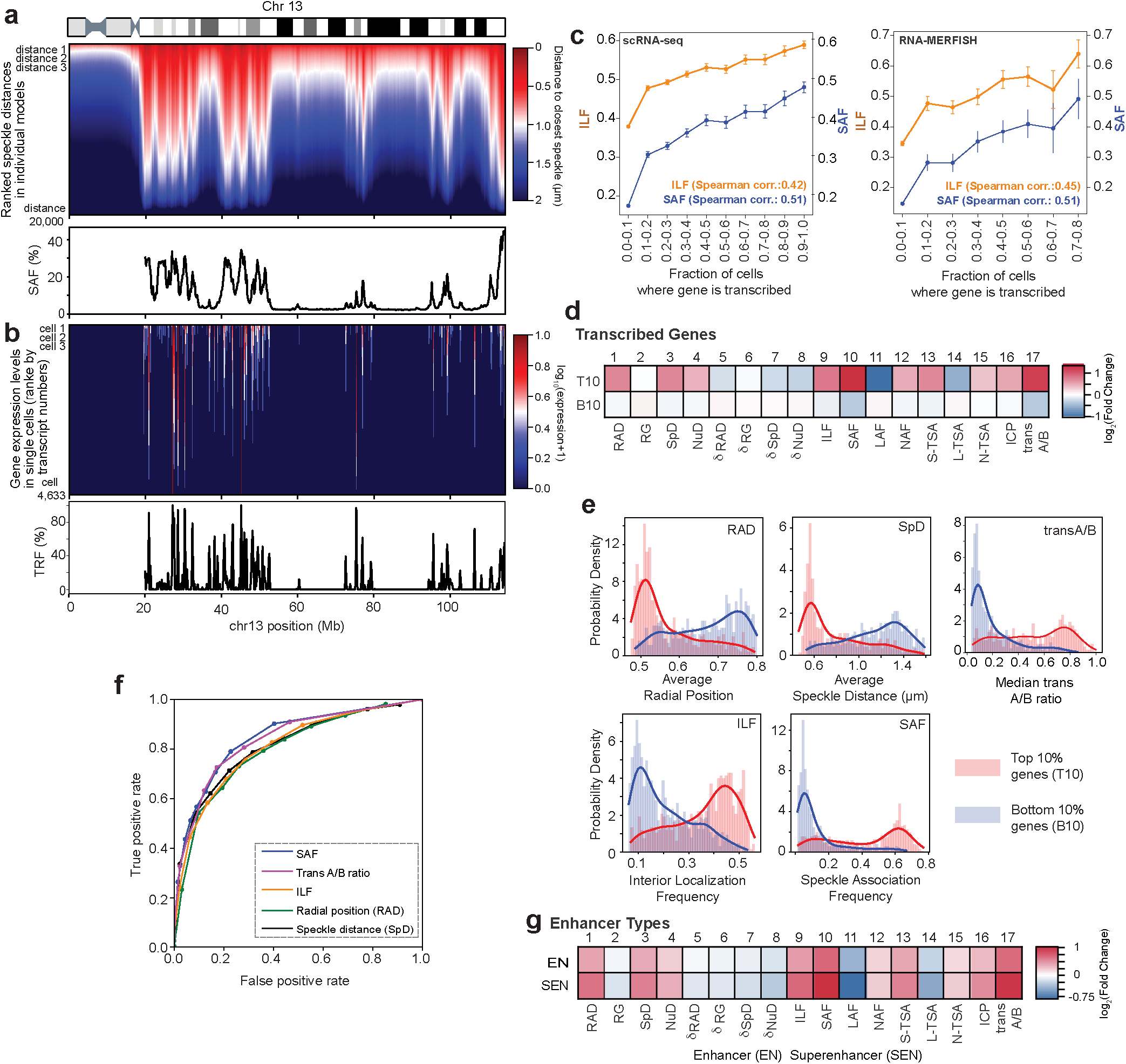
Relationship between 3D chromatin structure and transcriptional activity. **a**, (Top panel) Heatmap of gene speckle distances in chromosome 13 in 10,000 structures. The column shows for a given gene the gene-speckle distances in all 10,000 structures of the population. In each column, gene-speckle distances are sorted in ascending order from top-to-bottom, with short distances (dark red) to large distances (dark blue). (Bottom panel) Speckle association frequencies (SAF) for each chromatin region in chromosome 13. **b**, (Top panel) Heatmap of single cell mRNA counts of genes in chromosome 13 in all 4,633 G1 cells measured by single cell RNA-seq (scRNA-seq) experiment^42^. For a given gene, each column shows the observed mRNA transcript count in each cell of the population of cells. In each column, mRNA transcript counts are sorted in descending order from top-to-bottom, with high counts (dark red) to zero counts (dark blue). (Bottom panel) Transcription frequency (TRF) for each gene in chromosome 13 from scRNA-seq data^42^ (*Methods*). **c**, Interior localization frequency (ILF) and SAF values for genes with different TRF ranges from scRNA-seq^42^ (left) and nascent RNA-MERFISH imaging^8^ (right). Error bars show standard deviations of ILF and SAF values in each TRF range. **d**, Fold-change enrichment for each of the 17 structural features for chromatin with top 10% highest (T10) and bottom 10% lowest (B10) transcript numbers of actively transcribed genes according to scRNA-seq data^42^ (*Methods*). **e**, Distributions of several structural features for T10 and B10 regions. **f**, Receiver Operator Characteristic (ROC) curves for radial positions, speckle distances, ILF, SAF, and trans A/B ratios to distinguish T10 and B10 regions (Area under the curve values are 0.65, 0,72, 0.81, 0.85, 0.84, respectively). **g**, Fold-change enrichment for each of the 17 structural features for enhancer (EN) and superenhancer (SEN) chromatin regions.

Next, we study if genes with the 10% largest number of RNA transcripts (T10) are distinguished in their nuclear environment from genes with the 10% lowest number of transcripts (B10). T10 genes show strong enrichment for several structural features, particularly those related to nuclear speckles and trans A/B ratio (**Fig. 5d)**. T10 genes are also depleted in structural variability relative to nuclear bodies (*δ_RAD_*, *δ_SpD_* and *δ_NuD_*). Therefore T10 genes show a strong preference for a specific microenvironment and show relatively high homogeneity between cells (**Fig. 5d**). Lowly expressed B10 genes do not show any preferential positioning relative to nuclear bodies, and show more variable nuclear locations than T10 genes (**Fig. 5d**). They also show, significant depletion in SAF and trans A/B ratio, and thus, are overall clearly distinguished in their microenvironment from T10 genes.

We further assess, which single feature (among SpD, ILF, SAF, RAD, and trans A/B) is most discriminative in separating the two gene sets. Distributions of feature values are quite different between the two gene sets (**Fig. 5e**). However, SAF and the highly correlated trans A/B ratio outperform all other features in distinguishing T10 from B10 genes, as shown by the receiver operating characteristics (ROC) curves (**Fig. 5f**). Thus, speckle associated features, and SAF in particular, are more predictive of gene expression than the average radial position (RAD) (area under ROC curve: SAF: 0.85, RAD: 0.65) or other features derived from radial positions. This finding could indicate that the general preference of highly expressed genes at the nuclear interior may be an indirect consequence of favored associations with nuclear speckles, which themselves show stochastic preferences towards the nuclear interior^14,69^.

Next, we divide all genes into two groups: those dominanty controlled by enhancers (EN) and those controlled by superenhancers (SEN) (*Methods*). Overall, genomic regions with EN and SEN show similar enrichment patterns to those of T10 genes (**Fig. 5g**). However, genomic regions with SEN show substantially higher fold enrichments and depletions than EN genes, revealing stronger preferences in their nuclear microenvironment, particularly for higher SAF, interior positions, trans A/B ratio, ICP and depletion of LAF values (**Fig. 5g**). Notably, for both EN and SEN features related to cell-to-cell variability are depleted, revealing a higher homogeneity of their structural features in the cell population.

#### The organizing role of nuclear speckles

Our approach also allows a detailed analysis of experimental SON-TSA-seq data^14^. For instance, chromatin divided by their experimental SON-TSA-seq signals^14^ into ten groups show distinct structural enrichment patterns, which gradually change with increasing SON TSA-seq values (**Fig. 6a**). Chromatin in the first (d1,d2) and last (d9, d10) deciles show the highest fold enrichments and thus, the most stable microenvironment. These regions show the lowest structural variability in radial positions (*δ_RAD_*) and have the smallest (d1-d3) and largest (d8-d10) mean speckle distances (SpD), respectively (**Fig. 6b**). Thus these regions show high levels of structural homogeneity between cells in the population. In contrast, chromatin in deciles d4-d7 are structurally less defined (**Fig. 6a**), are highly variable in their nuclear positions (*δ_RAD_*) (**Fig. 6b**), and particularly chromatin in decile 6, show no preferred associations towards nuclear bodies (**Fig. 6a**).

**Fig. 6.**
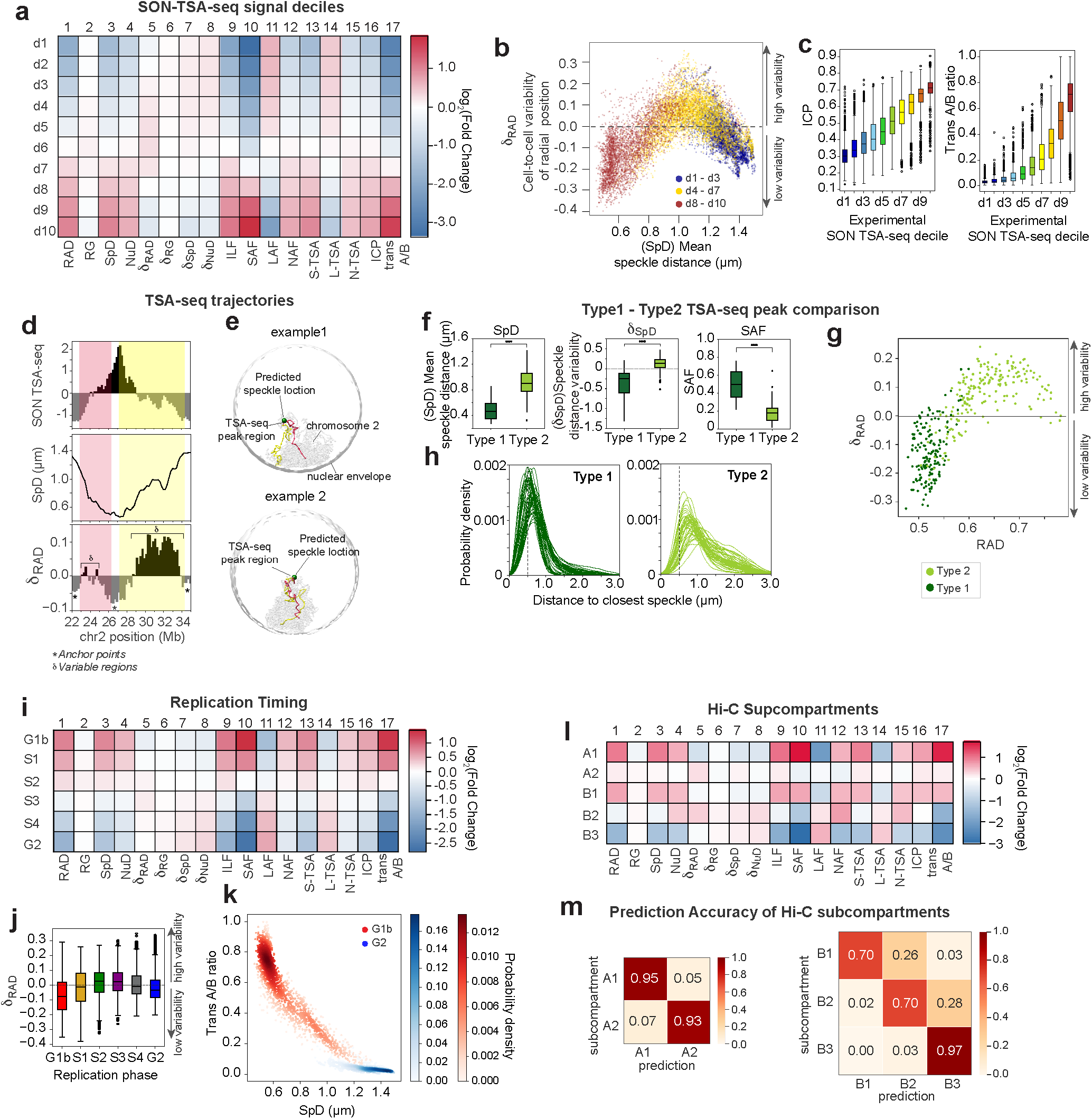
Structural features of microenvironments. **a,** Fold-change enrichment of 17 structural features for chromatin regions in experimental SON-TSA-seq decile groups^14^. **b**, Scatter plot of the radial cell-to-cell variabilities (*δ_RAD_*) vs mean speckle distances (SpD) of chromatin in experimental SON-TSA-seq decile groups^14^ (d1 – d3: blue, d4 – d7: yellow, d8 – d10: red). **c**, Distributions of inter-chromosomal contact probabilities (ICP, left) and trans A/B ratios (right) for chromatin in each experimental SON-TSA-seq decile group^14^. **d**, Experimental SON TSA-seq signals (top), SpD (middle) and *δ_RAD_* (bottom) profiles for a ~11 Mb region of chromosome 2 showing a so-called TSA-seq trajectory transition in the TSA-seq profile. Stars in the lower panel indicate anchor regions with low structural variability, while regions marked with δ indicate high variability regions. (Valley-to-peak: red region, peak-to-valley: yellow region). **e**, Two representative structures showing folding patterns of the chromatin fiber for the ~11 Mb TSA-seq trajectory as in d, together with the nuclear envelope, the closest predicted speckle location (green), and the rest of chromosome 2 (gray). The chromatin fiber is color coded in red and yellow to represent corresponding regions shown in d. **f**, Distributions of SpD (left), speckle distance variabilities (*δ_SpD_*, middle), and SAF (right) for regions where Type I and Type II TSA-seq peaks^14^ are located (Mann-Whitney-Wilcoxon test, two-sided). **g**, Scatter plot of *δ_RAD_* vs RAD for Type I (dark green) and Type II (light green) chromatin regions. **h**, Distributions of gene-speckle distances for randomly selected 50 individual Type I loci (left) and Type II loci (right) in the population. Gray dashed line indicates the 0.5 μm distance level. **i**, Fold-change enrichment of 17 structural features for regions at different replication phases^63^. **j**, Distributions of *δ_RAD_* values for chromatin in each replication phase^63^. **k**, Scatter density plot of trans A/B ratios vs SpD for chromatin in G1b (red) and G2 (blue) replication phase^63^. **l**, Fold-change enrichment of 17 structural features for chromatin regions in each subcompartment. **m**, Confusion matrices for the prediction of A1 and A2 (left) and B1, B2 and B3 (right) subcompartments using K-means clustering based on structural features (*Methods*).

We also observe a high correlation between the inter-chromosomal contact probability (ICP) (i.e. the fraction of a region’s trans vs. all interactions) and the experimental SON TSA-seq signals, and thus mean speckle distances (Pearson’s *r*=0.76 p=~0, **Fig. 6c,** left). Notably, the trans A/B ratio of a genomic region is also positively correlated with its SON-TSA-seq value (**Fig. 6c,** right). These observations imply that nuclear speckles act as major hubs to facilitate inter-chromosomal interactions of transcriptionally active genomic regions, confirming similar findings reported earlier^14,24^.

Our models also allow a structural interpretation of reported TSA-seq trajectories, steep transitions in TSA-seq profiles between low and high peaks (**Fig. 6d,** top panel)^14^. In our models, TSA-seq trajectories coincide with steep transitions in the average speckle distances and average radial positions (**Fig 6d**, middle panel). In a fraction of models, these chromosome regions fold from anchor regions at the outer nuclear periphery towards the nuclear interior, where the TSA-seq peak region often associates with a nuclear speckle and forms the apex of a chromosomal loop, which then traces back to the nuclear periphery (**Fig. 6e**). These anchor regions at the periphery and the loop apex show low structural variability (*δ_RAD_*), while loop regions in between show higher variability (**Fig. 6d**, lower panel). These findings are in agreement with similar observations in FISH experiments by the Belmont laboratory^14^.

SON TSA-seq experiments identified two types of transcription “hot zones”: Type I and Type II regions with high and intermediate SON TSA-seq signal peaks, respectively^14^. Our models confirm the expectation that Type I regions have significantly smaller mean speckle distances than Type II (Mann-Whitney-Wilcoxon two-sided test, p=1.3×10^−51^, **Fig. 6f**, left panel). However, TSA-seq data is inconclusive as to whether Type II regions persistently reside at intermediate speckle distances or localize at speckles in a small fraction of cells and far from them in others^14,70^. Our models uncover the latter case. The vast majority of Type II regions show a significantly higher variability in radial positions (**Fig. 6g**) and speckle distances (p=1.94e^−43^, Mann-Whitney-Wilcoxon test, two-sided, **Fig. 6f,** middle panel), and associate with speckles in a smaller fraction of cells (average SAF < ~17%) (**Fig. 6f**, right panel). Thus most Type II regions do not reside stably at intermediate speckle distances and show a wide and, in many cases, bimodal speckle distance distribution (**Fig. 6h**). In contrast, Type I regions show stable radial positions at close speckle distances (**Fig. 6h**), resulting in high SAF of about 50% (p=2.82e^−53^, Mann-Whitney-Wilcoxon test, two-sided, **Fig. 6f**, right panel).

#### The role of microenvironment in replication timing

Replication timing of chromatin^63^ is echoed in distinct structural features (**Fig. 6i)**. Chromatin replicated at early time points (G1b, S1) are most enriched in SAF, trans A/B, and have lowest structural variability (**Fig. 6i,j**). Chromatin replicating in the intermediate S2 and S3 phases show the highest cell-to-cell variability in nuclear positions and show no preferential association with speckles, nucleoli or the lamin compartment (**Fig. 6i,j)**. Late replicating chromatin (S4 and G2 phase) are depleted in interior locations and speckle associated features and strongly enriched in lamina associated features (**Fig. 6i)**. Overall, SAF and trans A/B ratio are more discriminative with higher fold changes than features related to the radial positions, RAD and ILF (**Fig. 6i**). For instance, trans A/B ratio and mean speckle distances clearly separate early from late replicating chromatin (**Fig. 6k**).

#### Chromatin compartmentalization

Hi-C subcompartments also show distinct enrichment patterns, thus represent distinct physical microenvironments (**Fig. 6l, Extended Data Fig. 6,** *Methods*). Chromatin in the A1 subcompartment is well separated from A2 chromatin across all studied structural features (**Fig. 6l**). A1 chromatin show strong enrichement patterns, with strong preferences in their microenvironment and small structure variations, thus a high level of uniformity. It is particularly enriched in speckle associated features and trans A/B. A2 chromatin has relatively weak enrichment patterns and high cell-to-cell variability in radial locations, speckle distances and overall wide distributions of feature values within their class (**Fig. 6l, Extended Data Fig. 6**). Thus, A2 chromatin shows no clear location preferences with respect to any studied nuclear bodies. B3 chromatin shows strong anti-correlated enrichment patterns with A1 across all structural features. B2 chromatin is well separated in its enrichment patterns from B3 chromatin, mainly due to enriched nucleoli and depleted lamin-based features and its high variability in nuclear locations, possibly due to prevalent locations of nucleoli at both central and peripheral regions (**Fig. 6l, Extended Data Fig. 6)**. B1 chromatin shares similar enrichment patterns with A1 chromatin, although with smaller fold enrichment (**Fig. 6l**). Thus, B1 genes containing polycomb silenced chromatin would be in a position of highest transcriptional potency, if activated.

The structural differences between subcompartments are so pronounced that we are able to predict Hi-C subcompartments from structural features alone without explicit considerations of chromatin interactions. Unsupervised K-means clustering based on structural feature vectors of compartment A chromatin predicts A1 and A2 subcompartment annotations with 94% accuracy, while chromatin in inactive subcompartments were predicted with an accuracy of 84%. These results are comparable in accuracy to supervised methods using Hi-C contact frequencies^71^ (**Fig. 6m,** *Methods*). This is an important finding, because subcompartment predictions by the Rao *et al*.^34^ approach for cell types other than GM12878 cells have failed so far. Our approach provides an alternative way of detecting subcompartment annotations, while also providing underlying structural interpretations.

## Discussion

We introduced an approach to determine a population of single cell 3D genome structures from ensemble Hi-C data. Our method is unique as it predicts a host of structural features in single cell models, and provides information about the structural microenvironment of genomic regions in single cells. This information is not available from Hi-C data without structural modeling. Therefore, our method considerably expands the scope of Hi-C data analysis and is widely applicable to other cell types and tissues for which Hi-C data is available.

The models and the derived structural features are a powerful resource to unravel the relationship between genome structure and function. We found that cell-to-cell heterogeneity of structures vary by genomic loci and is a strong indicator of functional properties. Structurally stable chromatin in the A compartment are dominantly associated to nuclear speckles, and show relatively high speckle association frequencies, high trans A/B ratio and overall lowest average radial positions. These regions contain highly transcribed genes, are enriched in superenhancers, SON TSA-seq signals and are replicated at the earliest time points. Moreover, these genomic regions compartmentalize in relatively large spatial partitions, formed by a high fraction of inter-chromosomal interactions. Chromatin of the A1 subcompartment is enriched in this category.

In contrast, active chromatin with high structural variability are characterized by the lack of preferences in nuclear locations. In a fraction of cells, these regions can be located in a silencing environment at the nuclear periphery, while in others, can be located towards the transcriptionally favorable interior. These genes show relatively low transcript frequencies, low inter-chromosomal contact probabilities with low trans A/B ratios and intermediate replication timing (phases S2, S3). In TSA-seq experiments, most of these regions were identified as Type II peaks, with intermediate TSA-seq values. We also noticed that these regions compartmentalize into relatively small spatial partitions (i.e., microphases), dominated by intra-chromosomal interactions. Chromatin of the A2 subcompartment is enriched in this category.

It is possible that the high structural variability of these regions could be linked to functional heterogeneity between cells. Several observations point to this conclusion. For instance, although being transcriptionally active, these regions are enriched in silencing H3K9me3 and depleted in activating H3K9ac histone modifications in comparison to active regions with low structural variability. Moreover, gene transcripts for these genomic regions are found in a substantially smaller fraction of cells and show overall lower transcriptional activity.

Interestingly, structural heterogeneity is also an indicator to distinguish nucleoli and lamina associated chromatin in the B compartment. Genomic regions with low structural variability are dominantly associated to the lamina compartment and constitutive LADs and enriched in the B3 subcompartment. Genomic regions with high structural variability are associated with nucleoli and pericentromeric heterochromatin and are enriched in the B2 subcompartment.

Our results suggest that nuclear speckles, together with the lamina compartment, are a major organizing factor in genome structure. Chromatin with low structural variability are associated with either nuclear speckles or constitutive LADs. Speckle locations are not randomly distributed in the nucleus, but are more likely to be excluded from the nuclear periphery^14,69^. Therefore, LADs, at the periphery, and speckles, towards the interior, provide structural anchor points. We hypothesize that A-LV and B-LV regions associated with these anchors act in a similar way to recently reported fixed points in the nuclear organization of mouse embryonic stem cells^9^. For instance, genomic regions with high SAF or LAF have low structural variability and act as anchor points for radial trajectories detected in SON TSA-seq experiments.

Moreover, the inter-chromosomal contact probability (ICP) and trans A/B ratio are highly correlated with mean speckle distances. Therefore, nuclear speckles appear to be the sole hub of inter-chromosomal interactions of active chromatin regions. These findings agree with similar observations from SPRITE experiments^24^. The relatively high fraction of trans interactions for speckle associated chromatin could provide an explanation for the preferential locations of speckles toward the nucler interior. We find that the probability of inter-chromosomal interactions increases towards the central regions of the nucleus (**Fig. 7a**). If some speckles will associate with regions from at least two chromosomes, they are more likely located at the interior. Over time, dynamic interactions with multiple chromosomes may restrain their locations towards the interior (**Fig. 7b**). These cooperative effects could bias the global speckle distributions towards the nuclear interior.

**Fig. 7.**
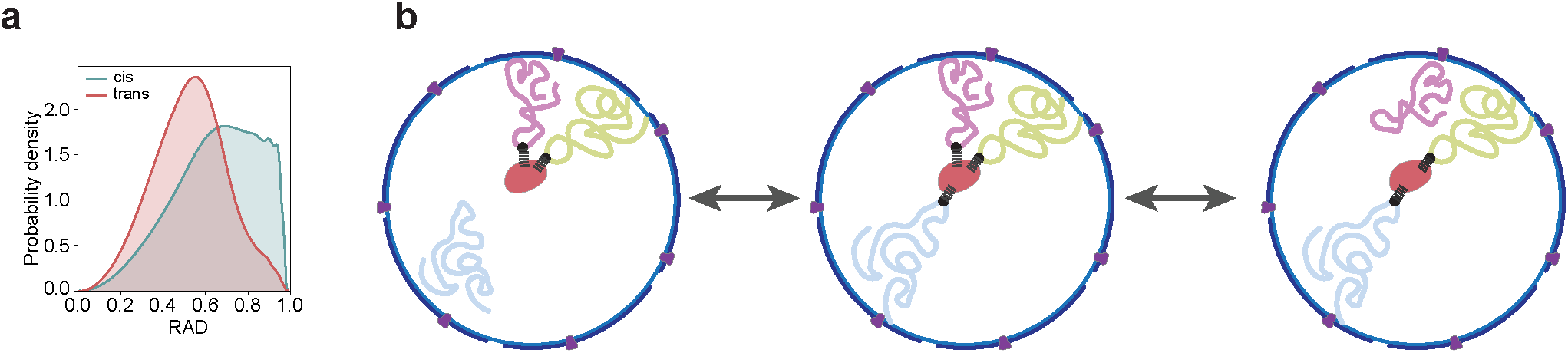
Inter-chromosomal interactions and speckle locations. **a,** Distributions of radial positions where cis and trans interactions occur in the models **b,** Scheme for the proposed effect of inter-chromosomal interactions on speckle (red) locations in the nucleus.

We demonstrate that the structural microenvironment of a genomic region is directly linked to its functional potential in gene transcription and replication. Chromatin with highest and lowest transcriptional activity are distinguished by their structural features. In particular, the frequency of close speckle associations (SAF) shows the highest correlation with the gene transcription frequency^9,72–74^. The known interior preference of highly activated genes could therefore be a consequence of preferential positions to nuclear speckles, which in turn have a stochastic preference towards the nuclear interior, confirming previous observations from TSA-seq experiments^14^. Chromatin replicated at the earliest time are also distinguished in their structural features from those replicating at the latest stages. Moreover, our observations confirm that Hi-C subcompartments define physically distinct chromatin environments, some of which (such as A1) linked to associations with nuclear bodies.

In summary, our method produces a large number of structural descriptors highly relevant for a better understanding of genome structure function relationships. These features can be calculated from Hi-C data alone, and thus are applicable to many different cell types for a comparative analysis of genome structures. In the future, we plan to incorporate nuclear shapes from imaging into the modeling process to include a more realistic representation of the shape and size of nuclear bodies and the nucleus.

## Acknowledgements

This work was supported by the National Institutes of Health (grant U54DK107981 and UM1HG011593 to F.A) as part of the 4D Nucleome Initiative, and an NSF CAREER grant (1150287 to F.A.).

## Extended Data Figures and Tables

**Extended Data Fig. 1.**
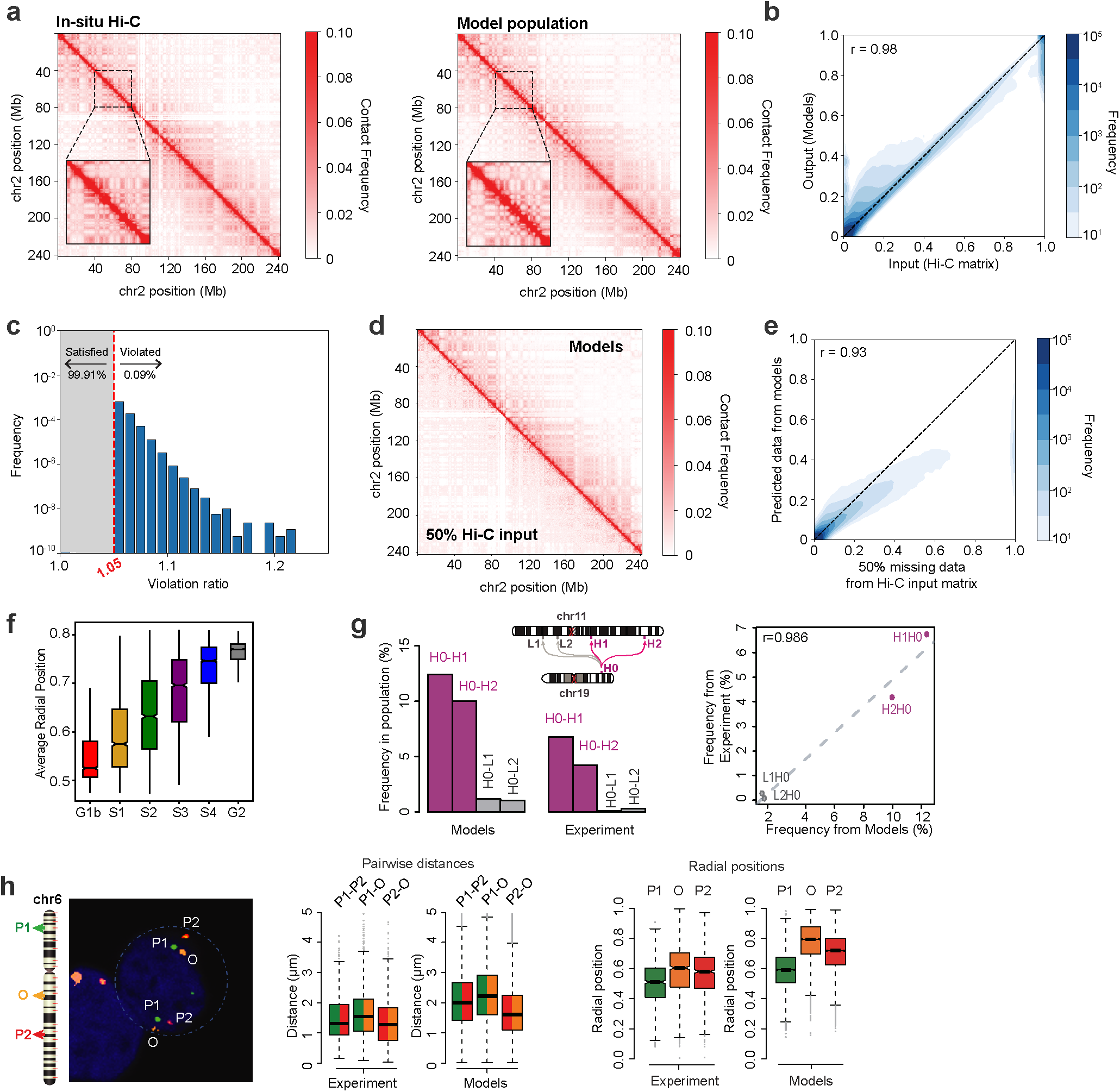
3D chromatin structure modeling and assessment. **a**, The Hi-C contact probability matrix (left) and the contact probability matrix calculated from the structure population (right) for chromosome 2. Zoomed-in heatmaps show the matrix between sequence position 40 – 80 Mb. **b**, Density scatter plot comparing the contact probabilities from Hi-C data and structure population (Pearson’s *r*=0.98, p=~0). **c**, Histograms of restraint-violation ratio from the structure population (*Methods*). A violation ratio less than 1.05 is considered satisfied and is not displayed in the histograms (99.9% of restraints fall in this category). **d**, The contact probability matrix for chromosome 2 showing the 50% randomly chosen dataset used as input (lower triangle) vs. the matrix generated from the structure population (upper triangle). **e**, Density plot comparing the contact probabilities that are generated from Hi-C data and missing in the input and their predicted contact probabilities calculated from the structure population (Pearson’s *r*=0.93, p=~0). **f**, Average radial positions of chromatin in different replication phases^63^. **g**, Comparison of the inter-chromosomal loci co-localization frequencies between the observed occurrence in FISH experiments^19^ and in the structure population (left), and scatter plot showing the co-localization frequencies from FISH experiments and the structure population (right). **h**, A FISH image with three different probes at far-separating loci on chromosome 6 (left), the comparison of pair-wise distances of these loci in experiment and models (middle), and the comparison of their relative radial positions in experiment and models (right).

**Extended Data Fig. 2.**
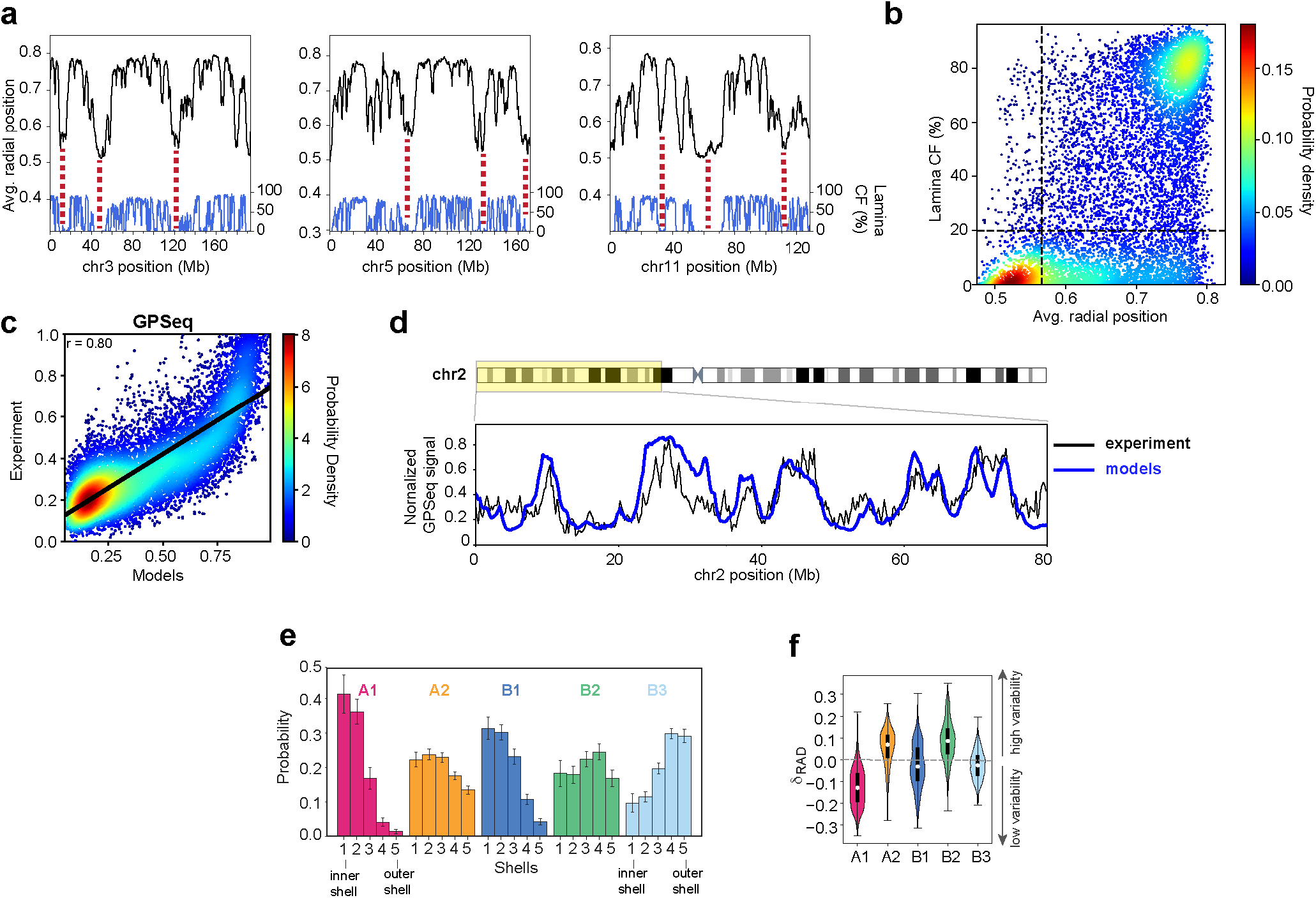
Assessment of radial positions. **a**, Average radial position profiles in chromosomes 3 (left), 5 (middle), and 11 (right). Also shown in blue are lamina CF from single cell lamin DamID experiments^66^. Valleys in the average radial position plots match well with low lamina CF regions (red dashed lines). **b**, Density scatter plot of average radial positions of chromatin regions from the structure population against the lamina contact frequencies from single cell lamin DamID experiments in haploid KBM7 cell type (CF; DamID data from^66^). 93% of chromatin regions with the 25% lowest average radial positions show either no detectable or only occasional contact with lamina (CF < 20%). Vertical and horizontal black dashed lines show the 25^th^ percentile average radial position and the 20% CF values, respectively. **c**, Scatter plot showing the comparison between experimental and predicted GPSeq scores^41^ (Pearson’s *r*=0.80, p=~0) **d**, Comparison of experimental and predicted GPSeq^41^ profiles for the 0 – 80 Mb region in chromosome 2. **e**, Probabilities for chromatin region of a given subcompartment to be located in any of five concentric shells, each containing the same total amount of chromatin (*Methods*).Shell 1 is the most interior shell. Error bars show standard deviation. **f**, Violin plots for distributions of cell-to-cell variabilities of radial positions (*δ_RAD_*) for chromatin regions in different subcompartments. Dashed line separates low and high levels of variability.

**Extended Data Fig. 3.**
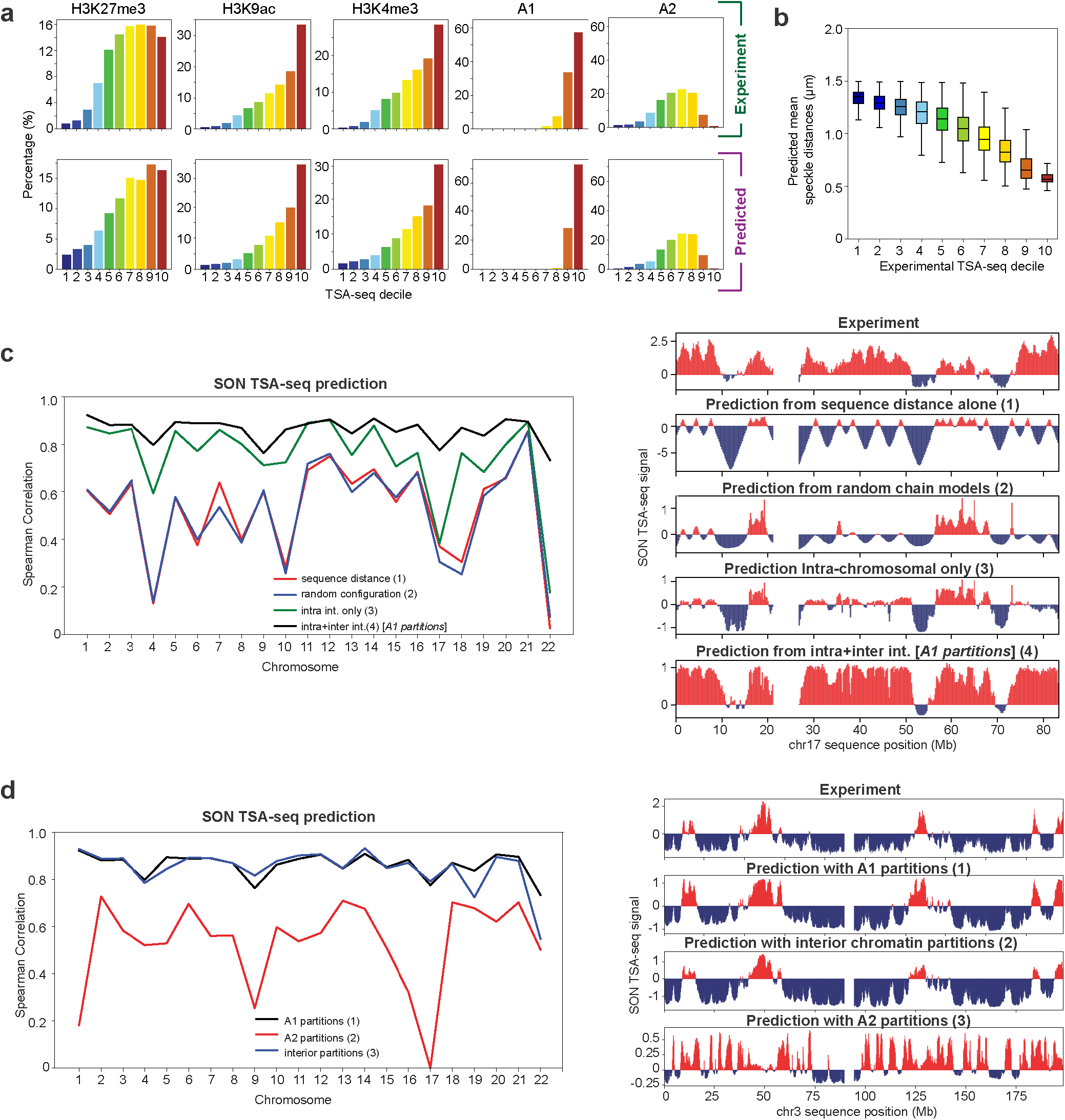
SON TSA-seq predictions using 3D models. **a**, Fraction of mapped histone modifications peaks from ChIP-seq experiments as well as number of A1/A2 chromatin regions in chromatin divided into decile groups based on their experimental^14^ (top) and predicted SON TSA-seq signals (*Methods*). **b**, Distributions of predicted mean distances to closest speckles (A1 partition centers) for chromatin regions in each experimental SON TSA-seq decile^14^. **c**, Spearman correlations between the experimental^14^ and predicted SON TSA-seq signals for each chromosome using different prediction methods (left, *Methods*); predictions using sequence distances to A1 clusters in sequence (red), 3D distances to A1 partitions in random chain chromosome territories (blue), 3D distances to A1 regions in the same chromosome only (green), and 3D distances to A1 partition centers using both intra-and inter-chromosomal relationships (black). Corresponding TSA-seq profiles of chromosome 17 for predicted and experimental data (Spearman correlations: 0.37, 0.30, 0.38, 0.78, respectively, right). **d**, Spearman correlations between experimental^14^ and predicted SON TSA-seq signals for each chromosome using different partitions as predicted speckle locations (left, *Methods*); predictions using A1 spatial partition centers (black), A2 spatial partition centers (red), and spatial partitions from chromatin with 10% lowest average radial positions in the population (blue). Corresponding TSA-seq profiles of chromosome 3 for predicted and experimental data (Spearman correlations: 0.88, 0.89, 0.58, respectively, right).

**Extended Data Fig. 4.**
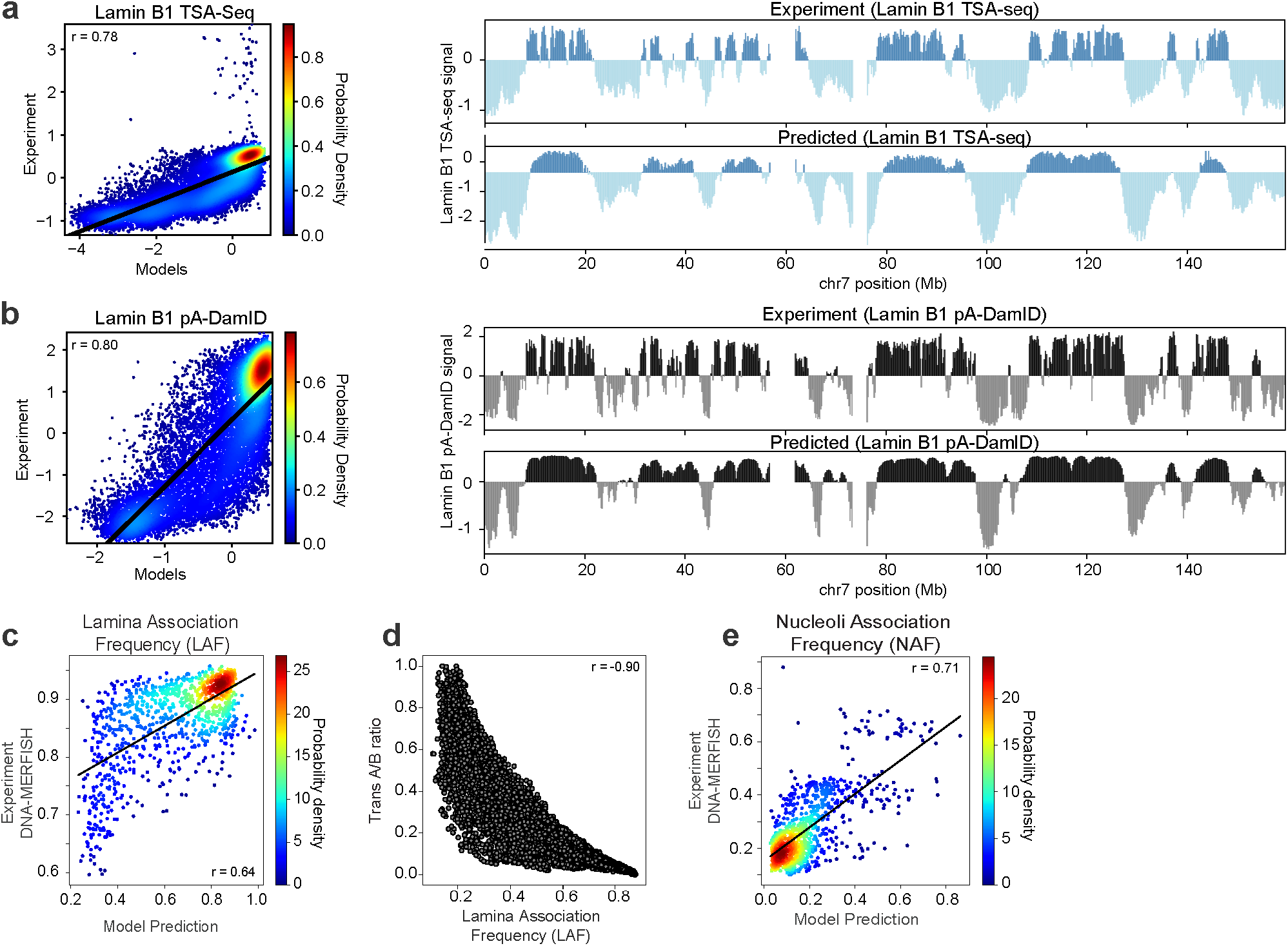
Predictions of lamin and nucleolus associated features. **a**, Scatter plot showing the comparison between experimental and predicted Lamin B1 TSA-seq signals^14^ (left, Pearson’s *r*=0.78, p=~0), and chromosome 7 profiles of experimental and predicted Lamin B1 TSA-seq signals^14^ (right). **b**, Scatter plot showing the comparison between experimental and predicted Lamin B1 pA-DamID signals^40^ (left, Pearson’s *r*=0.80, p=~0) and chromosome 7 profiles of experimental and predicted Lamin B1 pA-DamID signals^40^ (right). **c**, Comparison of predicted lamina association frequencies (LAF) in our models (*Methods*) with LAF determined from DNA-MERFISH experiments^8^ for 1,041 imaged loci (Pearson’s *r*=0.64, p=~0). **d**, Scatter plot of predicted median trans A/B ratios as functions of predicted LAF for each chromatin region in our models (Pearson’s *r*=-0.90, p=~0). **e**, Comparison of predicted nucleoli association frequencies (NAF) in our models (*Methods*) with NAF determined from DNA-MERFISH experiments^8^ for 1,041 imaged loci (Pearson’s *r*=0.71, p=~0).

**Extended Data Fig. 5.**
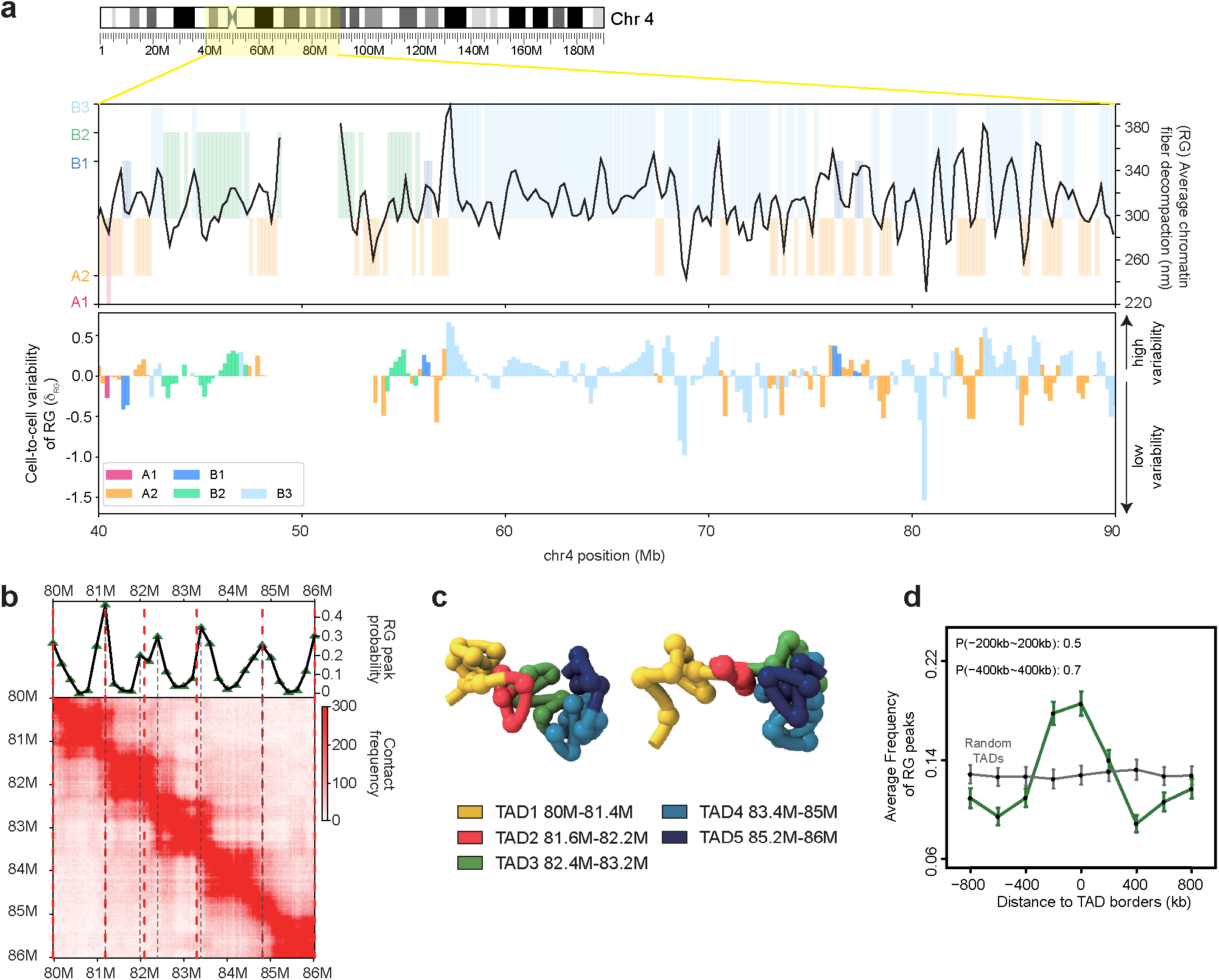
Chromatin compaction and TAD borders. **a,** Average radius of gyration (RG, i.e. local decompaction) profile for chromatin in the 40 – 90 Mb region of chromosome 4. The background is color coded by the subcompartment annotations of chromatin (top). Cell-to-cell variability of RG values (*δ_RG_*) in the structure population for the same chromatin regions. Negative values indicate regions with low RG variability (bottom). Bars are color coded by the subcompartment annotations of the corresponding chromatin regions. **b,** RG peak frequencies (i.e., the fraction of models showing a RG maximum at a given position) for a 6-Mb region in chromosome 4 (80–86Mb) (top), and Hi-C contact frequency heat map for the same region showing TAD borders identified by TopDom^75^ (bottom). Regions with RG peak frequency maxima are shown with gray dashed lines, and either overlap or are very close to TAD borders identified by TopDom (red dashed lines). **c,** Two representative structures showing chromatin folding patterns for chromatin regions in b. TAD identities are shown by color code. **d**, Averaged RG peak frequencies for loci at TopDom TAD borders (green) compared to randomly selected loci (gray). In around 50% of structures, there is a RG peak in the immediate neighboring region of a TAD border (±200kb). In ~70% of structures there is a RG peak within a ±400kb range of a TAD border. Standard errors calculated from all TAD borders are shown with error bars.

**Extended Data Fig. 6.**
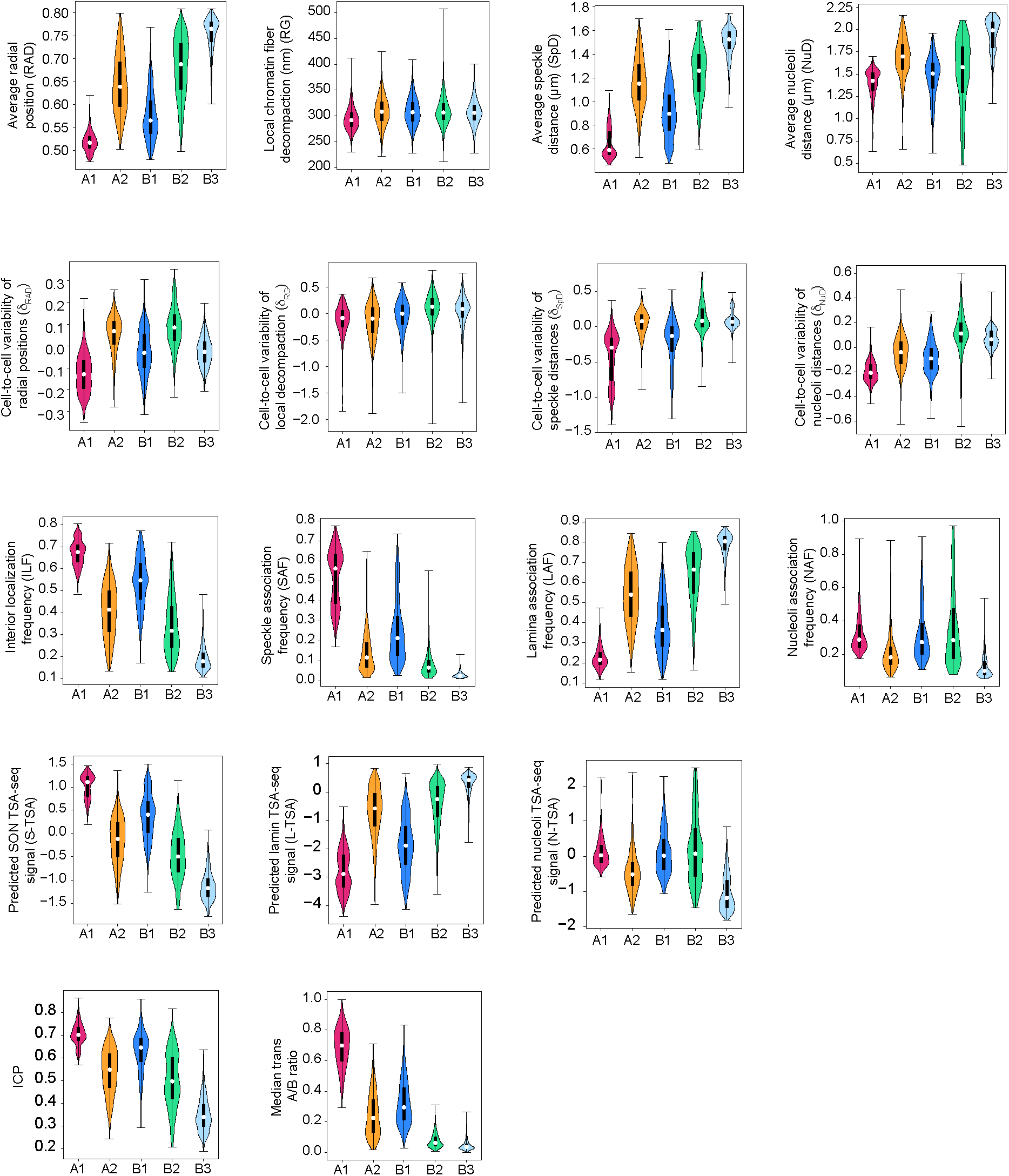
Structural features of chromatin in different subcompartments. Violin plots for the distributions of 17 structural features calculated from the structure population for chromatin in different subcompartments. White circles and black bars in the violins show the median value and the interquartile range (IQR: Q1 – Q3), respectively.

**Extended Data Table 1.** Properties of subcompartment interaction networks and spatial partitions. Population averages of features for chromatin interaction networks (CIN) and spatial partitions of chromatin in different subcompartments (*Methods*).

|  | Subcompartments |  |  |  |  | Compartments |  |
| --- | --- | --- | --- | --- | --- | --- | --- |
| CIN/Partition Features | A1 | A2 | B1 | B2 | B3 | A-LV | A-HV |
| Average neighborhood connectivity in CINs | 25.92 | 12.15 | 12.88 | 13.58 | 15.63 | 20.48 | 8.46 |
| Maximal cliques enrichment in CINs | 5.99 | 1.64 | 2.22 | 2.16 | 1.70 | 4.29 | 1.39 |
| Average radial position of partitions | 0.57 | 0.70 | 0.60 | 0.71 | 0.77 | 0.56 | 0.69 |
| Average size of partitions (number of 200 kb regions) | 71.00 | 32.90 | 33.28 | 37.73 | 59.01 | 54.60 | 17.68 |
| Average number of partitions in each structure | 53.86 | 159.23 | 91.63 | 109.79 | 141.85 | 52.73 | 155.94 |
| Average fraction of trans edges in partitions (%) | 41.52 | 25.49 | 35.26 | 14.29 | 9.57 | 34.14 | 15.58 |

**Extended Data Table 2.** Genome-wide correlations between experimental and predicted SON TSA-seq data using different approaches. All p-values are ~0. Chromosome X is discarded from genome-wide correlation calculations.

|  | **Pearson's *r*** | **Spearman's *r*** |
| --- | --- | --- |
| SON TSA-seq<sup>14</sup> predictions with A1 partitions | 0.87 | 0.89 |
| SON TSA-seq<sup>14</sup> predictions with interior partitions | 0.86 | 0.88 |
| SON TSA-seq<sup>14</sup> predictions with A2 partitions | 0.18 | 0.38 |
| SON TSA-seq<sup>14</sup> predictions with A1 sequence distances | 0.35 | 0.64 |
| SON TSA-seq<sup>14</sup> predictions from random configurations (only-intra) | 0.60 | 0.58 |
| SON TSA-seq<sup>14</sup> predictions from folded chromosomes<br>(only-intra) | 0.73 | 0.79 |

## Methods

### Population Based 3D Structural Modeling

#### ▪ General description

Our goal is to generate a population of 10,000 diploid genome structures, so that the accumulated chromatin contacts across the entire population are statistically consistent with the contact probability matrix **A** = (*A_ij_*)_*N*×*N*_ derived from Hi-C experiments^19,39^. To achieve this goal, we utilize population-based modeling, our previously described probabilistic framework to de-multiplex the ensemble Hi-C data into a large population of individual genome structures of diploid genomes statistically consistent with all contact frequencies in the ensemble Hi-C data^38,39,61^.

The structure optimization is formulated as a maximum likelihood estimation problem solved by an iterative optimization algorithm with a series of optimization strategies for efficient and scalable model estimation^38,39,52^. Briefly, given a contact probability matrix **A** = (*A_ij_*)_*N*×*N*_, we aim to reconstruct all 3D structures ***X*** = { ***X*_2_**… ***X_M_***} in the population of *M* models, each containing 2*N* genomic regions for the diploid genome (at 200 kb base-pair resolution), and 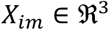, *i*. = 1.2*N* as coordinates of all diploid genomic regions in model *M*. We introduce a latent indicator variable **W** = (***W_ijm_***)_2*N*×2*N*×*M*_ for complementing missing information (i.e. missing phasing and ambiguity due to genome diploidy). **W** is a binary-valued 3^rd^-order tensor specifying the contacts of homologous genomic regions in each individual structure of the population, such that 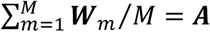. We can jointly approximate the structure population **X** and the contact tensor **W** by maximizing the log-likelihood of the probability:

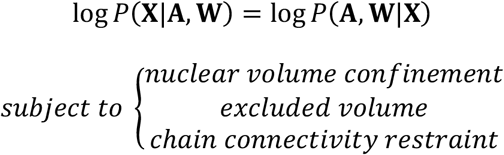

where

i. Nuclear volume constraint: All chromatin spheres are constrained to the nuclear volume with radius 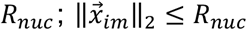, where 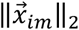 is the distance of the region *i*. from the nuclear center in structure *m*.
ii. Excluded volume constraint: This constraint prevents overlap between two regions represented by spheres, defined by their excluded volume radii 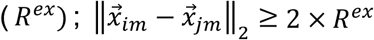.
iii. Polymer chain constraint: Distances between two consecutive 200-kb spheres within the same chromosomes are constrained to their contact distance to ensure chromosomal chain integrity; 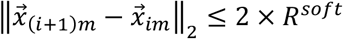, where *R^soft^* = 2 × *R^ex^*.

Our modeling pipeline uses a step-wise iterative process, in which the optimization hardness is gradually increased by adding contacts with decreasing contact probabilities in the input matrix. The iterative optimization procedure involves two steps, each optimizing local approximations of the likelihood function: (1) Assignment step (A-step): Given the estimated structures **X** at step k, estimate **W**; and (2) Modeling step (M-step): Given the estimated **W**, generate model population X at step k+1 that maximizes likelihood to observe W. Structures in the M-step are calculated using a combination of optimization approaches, including simulated annealing molecular dynamics simulations.

Moreover, during each optimization cycle we also use iterative refinement steps, a methodological innovation for effective reassignment of restraints during the optimization process, which allows genome structure generation at higher resolution and improved accuracy in comparison to our previous approach^38,39^ (see Iterative refinement method in *Supplementary Information*).

After 11 iterations, our method converges and the genome-wide contact probabilities from the structure population agree remarkably well with those from the Hi-C experiment.

#### ▪ Genome representation

The nucleus is modeled as a sphere with 5 μm radius (*R_nuc_*)^39^. Chromosomes are represented by a chromatin chain model at 200-kb base-pair resolution. Each 200-kb chromatin region, in the diploid genome, is modeled as a sphere, defined by an excluded volume radius (*R^ex^* = 118 nm). *R^ex^* is estimated from the sequence length, the nuclear volume and the genome occupancy (40%), as described in ref.^39^. The full diploid genome is represented with a total of 30,332 spheres.

##### Random starting configurations

Optimizations are initiated with random chromosome configurations. Chromatin regions are randomly placed in a bounding sphere proportional to its chromosome territory size and randomly placed within the nucleus.

#### ▪ Comparison between contact frequency maps from Hi-C experiment and model population

To quantify the agreement between Hi-C experiment and model population, we perform the following analyses:

1. Comparison between input and output Hi-C maps are evaluated by Pearson and stratum adjusted (SCC)^62^ correlation coefficients (Table S1).
2. Restraint violation ratios. On average about 175,304 contact restraints are imposed in each of the 10,000 structures. The restraint score of each contact restraint *i* is calculated as: 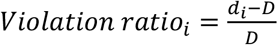, where *d_i_* is the distance between the contact loci in the model, and *D* is the target contact distance (2 × *R^soft^*).
3. Residual ratio. The residual ratio Δ*r* is defined as:

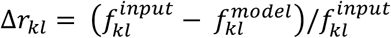

with 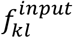 and 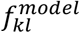 as the contact probabilities between regions *k* and *l* from experiment and models, respectively. Residual ratios are very small, and centered at a median of 0.03 (mean= −0.05) for intra-chromosomal and 0.001 (mean = −0.002) for inter-chromosomal contacts (Fig. S1), showing excellent agreement between experiment and model.
4. Prediction of missing Hi-C data from sparse data model. A sparse Hi-C input data set is generated by randomly removing 50% of the non-zero data entries from the Hi-C contact frequency matrix.

#### ▪ Robustness and Converge Analysis

##### Replicates

Technical replicates are calculated from different random starting configurations. Resulting contact frequency maps and the average radial positions of all chromatin regions between replica populations are nearly identical (Fig. S2). All observed structural features discussed in this paper are reproduced in the technical replicate population.

##### Population size

To test convergence with respect to population size, we generate 5 different populations with 50, 100, 1,000, 5,000 and 10,000 structures. Chromatin contact frequencies and structural features for each structure populations are compared against results with a population size of 10,000 structures. At a population of 1,000 structures, a size much smaller than our target population, contact frequency values and average radial positions are already converged to a very high correlation with those from a 10,000 structure population (Fig. S3).

### Chromatin interaction networks and identification of spatial partitions

#### ▪ Building chromatin interaction networks

A chromatin interaction network (CIN) is calculated for each model and for chromatin in each subcompartment separately as follows: Each vertex represents a 200-kb chromatin region. An edge between two vertices *i*, *j* is drawn if the corresponding chromatin regions are in physical contact in the model, if the spatial distance *d_ij_* ≤ 2 × (*R^soft^*).

##### Network properties

###### Maximal Clique Enrichment

A clique is a subset of nodes in a network where all nodes are adjacent to each other and fully connected. The maximal clique refers to the clique that cannot be further enlarged. The number of maximal cliques, *c*, is calculated using the *graph_number_of_cliques* function in the *NetworkX* python package^76^. The maximal clique enrichment (MCE) of the subcompartment *s* in the structure *m* is calculated as:

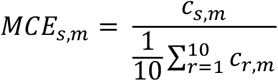

Where *c_s,m_* is number of maximal cliques for subcompartment s in structure *m*; *c_r,m_* is the number of maximal cliques of a CIN constructed from randomly shuffled subcompartment regions in the same structure *m*. High MCE values shows formation of a structural subcompartment with high connectivity between 200-kb regions of the same state.

###### Neighborhood Connectivity

To calculate the neighborhood connectivity (NC) of a subcompartment CIN, we first calculate the average neighbor degree for each node using the *average_neighbor_degree* function in the *NetworkX* python package^76^. The overall neighborhood connectivity of the subcompartment s in the structure *m* is then calculated as:

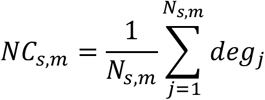

where *N_s,m_* is the number of nodes in the CIN of the subcompartment*s s* in the structure *m*, and *deg_j_*, is the average neighbor degree of node *j*.

#### ▪ Identifying spatial partitions via Markov clustering

Spatial partitions of subcompartments as well as regions in A compartment with low and high structural variability (A regions in the first and last quartile based on their radial position variability, see *Cell-to-cell variability of features section* below) are identified by applying Markov Clustering Algorithm (MCL)^77^, a graph clustering algorithm, which identifies highly connected subgraphs within a network. MCL clustering is performed for each subcompartment CIN in each structure by using the *mcl* tool in the *MCL-edge* software^77^. Unless otherwise noted, the 25% smallest subgraphs (with less than 7 nodes, many of those singletons) are discarded from further analysis to focus on highly connected subgraphs. The highly connected subgraphs are referred to as “spatial partitions” throughout the text.

In addition to subcompartment/compartment partitions, we also predict speckle, and nucleoli partitions as follows:

i. Speckle partitions: <u>Case 1</u>: *Predictions of speckle locations with knowledge of A1 subcompartment annotations* Speckle locations are identified as the geometric center of A1 spatial partitions identified by Markov clustering of A1 CINs. In each structure, A1 spatial partitions are considered with sizes larger than 3 nodes (chromatin regions). <u>Case 2</u>: *Predictions of speckle locations without knowledge of subcompartments* We first identify chromatin expected to have high speckle association. These regions are identified as those with unusually low and stable interior radial positions. We select 10% chromatin regions with the lowest average radial positions. (78.4% of these regions are part of the A1 subcompartment). We then generate CINs for the selected group of chromatin regions in each structure of the population. Approximate speckle locations are then identified as the geometric center of the resulting spatial partitions identified by Markov clustering of the CINs. Spatial partitions are considered with sizes larger than 3 chromatin regions. <u>Case 3</u>: *Predictions using locations of A2 partition centers* For comparison, we also identify speckle locations as the geometric center of A2 spatial partitions identified by Markov clustering of A2 CINs similar to Case 1. In each structure, A2 spatial partitions are considered with sizes larger than 3 chromatin regions.
ii. Nucleoli partitions: Following the same protocol as in Case 2 for speckle partitions, we first identify chromatin expected to have high nucleoli association. These regions are identified as those previously reported nucleoli associated domain (NAD)^68^ regions and nucleolus organizing regions (NOR, on short arms of chromosomes 13, 14, 15, 21, and 22). Using these regions, we generate CINs in each structure of the population. Approximate nucleoli locations are then identified as the center of mass of the resulting spatial partitions identified by Markov clustering of the CINs. Only top 25% largest spatial partitions are used as predicted nucleoli. For NOR regions, we use the first 25 restrained 200-kb regions that are closest in sequence to NOR regions in these five chromosomes, as NOR regions do not have Hi-C data and they are not restrained during the modeling protocol.

##### Properties of partitions

###### Size of partitions

The size of a spatial partition is calculated as 0.2 *x N* Mb where *N* is the number of nodes in the partition that represents a 0.2 Mb region.

###### Fraction of inter-chromosomal edges (contacts)

For each spatial partition, the inter-chromosomal edge fraction (ICEF) is calculated as:

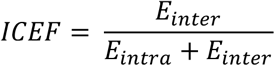

where *E_intra_* and *E_inter_* are number of intra- and inter-edges in the partition, respectively.

### Structural features

Unless otherwise noted, mean values of structural features for each genomic region are calculated from 2 copies and 10,000 structures (total 20,000 configurations) in the following structural feature calculations.

#### ▪ Mean radial position (RAD, #1)

Radial position of a chromatin region *i*. in structure *m* is calculated as:

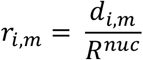

where *d_i,s_* is the distance of *i* to the nuclear center, and *R^nuc^* is the nucleus radius which is 5 μm. *r_i,s_* = 0 means the region *i* is at the nuclear center while *r_i,s_* = 1 means it is located at the nuclear surface.

##### Other radial position related analyses

i. *Overlap of subcompartment borders and large radial position transitions*: To identify regions coinciding with large transitions in radial positions, we first calculate each region’s gradient in radial position from their average radial position profiles. Peaks and valleys in the gradient profile coincide with the regions of large radial transitions in the chromosome and are identified with the *detect_peaks* python package^78^. We obtain 1408 regions with large radial transitions with minimum peak height (mph) set to 0.01 (the gradient values range between −0.06 – 0.05.) to filter out regions with minimal radial transitions. We then check if these identified regions coincide with the subcompartment borders, i.e. where two neighboring chromatin regions are in different subcompartments. We determine an overlap if there is a subcompartment border within a 1-Mb window of a given identified region with a large radial transition.
ii. *Shell analysis:* To map the preferred positions of 200-kb regions in the nucleus, we divide the nuclear volume of each model into 5 concentric shells *L* = {*L*_1_,*L*_2_,*L*_3_,*L*_4_,*L*_5_} so that each shell contains the same amount of chromatin in each single structure. We then calculate the probability of a subcompartment *s* to be in any shell from *L*:

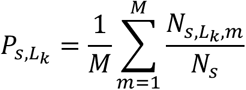

where *N_s,L_k_,m_* is the number of regions from subcompartment *s* in shell *L_k_* in structure *m*, *N_s_* is the total number of regions in subcompartment *s*, and *M* is the total number of structures.
iii. *Comparison with GPSeq:* GPSeq scores^41^ are rescaled to have values between 0 – 1, where scores 0 and 1 correspond to a chromatin region being at the nuclear lamina and nuclear center, respectively^41^. Average radial positions extracted from our structures vary between 0.48 – 0.94 with higher values corresponding to proximity to nuclear lamina. For comparison with GPSeq, we subtract the average radial positions from 1 and then rescale the values to be between 0 – 1.
iv. *Average radial positions of regions from different replication phases:* Genomic regions are divided into 6 groups (G1b, S1, S2, S3, S4, G2) based on their mapped replication phases^63^. For each group, the distribution of the average radial positions is then determined from the structure population.

#### ▪ Local chromatin fiber decompaction (RG, #2)

##### Radius of gyration of chromatin fiber

The local compaction of the chromatin fiber at the location of a given locus is estimated by the radius of gyration (RG) for a 1 Mb region centered at the locus (i.e. comprising +500kb up- and 500 kb downstream of the given locus). To estimate the RG values along an entire chromosome we use a sliding window approach over all chromatin regions in a chromosome.

The RG for a 1 Mb region centered at locus *i* in structure *m*, is calculated as:

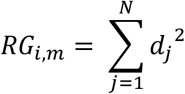

where *N* is the number of chromatin regions in the 1-Mb window, and *d_j_* is the distance between the chromatin region *j* to the center of mass of the 1-Mb region.

##### Other RG related analysis

i. *TAD border detection:* To investigate if chromatin regions with maxima in RG profiles coincide with TAD borders, we first identify peak regions in the average RG profiles with the *detect_peaks* python package^78^. 2068 peak regions are detected genome-wide with minimum peak distance (mpd) set to 3 (peaks must be at least 3 data points/600-kb apart from each other). We then check if these identified regions coincide with TAD borders detected by TopDom^75^, HiCseg^79^, InsulationScore^80^, and TADbit^56^. We determine an overlap if there is a TAD border within ±200-kb window of a peak region.
ii. *RG peak frequency:* Peak regions in the RG profiles are detected in each individual structure using *detect_peaks* python package^78^ with same parameters as in the previous section. The RG peak frequency (PF) of a region *i*. is then calculated as:

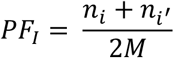

where *n_i_* and *n_i_*′ are the number of structures in which region *i*. and its homologus copy has an RG peak, and *M* is the number of genome structures in the population.

#### ▪ Mean gene-speckle and gene-nucleolus distances (SpD, NuD, #3,4)

For each 200-kb region, the closest speckle partition (or nucleolus partition) in each single structure is identified and the center-to-center distance is calculated (from the center of the region to the geometric center of the partition). The distances across the population are then averaged for each region to calculate mean speckle (or nucleolus) distances.

##### Other related analysis

###### Speckle distance heatmaps

A speckle distance heatmap for a chromosome visualizes, for a given chromatin region, the speckle distance variability across the population of models. For each copy of a chromatin region, the distance to the nearest predicted speckle is calculated in each structure of the population. These distances (20,000 distances total due to 2 copies and 10,000 structures) are ranked from lowest to highest values and plotted along a column of the speckle distance heatmap and color coded according to the distance. Colors range from low distance (red) to large distances (blue).

#### ▪ Cell-to-cell variability of features (δ_RAD_, δ_RG_, δ_SpD_, δ_NuD_, #5-8)

Cell-to-cell variability of any structural feature (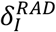 for radial positions, 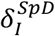 speckle distances, 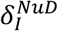 nucleoli distances, and 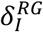 local decompaction) for a chromatin region *I* is calculated as:

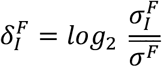

where 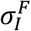 is the standard deviation of the values for structure feature *F* calculated from both homologous copies of the region across all 10,000 genome structures in the population; 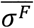 is the mean standard deviation of the feature value calculated from all regions within the same chromosome of region *I*. Positive 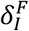 values 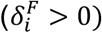 result from high cell-to-cell variability of the feature (e.g. radial position); whereas negative values 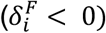 indicate low variability.

Regions in A compartment with positive and negative 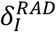 are called A-HV (high variability) and A-LV (low variability), respectively. Likewise, regions in B compartment with positive and negative 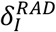 are called B-HV and B-LV, respectively. The number of 200-kb regions in each group are 3164, 2731, 3839, and 3918 for A-LV, A-HV, B-LV, and B-HV, respectively.

#### ▪ Interior localization frequency (ILF, #9)

For a given 200-kb region, the interior localization frequency (ILF) is calculated as:

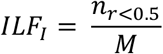

where *n*_*r*<0 5_ is the number of structures where either copy of the region *I* has a radial position lower than 0.5, and *M* is the total number of structures which is 10,000 in our population.

#### ▪ Nuclear-body association frequencies (SAF, LAF, NAF, #10-12)

For a given 200-kb region, the association frequency to nuclear bodies (SAF, LAF, and NAF for speckle, lamina, and nucleoli association frequencies, respectively) are calculated as:

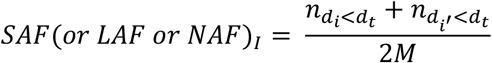

where *M* is the number of structures in the population (2 homologous copies of each chromosome are present per structure); *n*_*d*_*i*_<*d_t_*_ and *n*_*d_i_*′<*d_t_*_ are the number of structures, in which region *i*. and its homologous copy *i*′ have a distance to the nuclear body of interest (NB) smaller than the association threshold, *d_t_*. The *d_t_*s are set to 500 nm, 0.35*xR_nuc_*, and 1000 nm for SAF, LAF, and NAF, respectively. We try different distance thresholds, and the select thresholds resulted in the best correlations with experimental data. For SAF and NAF calculations, we use the predicted speckle and nucleolus partitions to calculate distances (see *Identifying spatial partitions via Markov clustering*). For LAF, we use the direct distances of regions to the nuclear envelope. For all association frequency calculations, we calculate distances from the surface of the region to the center-of-mass of the partition or to the surface of the nuclear envelope.

##### Other related analyses

i. *Predicting lamin B1 DamID signals using LAF*: The predicted laminaDamID signal of region *I* is calculated as:

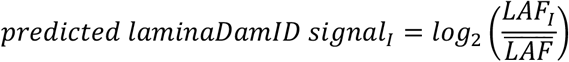

where 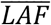 is the mean lamina association frequency calculated from all regions in the genome.
ii. *Comparison with imaging data*: We compare our SAF, LAF and NAF values with imaging data^8^. To calculate association frequencies from imaging and models, we use different distance thresholds (250, 500, 750, 1000 nm distance thresholds for SAF and LAF when calculated from imaging or models, and additional thresholds of 1250, 1500, 1750, 2000 nm for LAF when calculated from models) to define an association to the nuclear body of interest. We find that the best correlations are obtained when the following distance thresholds are used:

- SAF: 500 nm for imaging, 750 nm for models
- NAF: 1000 nm for imaging, 1000 nm for models
- LAF: 1000 nm for imaging, 2000 nm for models For SAF comparisons, we use the predicted speckle partitions from interior regions (Case 2 for speckle partitions in *Identifying spatial partitions via Markov clustering*).

#### ▪ TSA-seq (S-TSA, L-TSA, N-TSA, #13-15)

To predict TSA-seq signals for speckle, nucleoli, and lamina from our models, we use the following equation:

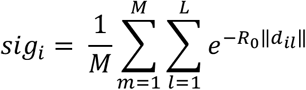

where *M* is the number of models, *L* is the number of predicted speckle locations in structure *m*, *d_ii_* is the distance between the region *i*. and the predicted nuclear body location *l*, and *R*_0_ is the estimated decay constant in the TSA-seq experiment^14^ which is set to 4 in our calculations. The normalized TSA-seq signal for region *i*. then becomes:

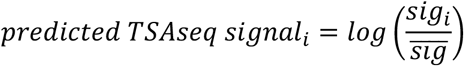

where 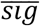 is the mean signal calculated from all regions in the genome. The predicted signal is then averaged over two copies for each region. The predicted speckle, and nucleoli partitions are used for distance calculations (see *Identifying spatial partitions via Markov clustering*). For lamina TSA-seq, we use direct distances of each 200-kb chromatin region to the nuclear surface in each structure, which is calculated as (1 – *r_m_*) × *R_nuc_* where *r_m_* is the radial position of the 200-kb region in structure *m* and *R_nuc_* is the nucleus radius which is set to 5 μm.

##### Other related analysis

i. *Predicting SON TSA-seq signals using only cis relationships in folded chromosomes*: To identify contributions of cis interactions in SON TSA-seq signals, speckle locations are defined by the geometric center of consecutive A1 sequence blocks formed by more than 1 A1 chromatin region (instead of the geometric center of A1 spatial partitions, which can be formed by both cis and trans chromosomal interactions). For single A1 regions, the bead center location is used instead. For each chromatin region, we then calculate its spatial distances to these predicted speckle locations in the folded chromosome, which are used to predict the resulting TSA-seq signals from cis interactions only.
ii. *Predicting SON TSA-seq signals using only cis relationships in random conformations*: We also repeat the same calculations as defined in the previous section, but instead of the folded chromosomes, use models with random chain configurations, generated without Hi-C data (i.e. only chain connectivity and excluded volume). TSA-seq data is calculated accordingly from the corresponding distances based on the random polymer chain configurations.
iii. *Predicting SON TSA-seq signals using speckle distances based on A1 sequence locations*: Speckle locations are approximated by the sequence positions of A1 regions, either as median sequence position for a block of consecutive A1 chromatin regions or the sequence positions of individual A1 regions, if their neighboring regions are not part of the A1 subcompartment. The distance 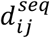 between a chromatin region *i* and speckle position *j*, separated in sequence by *n* chromatin regions, is then defined as 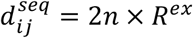, where *R^ex^* = 118 *nm* is the excluded volume radius of a chromatin region in the models (see *Genome representation*). These distances are then used to predict SON TSA-seq signals as defined above.
iv. *Histone modification histograms based on predicted SON TSA-seq deciles*: Following the procedure described in ref^14^, we divide the 200-kb chromatin regions in our models into 10 decile groups based on their predicted SON TSA-seq signals; deciles 1 and 10 contain regions with the lowest and highest 10% predicted TSA-seq signals, respectively. We then count the number of mapped peaks of H3K27me3, H3K4me3, and H3K9ac as well as the number of A1, A2, A1+A2 regions in each decile, and calculate the fraction of histone modification peaks or A1/A2 regions accrued in each decile. For mapping histone modification peaks to 200-kb bins to match our models’ resolution, see *Mapping experimental data to models* in *Supplementary Information*. Same histograms using experimental TSA-seq deciles are re-generated from Fig. 8 in ref^14^ using WebPlotDigitizer^81^.

#### ▪ Mean inter-chromosomal neighborhood probability (ICP, #16)

For each target chromatin region *i*, we define the neighborhood {*j*} if the center-to-center distances of other regions {*j*} to the target region are smaller than 500 nm, which can be expressed as a set; *Ne_i_* = {*j*: *j* ≠ *i*, *d_ij_* < 500 *nm*}. Inter-chromosomal neighborhood probability (ICP) is then calculated as:

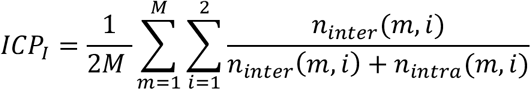

where *M* is the number of structures, *n_intra_*(*m*, *i*) and *n_inter_*(*m*, *i*) are the number of intra- and inter-chromosomal regions in the set *Ne_i_* in structure *m* for haploid region *i*.

#### ▪ Median trans A/B ratio (#17)

For each chromatin region *i*, we define the trans neighborhood {*j*} if the center-to-center distances of other regions from other chromosomes to itself are smaller than 500 nm, which can be expressed as a set; 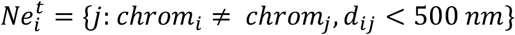. Trans A/B ratio is then calculated as:

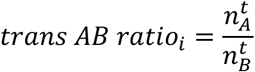

where 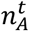 and 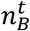 are the number of trans A and B regions in the set 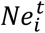 for haploid region *i*. The median of the trans A/B ratios for a region is then calculated from all the trans A/B ratios of the homologous copies of the region observed in all the structures of the population. The values are then rescaled to have values between 0 – 1.

### Comparison of gene expression with structural features

#### ▪ Transcription frequency

Transcription frequency (TRF) of each gene in the single cell RNA-seq (scRNA-seq) data is defined as the fraction of cells in the population of cells, where the gene has non-zero mRNA transcription counts in the scRNA-seq data^42^. TRF is also calculated from the recently published nascent RNA-MERFISH imaging data as the fraction of cells where the gene is transcribed (transcription: on) in the population of imaged cells^8^.

#### ▪ Gene expression heatmaps

Gene expression heatmaps for each chromosome visualize the variability of mRNA counts (the expression levels) for each gene in a population of cells^42^. For each chromatin region, the observed mRNA count in each cell of the population of models is ranked from highest to lowest values and plotted along a column. Colors ranged from high mRNA counts (red) to 0 (dark blue).

#### ▪ ROC curve for assessing performance to classify lowly or highly expressed genes

We first identify the top 10% (T10) and the bottom 10% (B10) genes with the highest and the lowest total non-zero mRNA counts (i.e. gene expression values) in the scRNA-seq data^42^. Several structural features (mean radial positions, ILF, mean speckle distances, SAF, variability of radial positions and speckle distances) are then calculated for all chromatin regions mapped to T10 genes and B10 genes.

To determine the most informative structural features for distinguishing T10 genes from B10 genes, we perform receiver operator characteristic (ROC) analysis. Specifically, for each structural feature, we define 10 threshold levels, equally separating the range of values for each structural feature. Then we determine how well the gene in the T10 and B10 groups are separated by each threshold value by calculating the corresponding number of true positives/negatives (TP, TN) and false positive/negatives (FP, FN).

For each structural featue *f* and for each threshold level, *t*, the true positive rate (TPR) and false positive rates (FPR) are then calculated as

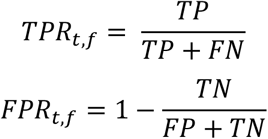

The ROC curves are then plotted for each feature using TPR/FPR values.

### Other structural analyses

#### ▪ Experimental GRO-seq and TSA-seq data analysis

##### Averaging TSA-seq and GRO-seq signals in concentric shells around subcompartment partitions

To quantify average TSA-seq^14^ and GRO-seq^67^ signals for chromatin with respect to the distance to spatial partition centers of each subcompartment, the nuclear volume around a spatial partition center is divided into concentric shells, with each consecutive shell radius increasing by 200 nm. The signals are then averaged over concentric shells around partition centers as follows: In each individual genome structure, the signals of chromatin located in the same shell volume is averaged, irrespective of the chromatin’s subcompartment assignment. The average signal per shell are further averaged over all partition centers in the same subcompartment and over all structures of the population. Note that this measure only relies on the geometric position of a partition center and the folded genome (i.e. calculates average gene expression from all chromatin in a shell, independent of subcompartment annotations).

#### ▪ Neighborhood composition

The neighborhood composition (NeC) shows how frequent chromatin regions in different subcompartments are in spatial proximity to regions of a specific subcompartment. The average percentage of subcompartment *Q* in the neighborhood composition of subcompartment *S* in the population is calculated as:

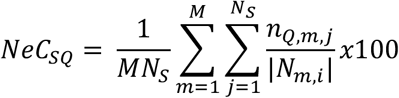

where *M* is the number of structures in the population, *N_s_* is the number of 200-kb regions belonging to subcompartment *S*, {*N_m,i_*}is the set of 200-kb chromatin regions in the neighborhood of the region *i*. in structure *m*, and *n_Q,m,i_* is the number of chromatin regions from subcompartment *Q* in the set {*N_m,i_*}. We define the neighborhood of *i*. in structure *m* as *N_m,i_* = {*j*: *j* ≠ *i*, *d_ij_*, < 500 *nm*}, which contains the list of all chromatin regions with less than 500 nm center-to-center distance (*d_ij_*) to chromatin region *i*.

The neighborhood composition enrichment (NeCE) of subcompartment *Q* in the neighborhood of subcompartment *S* is calculated as:

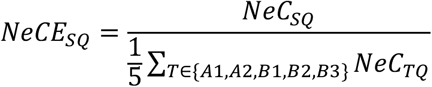

where *NeC_SQ_* is the neighborhood composition percentage calculated for subcompartment *Q* in the neighborhood of subcompartment *S* and the denominator is the average percentage of subcompartment *Q* observed in the neighborhood of all subcompartments. Values greater than 1 (*NeCE_SQ_* > 1) indicate that subcompartment *Q* is enriched in the neighborhood of subcompartment *S*, whereas values lower than 1(*NeCE_SQ_* < 1) show depletion of *Q* around *S*.

#### ▪ Enrichment heatmaps for various features

##### Enrichment of structural features, experimental TSA-seq, DamID, and GRO-seq signals, and histone modifications in various groups

To identify structural feature or experimental signal enrichments for chromatin in different groups (subcompartments, TSA-seq deciles, superenhancers, enhancers, replication phases, A/B-LV/HV groups, and T10/B10 genes), we first normalize each feature value to range between 0 and 1. We then calculate the enrichment of a structural feature *f*, for group *g* as:

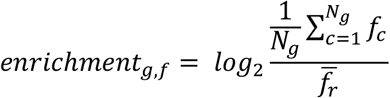

where *N_g_* is the number of 200-kb chromatin regions in group *g*, *f_c_*. is the structure feature value for chromatin region *c*. For 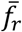, we first randomly select the same number (*N_g_*) of regions in the genome and calculate the average feature value, and repeat this step 1000 times. For the enrichment of histone modifications in A-LV and A-HV groups, we randomly select the same number of regions only from regions in compartment A. We then take the average of 1000 different average feature values calculated from randomly selected regions.

For visualization purposes, we reverse the ranges of radial positions, mean-speckle, and mean-nucleoli distances in the structural feature enrichment heatmaps, so lower values would be indicated with red.

##### Enrichment of replication phases, LADs, and subcompartments in A-LV, A-HV, B-LV, and B-HV groups

We calculate the enrichment of various tags *t* (based on replication phases, LADs, or subcompartments), in group *g* as:

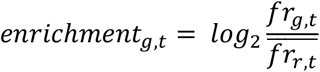

where *fr_g,t_* is the fraction of regions with tag *t* in group *g*. For 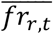, we first randomly select the same number of genomic regions (as in group *g*) and calculate the fraction of regions with tag *t* among those regions, and repeat this step 1000 times. We then take the average of 1000 different fraction values.

#### ▪ K-means clustering of A and B compartments

For clustering, we first normalize all 17 structural features using log_2_-transformation. We then perform K-means clustering using all transformed features for A and B subcompartments separately. We use scikit-learn python package to perform K-means clustering^82^ and set the *n_clusters* parameter to 2 for A and 3 for B compartments. Clusters are then compared with actual subcompartment assignments to compute clustering accuracy. The highest prediction accuracies are obtained when clustering is performed with a subset of structural features for both A and B subcompartments. The used features in the clustering are cell-to-cell variability of radial positions, SAF, NAF, median trans A/B ratios for A, and cell-to-cell variability of radial positions and nucleoli distances, nucleoli TSA-seq, ICP, median trans A/B ratios for B subcompartment predictions, respectively.

#### ▪ Comparison with 3D in situ hybridization (3D-FISH) data

FISH probes are mapped to 200-kb chromatin regions in our models according to the highest overlap. Radial positions and pairwise distances for each mapped probe are determined in each structure in the population and compared to the radial positions and pair distances in FISH experiments. FISH and model radial positions are normalized by their maximum values. Intra-chromosomal distances in models are defined by their surface-to-surface distances of the corresponding probe regions (in both copies of the chromosome). Colocalization fraction of inter-chromosomal pairs are calculated as following: first the center-to-center distances of all possible probe pairs (*i* – *j*, *i* – *j*′, *i*′ – *j*, *i*′ – *j*′ where *i*′ and *j*′ are the homologous copies of each 200-kb chromatin regions, *i*. and *j*) are calculated in each structure. The minimum distance from all possible pairs in each structure is then used to calculate the fraction of models in which both regions are colocalized. We assume a loci pair is colocalized in a structure if the calculated minimum distance in that structure is lower than 1 μm (*d_min_* < 1 *μm*).

#### ▪ Radial positions of trans and cis interactions

We select 1,000 random structures from the population and identify all the trans and cis chromatin interactions. Then we calculate the average radial position of the location where the trans or cis interaction occurs by taking the mean of the radial positions of the two loci forming the interaction in that structure.

## Data visualization

CINs are visualized by Cytoscape^83^. 3D models and spatial partitions are visualized by using Chimera^84^.

