## Supplementary Information for "Population-based structure modeling reveals key roles of nuclear microenviroment in gene functions"

**Table of Contents**

1. SI Figures/Tables

   **Fig. S1.** Residual ratios

   **Fig. S2.** Comparison of replicate structure populations

   **Fig. S3.** Population size convergence plots

   **Fig. S4.** Chromatin interaction networks for subcompartments

   **Fig. S5-25**. Structural feature profiles for Chr2-22.

   **Table S1**. Pearson and stratum adjusted correlation coefficients between input/output for each chromosome.

   **Table S2**. Experimental data used

2. Preprocessing Hi-C data
3. Iterative refinement
4. Mapping experimental data to 200 kb
5. FISH experiments

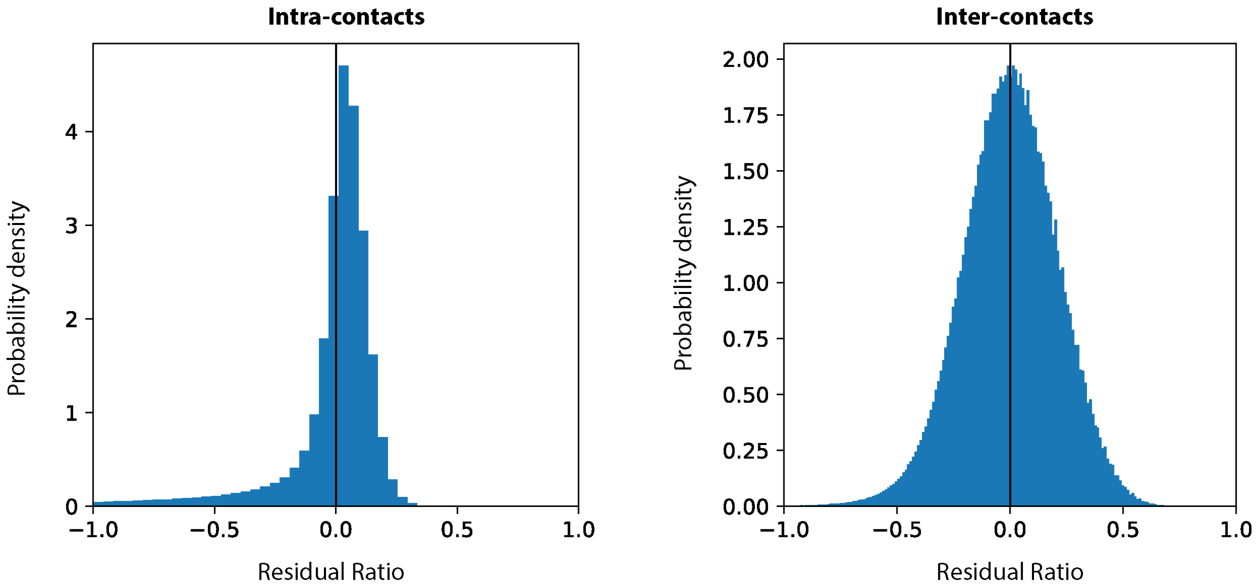


**Fig. S1.** Residual ratios. The residual ratio $\Delta r$ is defined as $\Delta r_{kl} = \left(f_{kl}^{input} - f_{kl}^{model}\right)/f_{kl}^{input}$ with $f_{kl}^{input}$ and $f_{kl}^{model}$ as the contact probabilities between regions *k* and *l* from experiment and models, respectively. Residual ratios are very small, and centered at a median of 0.03 (mean= -0.05) for intra-chromosomal (left) and 0.001 (mean = -0.002) for inter-chromosomal (right) contacts, showing excellent agreement between experiment and model.

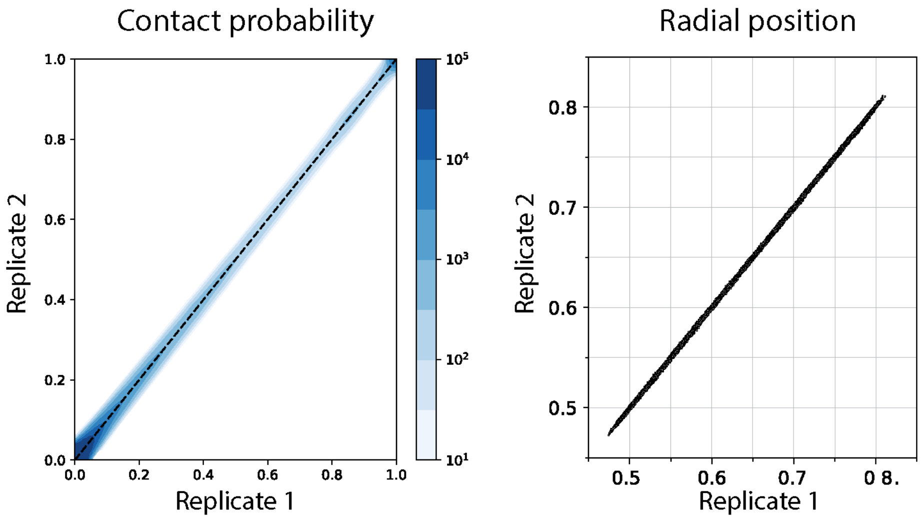


**Fig. S2.** Comparison of replicate populations. Scatter plots of contact probabilities (left, Pearson corr. = 0.99, p<0.0001) and radial positions (right, Pearson corr. = 0.99, p<0.0001) for two replicates.

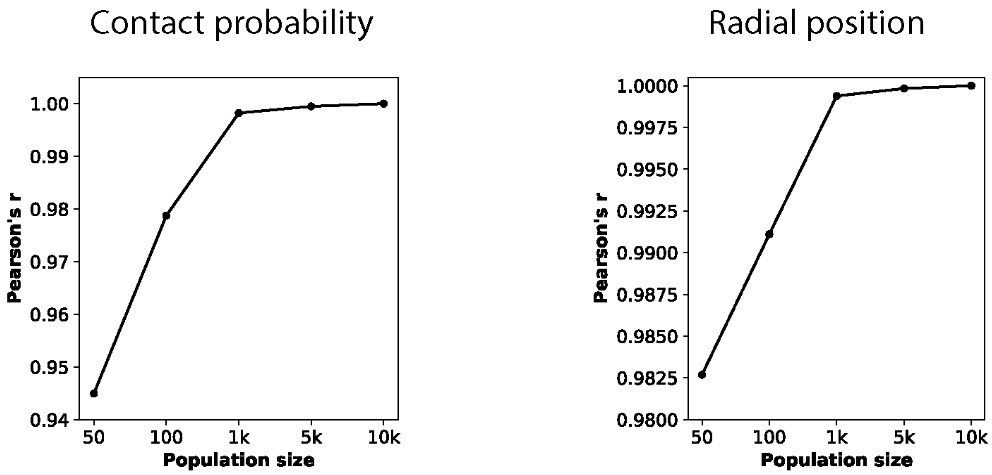


**Fig. S3.** Population size convergence plots. Pearson correlations between the population with 10,000 structures and populations with smaller sizes (50, 100, 1,000, and 5,000 structures). Contact probabilities (left) and radial positions of chromatin regions (right) already converge at 1,000 structures and have very high correlations (>0.99) with the 10,000 structure population.

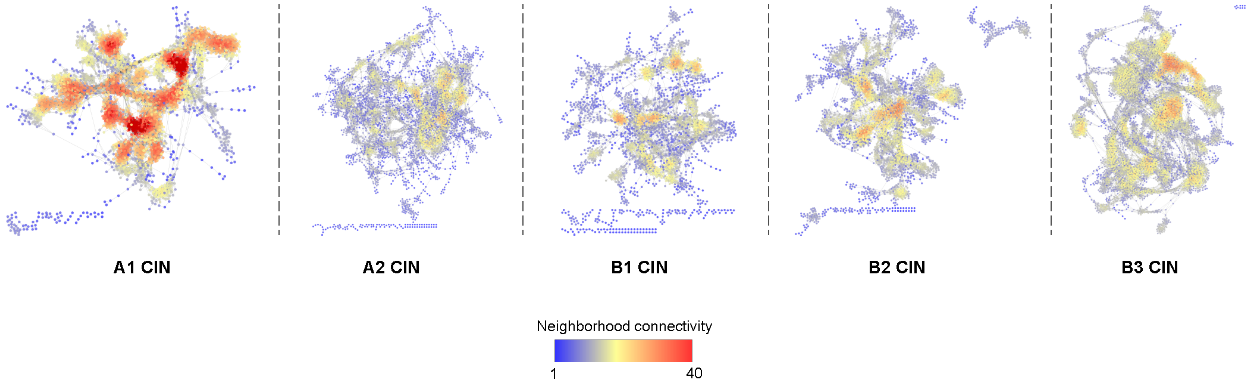


**Fig. S4.** Representative chromatin interaction networks (CIN) for chromatin in each subcompartment in a single structure. Each node in CINs represents a single chromatin region connected by edges if the two regions are in physical contact in the 3D structure. Nodes are colored by their neighborhood connectivity (i.e. the average contacts formed by their neighbor nodes) ranging from low (blue) to high (red).

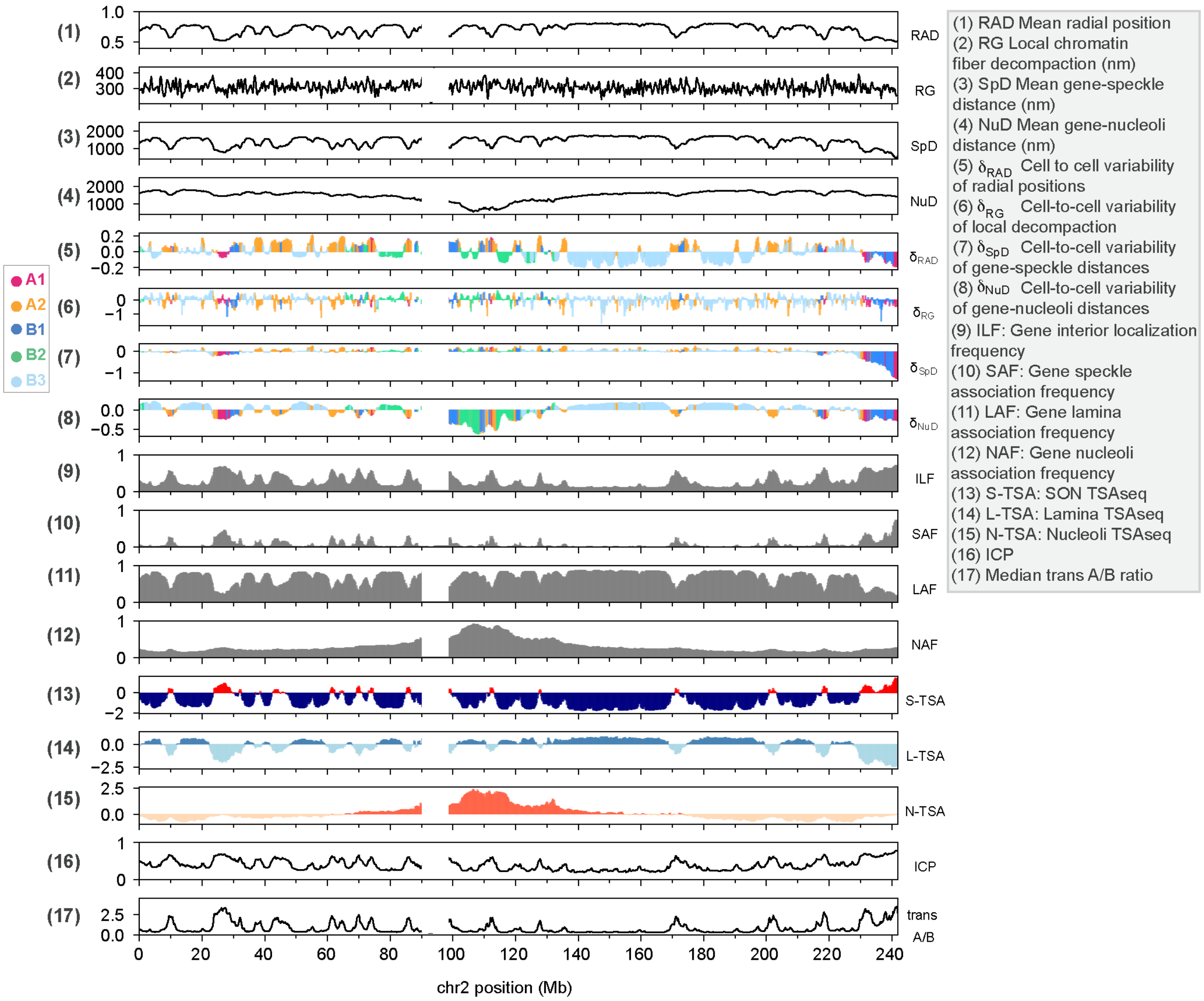


**Fig. S5.** Structure feature profiles for chromosome 2.

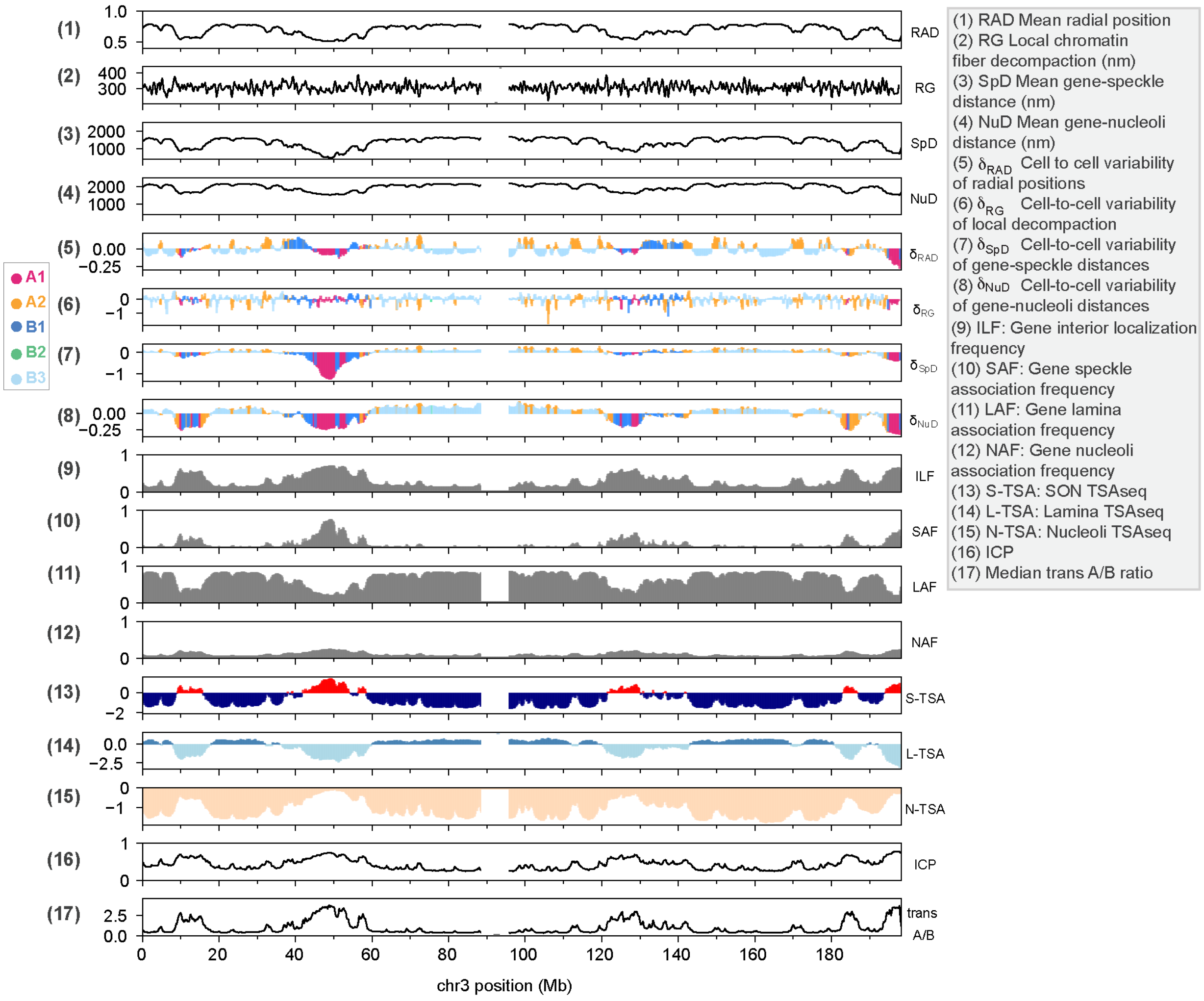


**Fig. S6.** Structure feature profiles for chromosome 3.

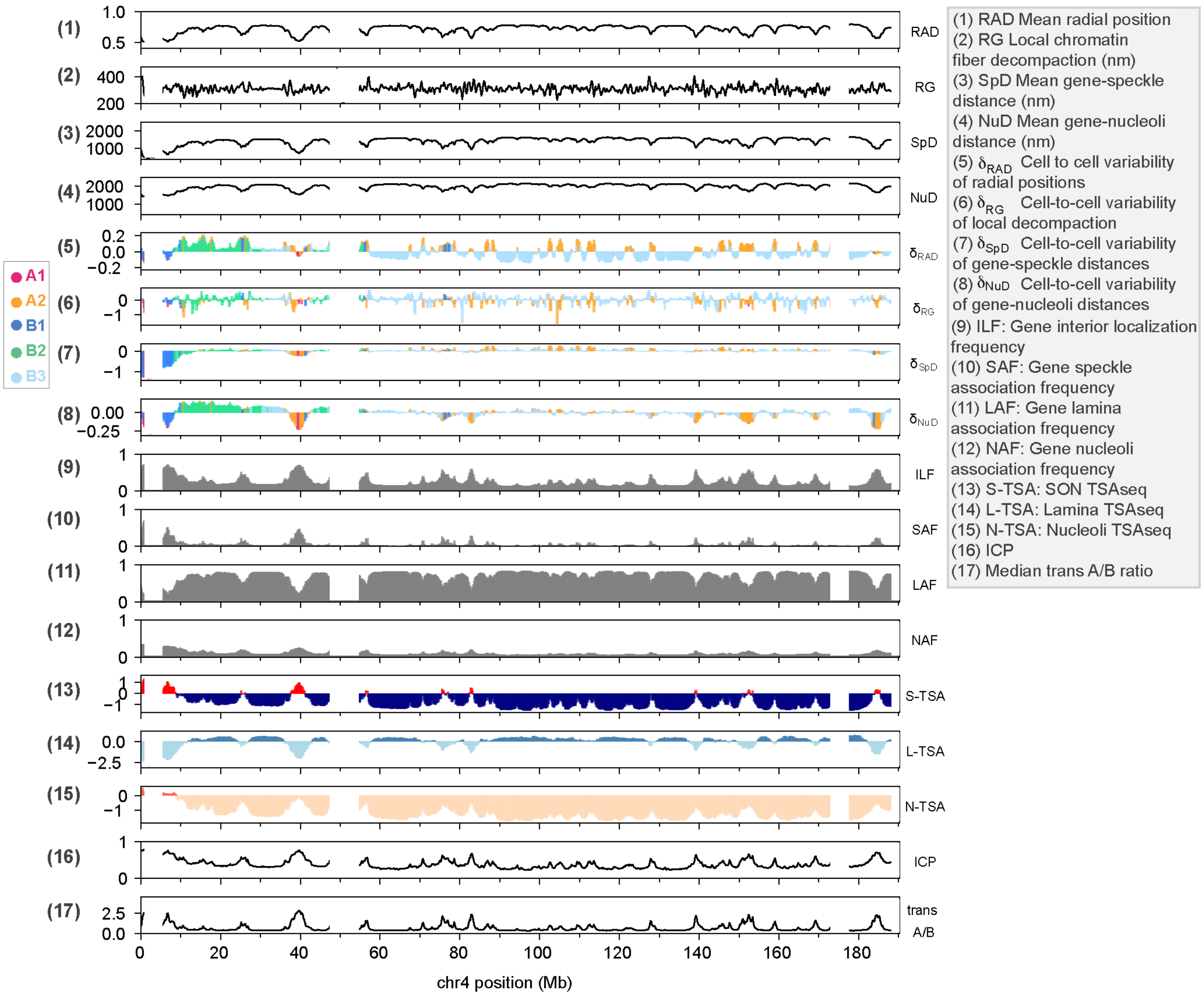


**Fig. S7.** Structure feature profiles for chromosome 4.

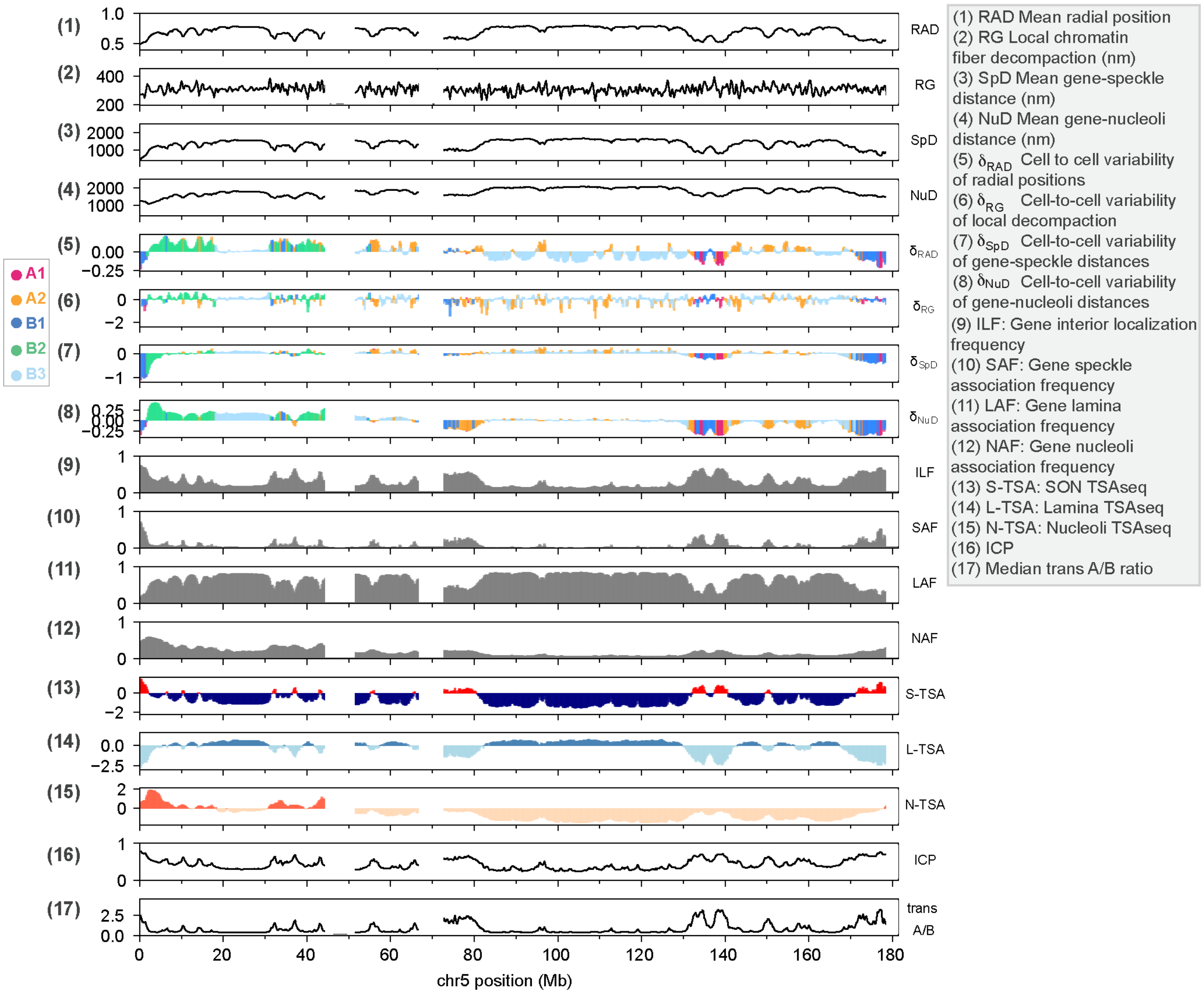


**Fig. S8.** Structure feature profiles for chromosome 5.

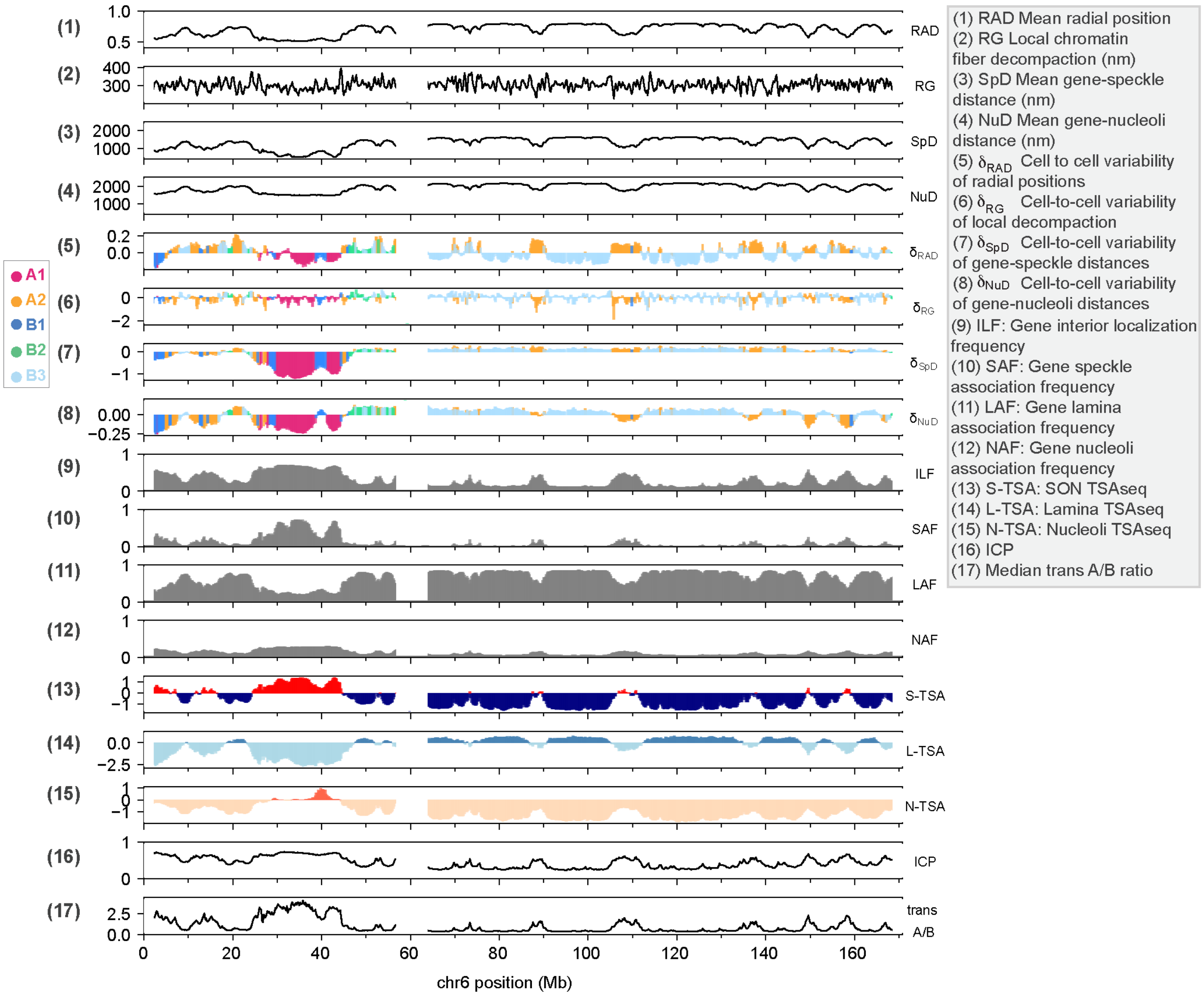


**Fig. S9.** Structure feature profiles for chromosome 6.

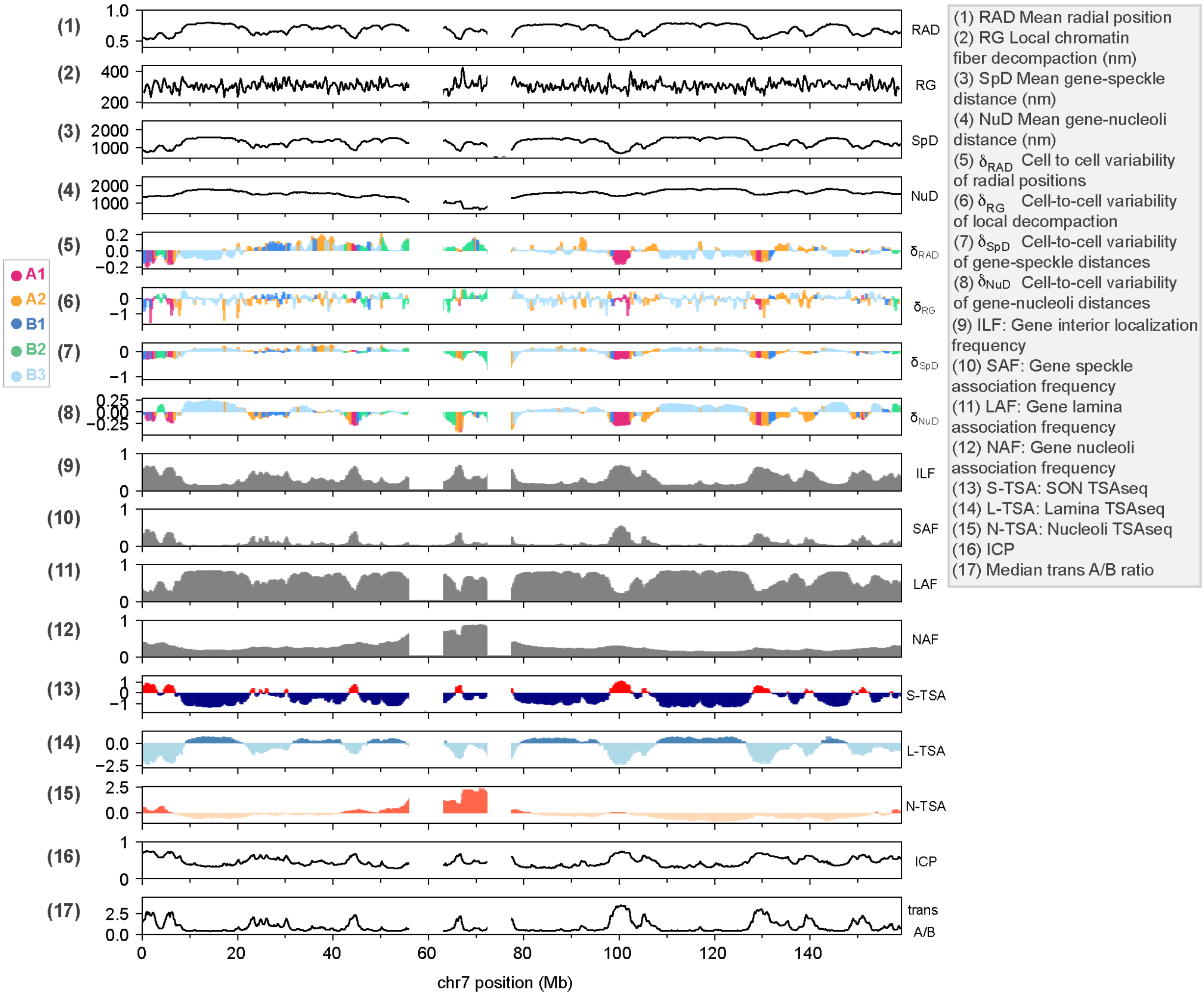


**Fig. S10.** Structure feature profiles for chromosome 7.

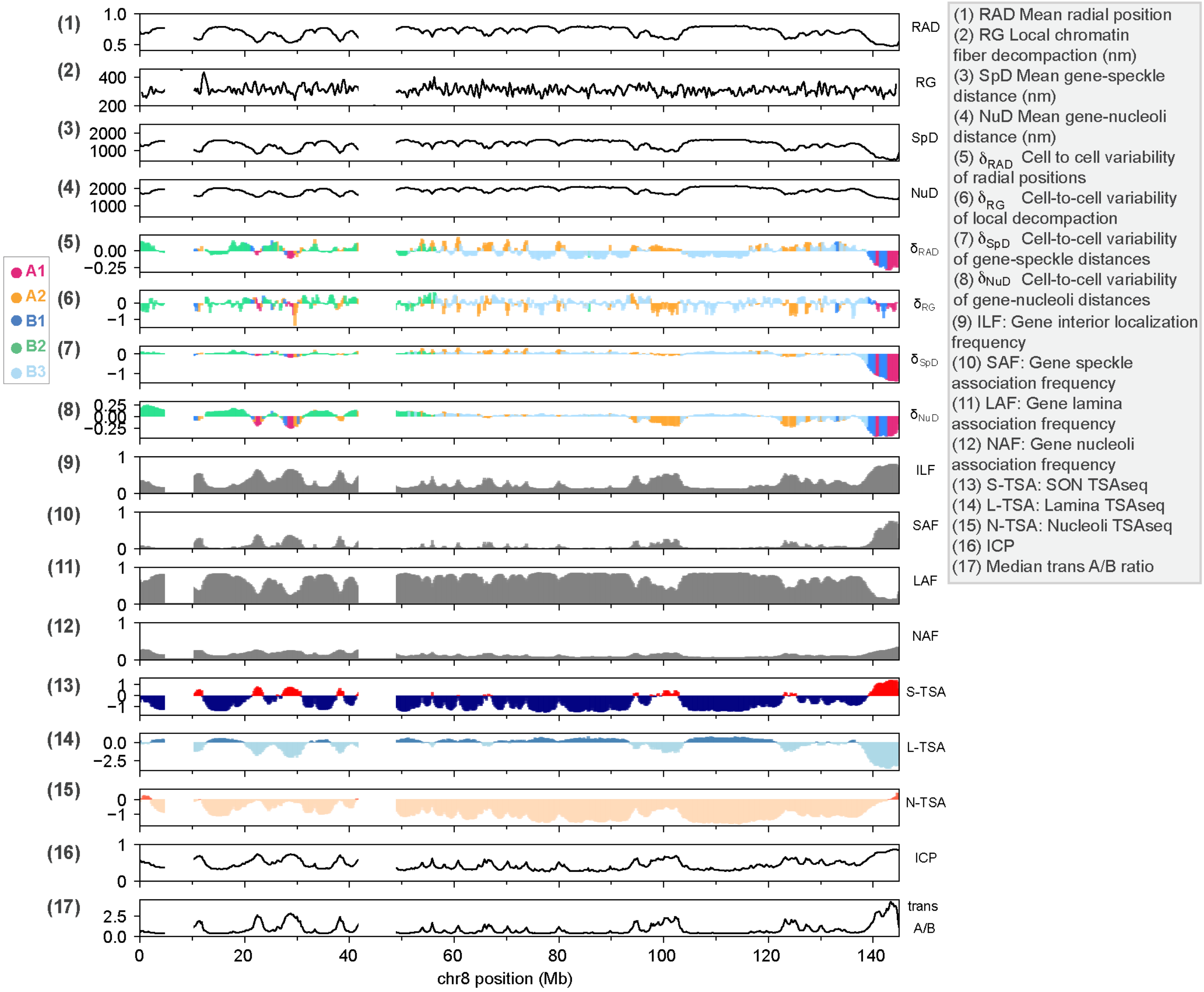


**Fig. S11.** Structure feature profiles for chromosome 8.

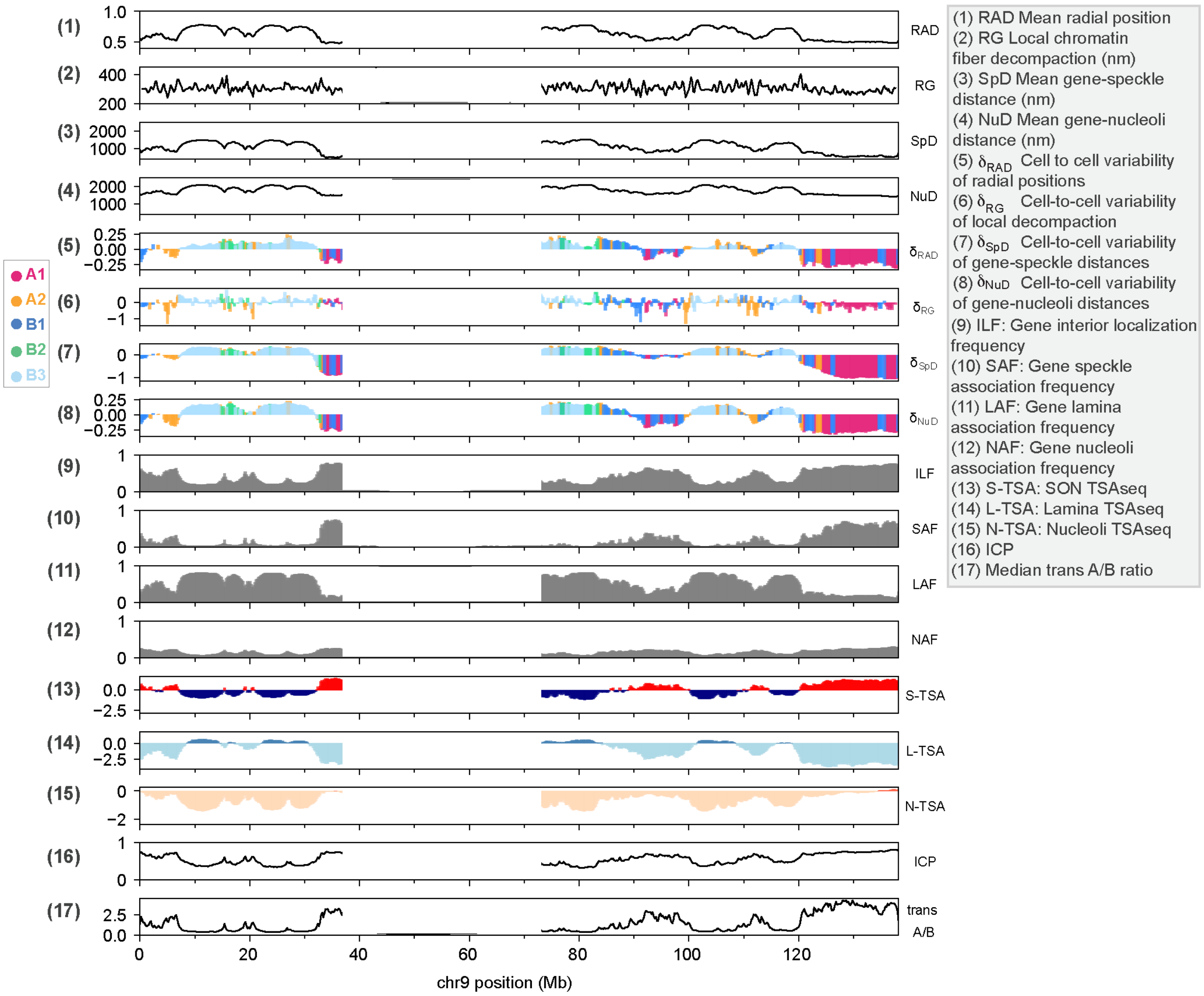


**Fig. S12.** Structure feature profiles for chromosome 9.

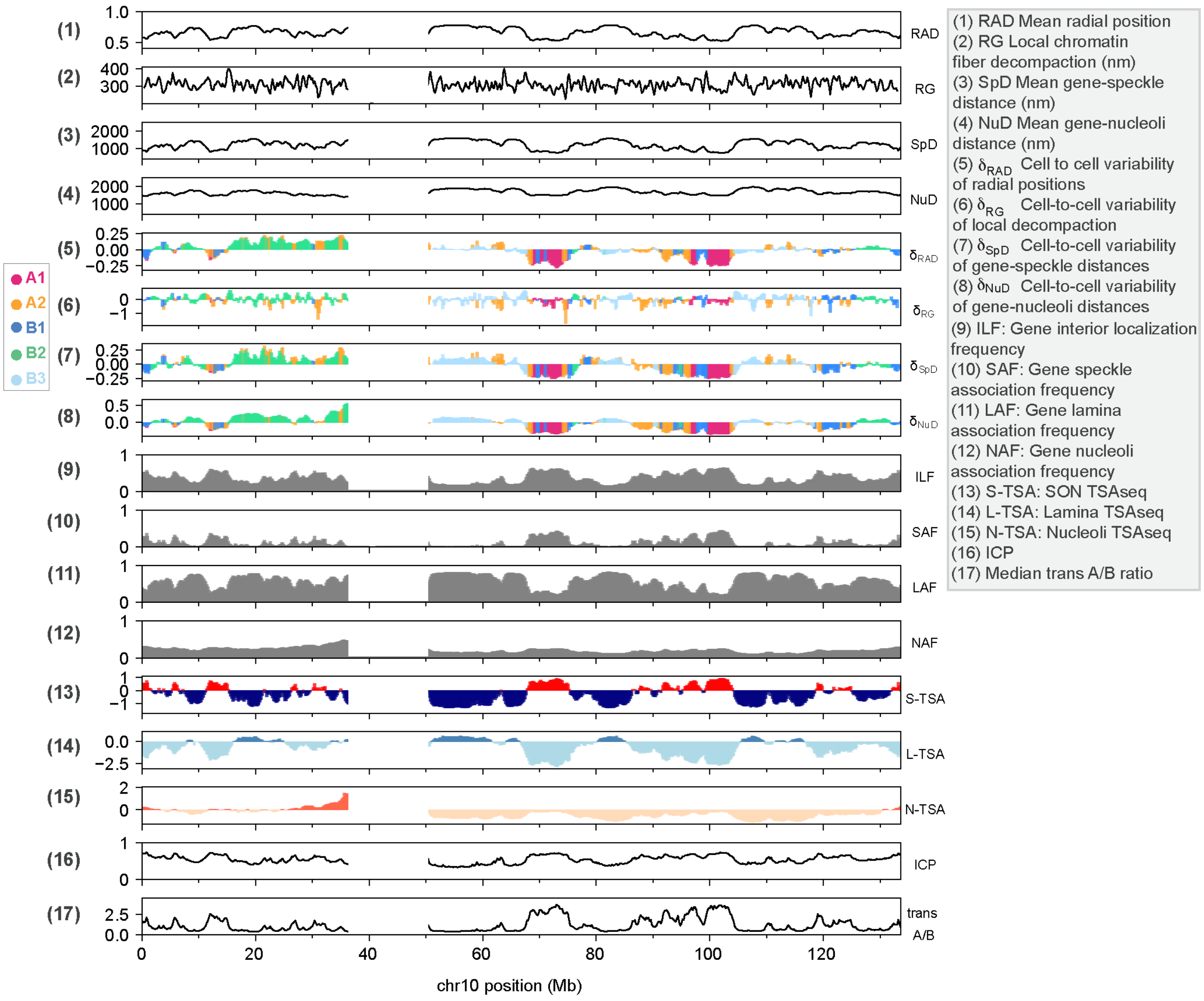


**Fig. S13.** Structure feature profiles for chromosome 10.

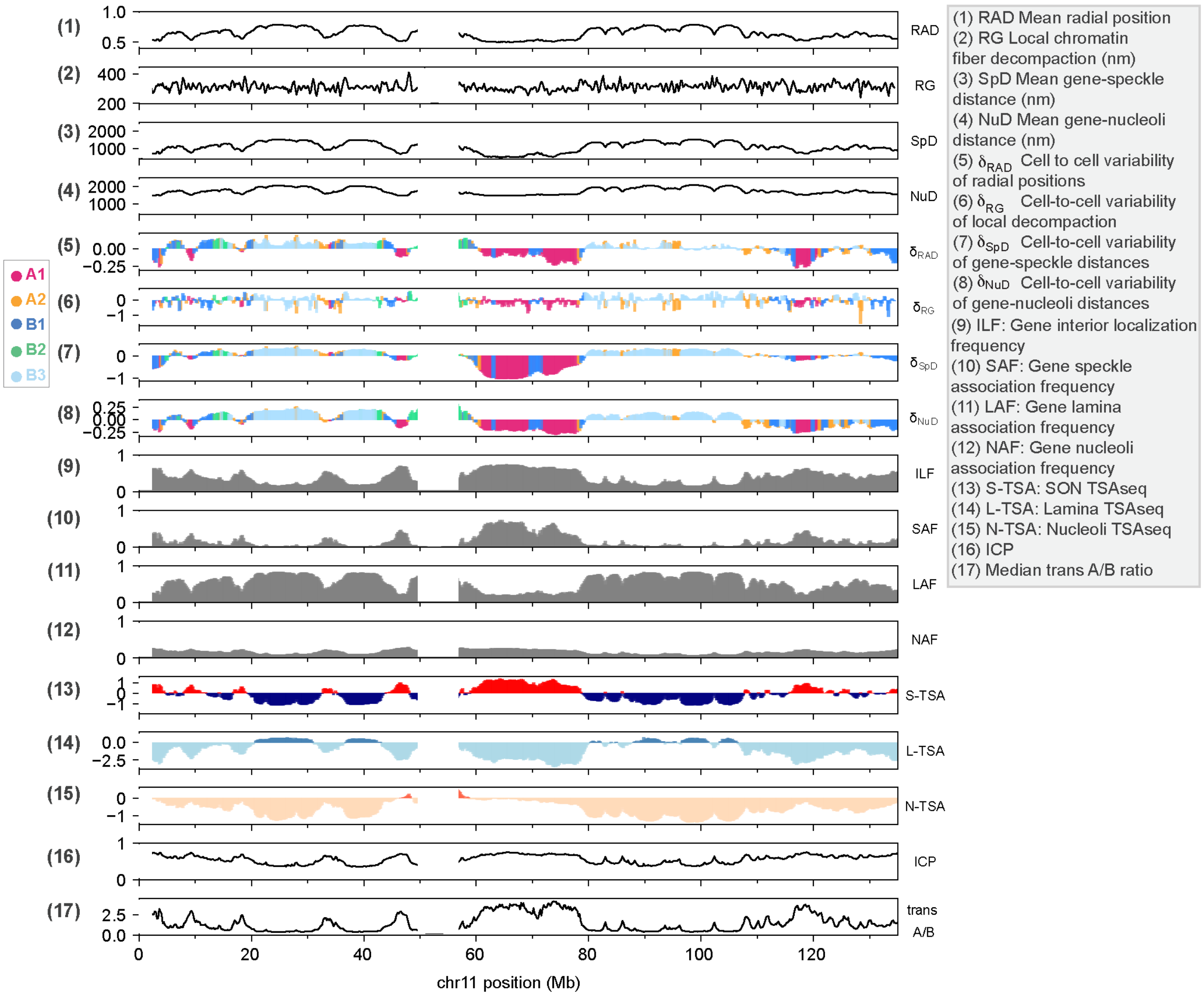


**Fig. S14.** Structure feature profiles for chromosome 11.

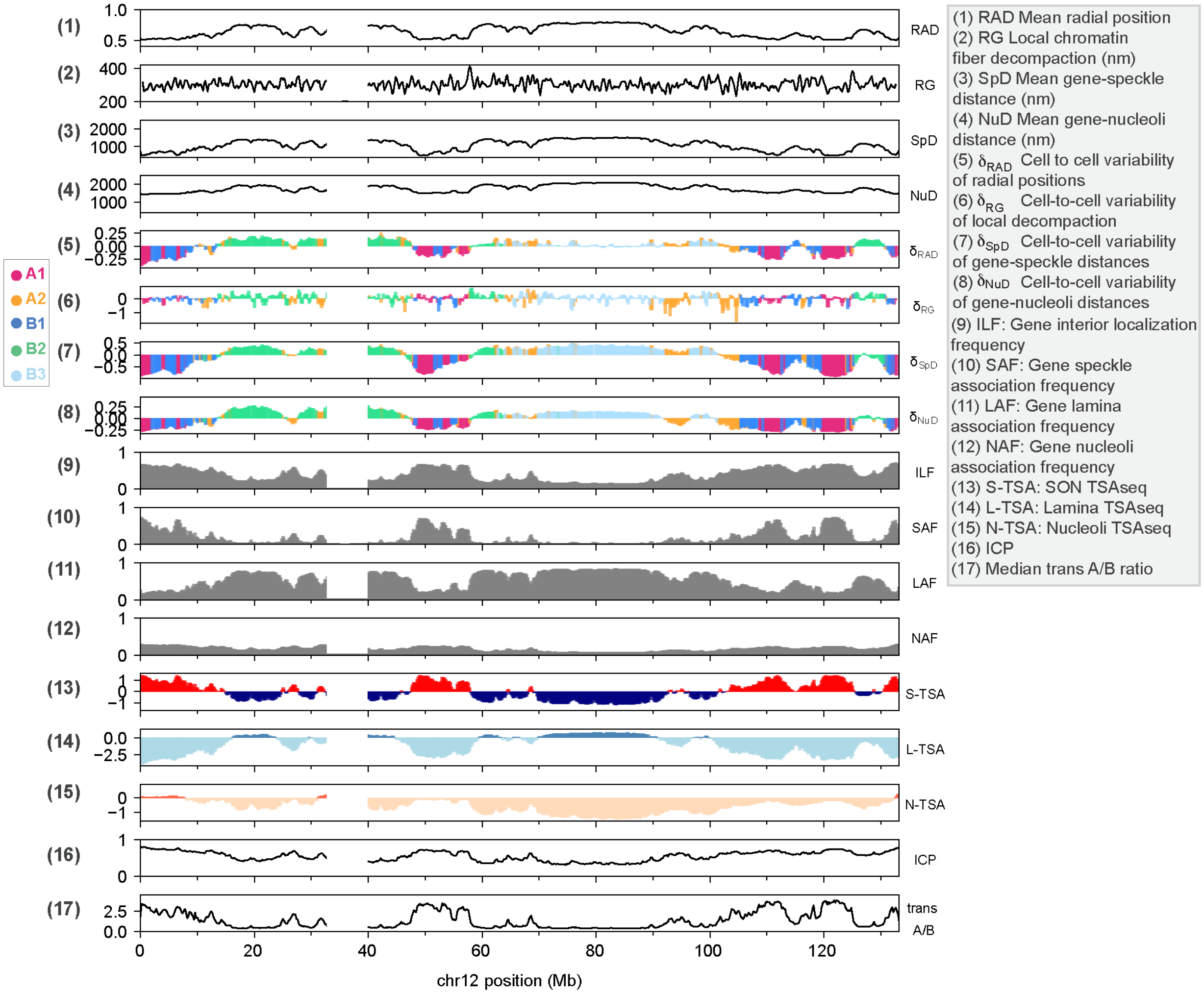


**Fig. S15.** Structure feature profiles for chromosome 12.

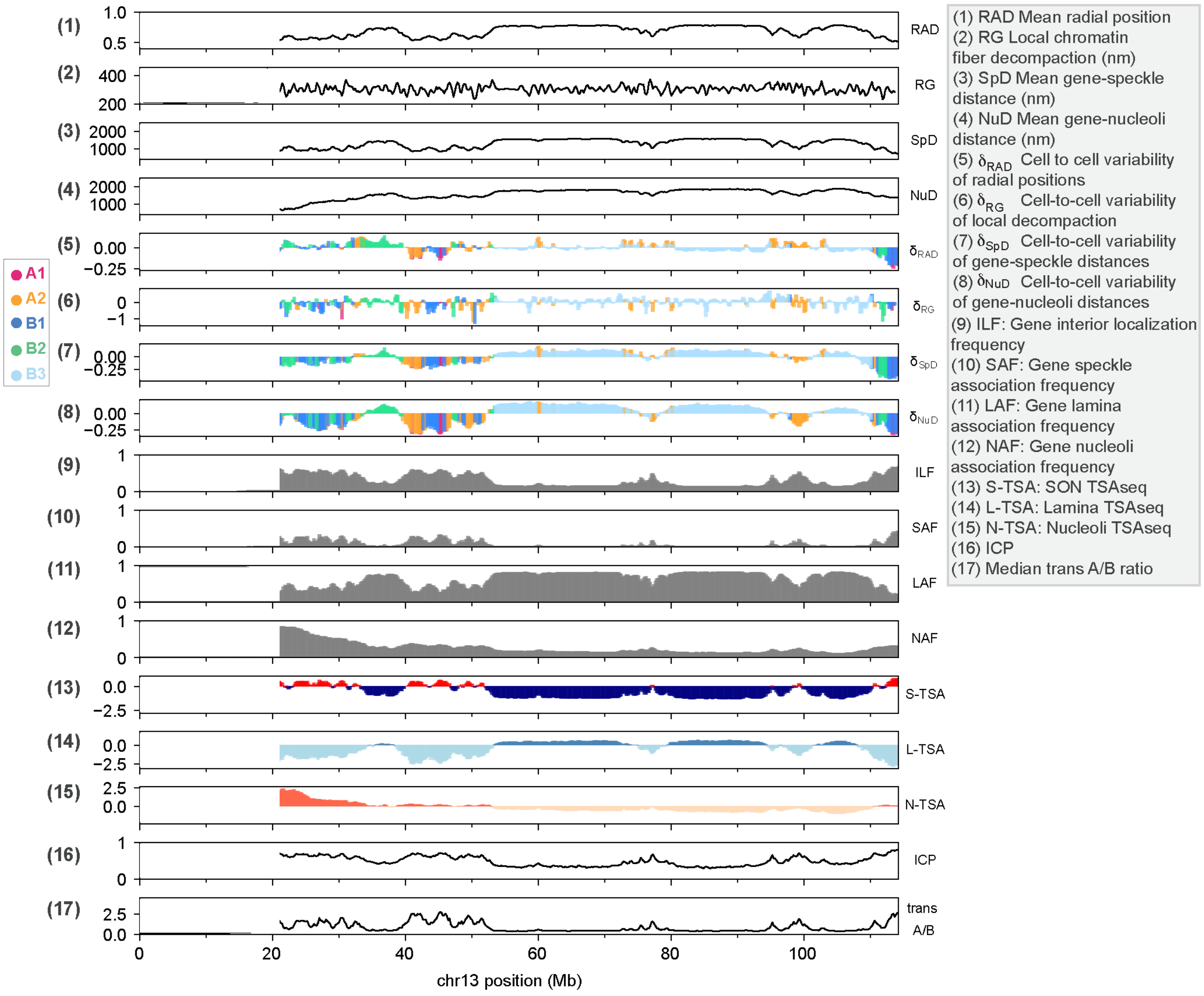


**Fig. S16.** Structure feature profiles for chromosome 13.

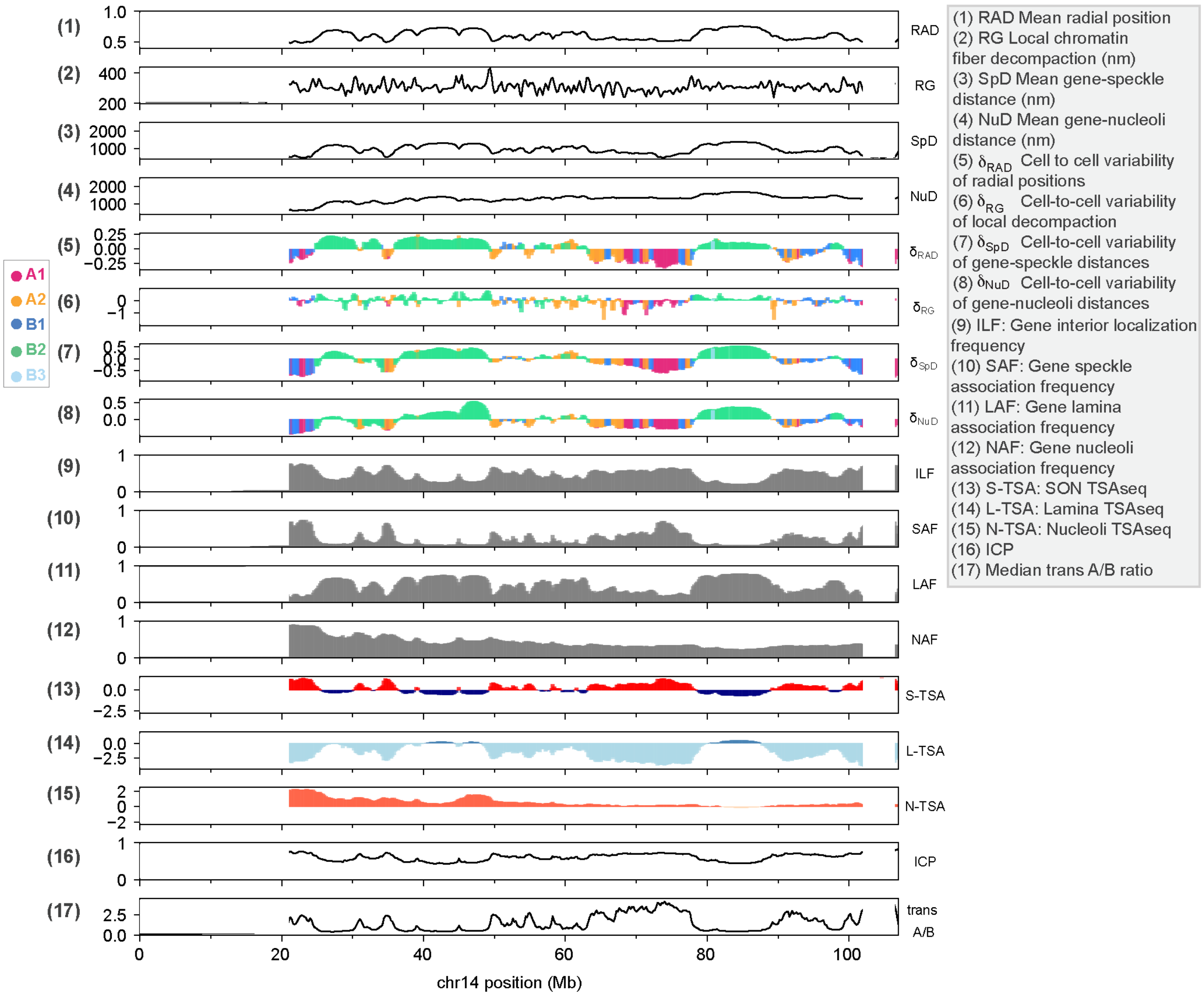


**Fig. S17.** Structure feature profiles for chromosome 14.

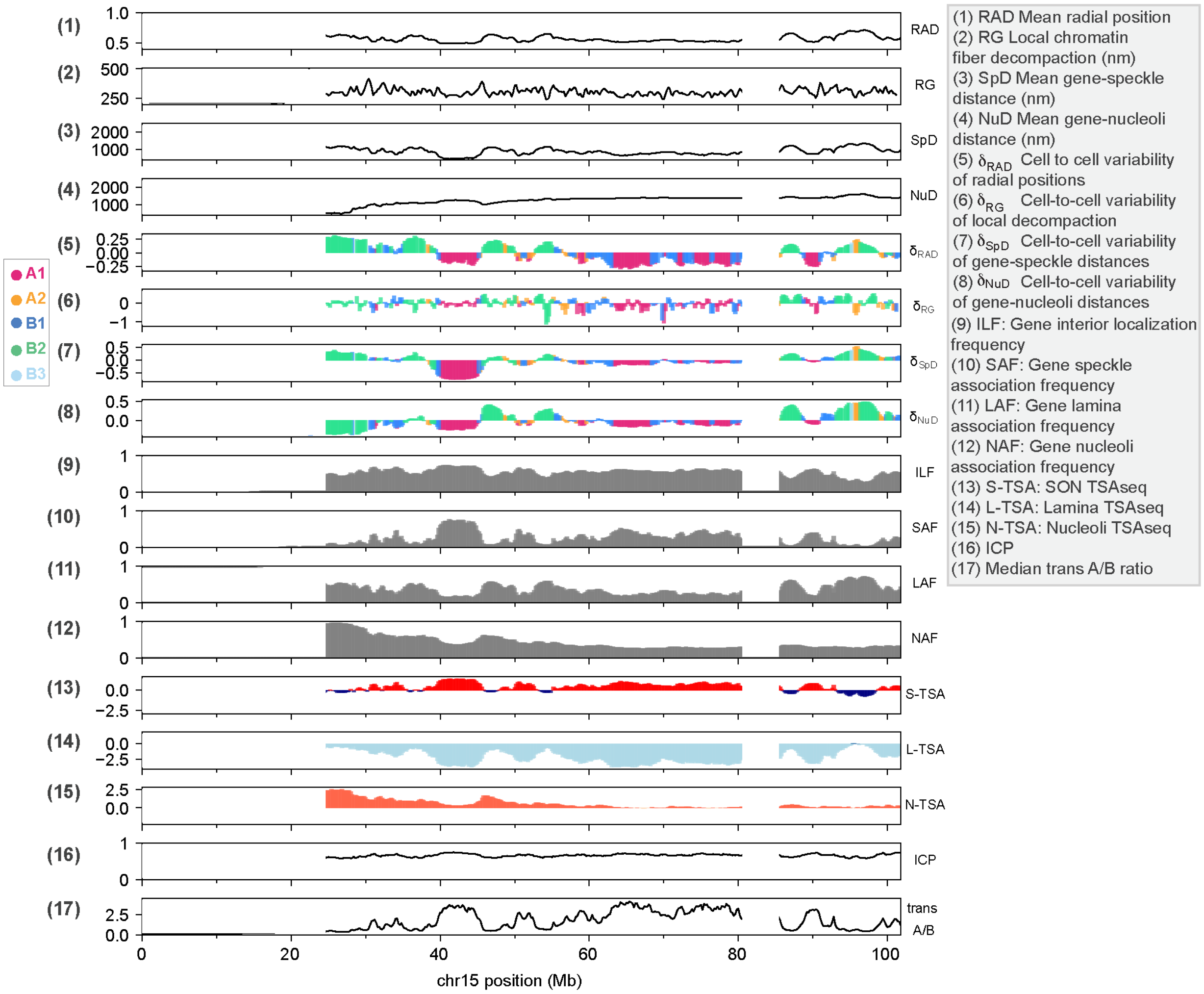


**Fig. S18.** Structure feature profiles for chromosome 15.

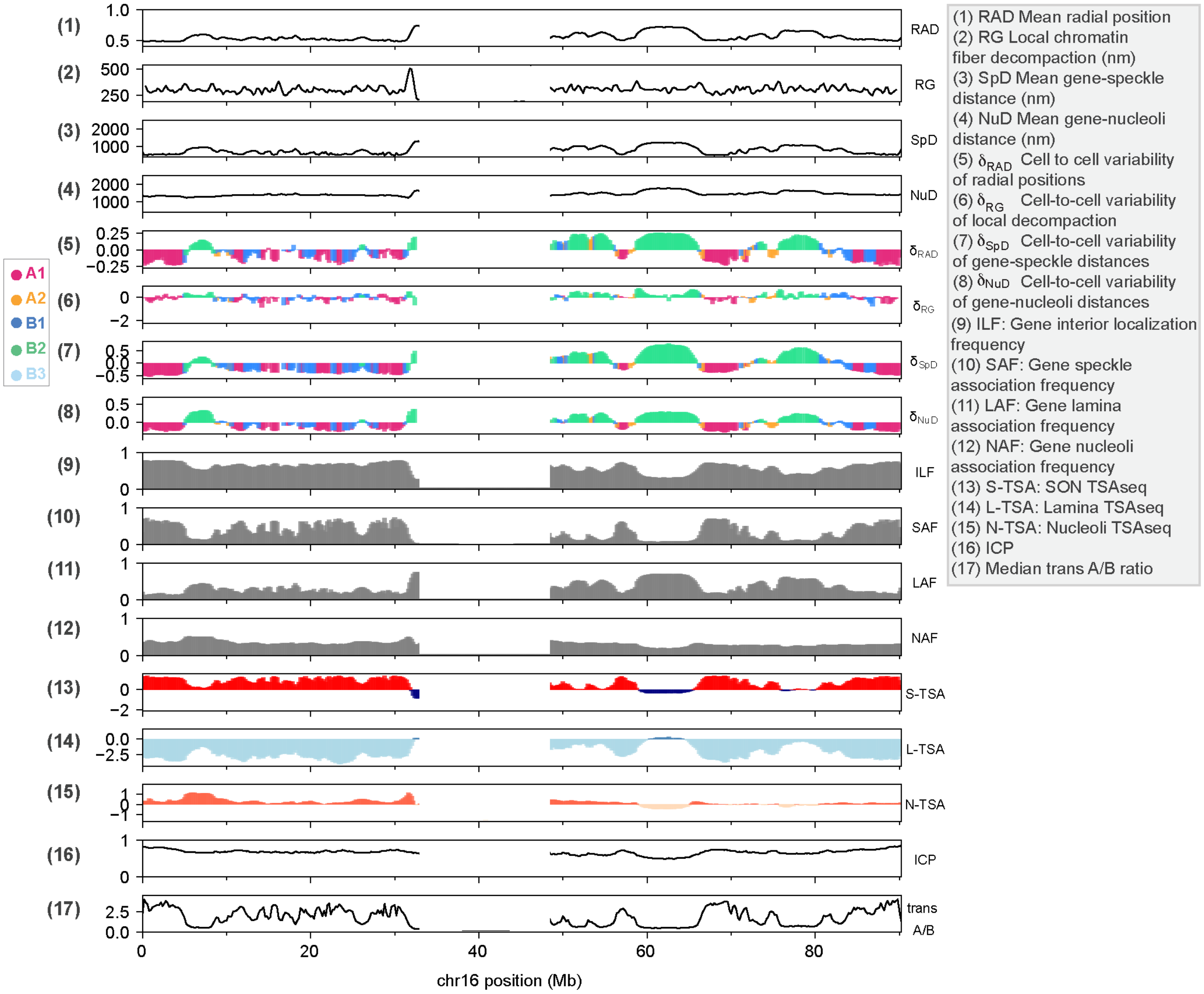


**Fig. S19.** Structure feature profiles for chromosome 16.

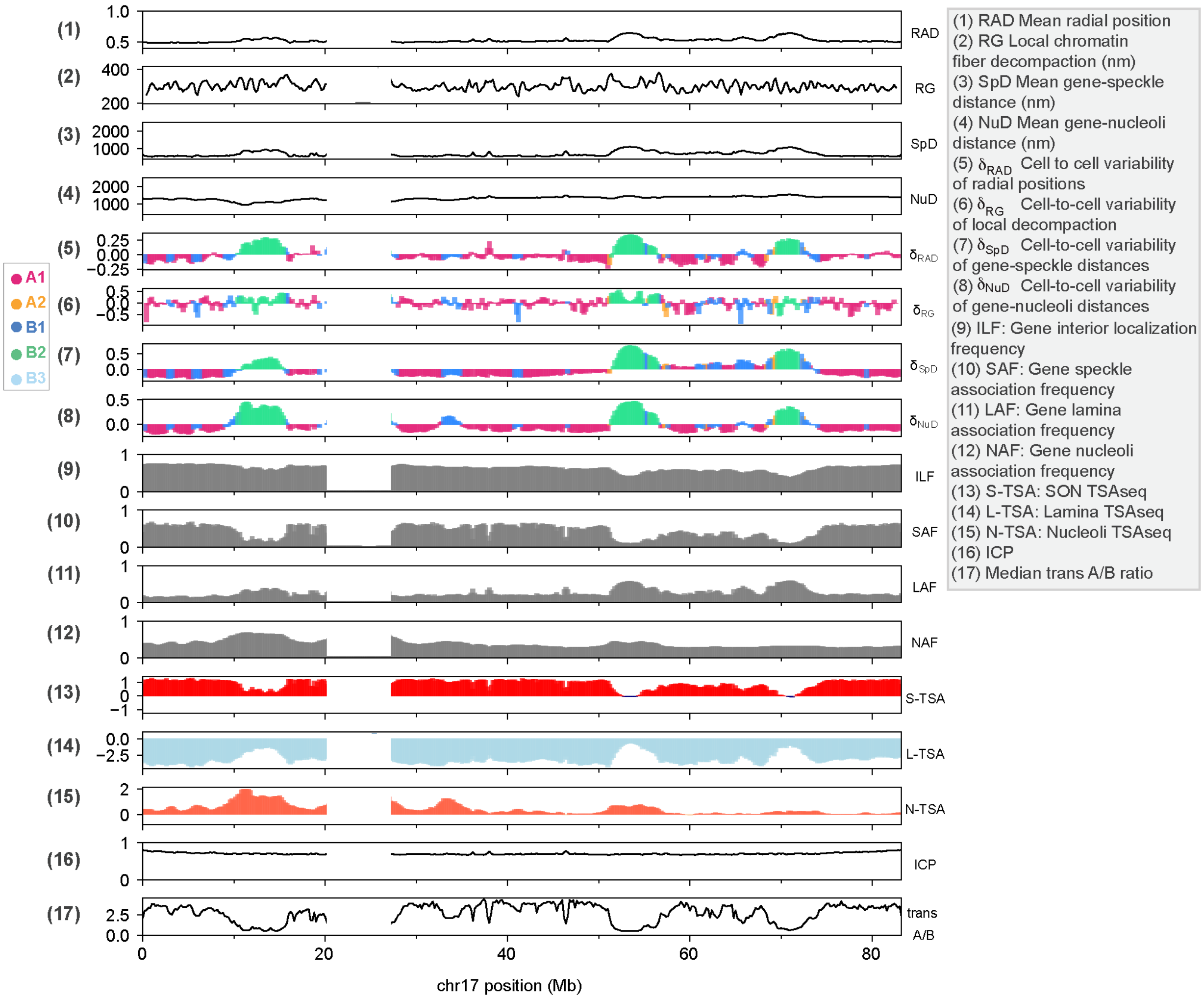


**Fig. S20.** Structure feature profiles for chromosome 17.

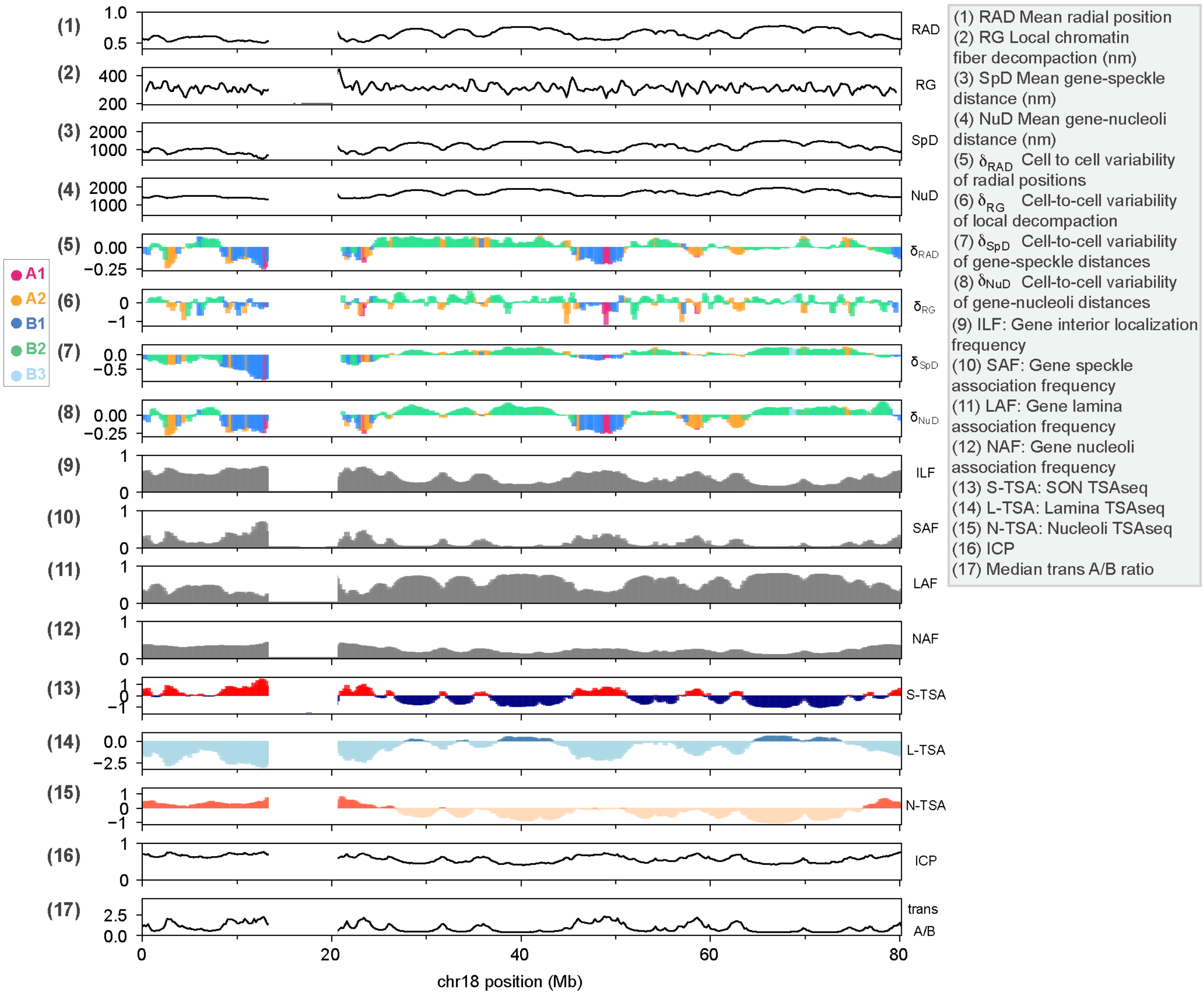


**Fig. S21.** Structure feature profiles for chromosome 18.

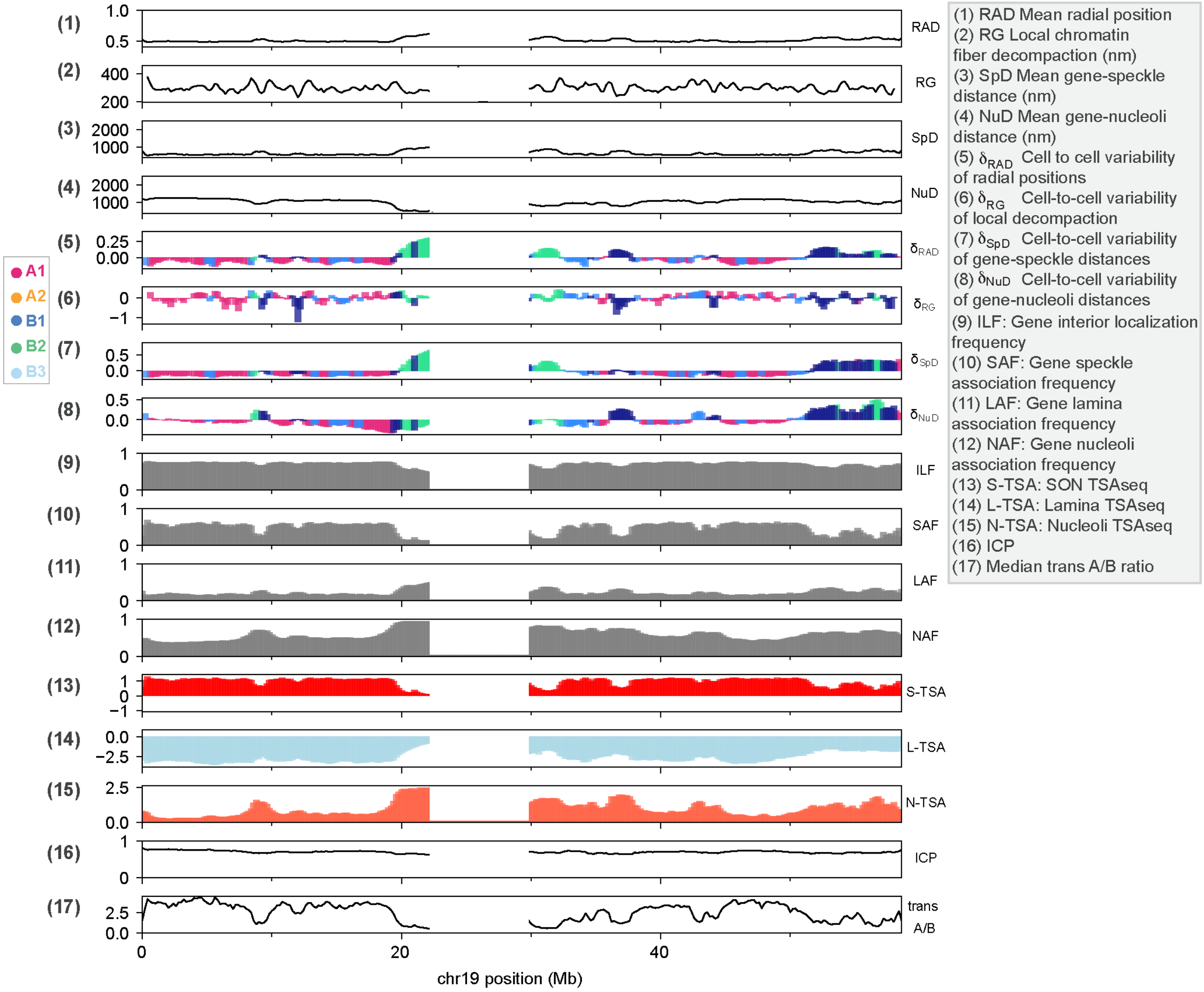


**Fig. S22.** Structure feature profiles for chromosome 19.

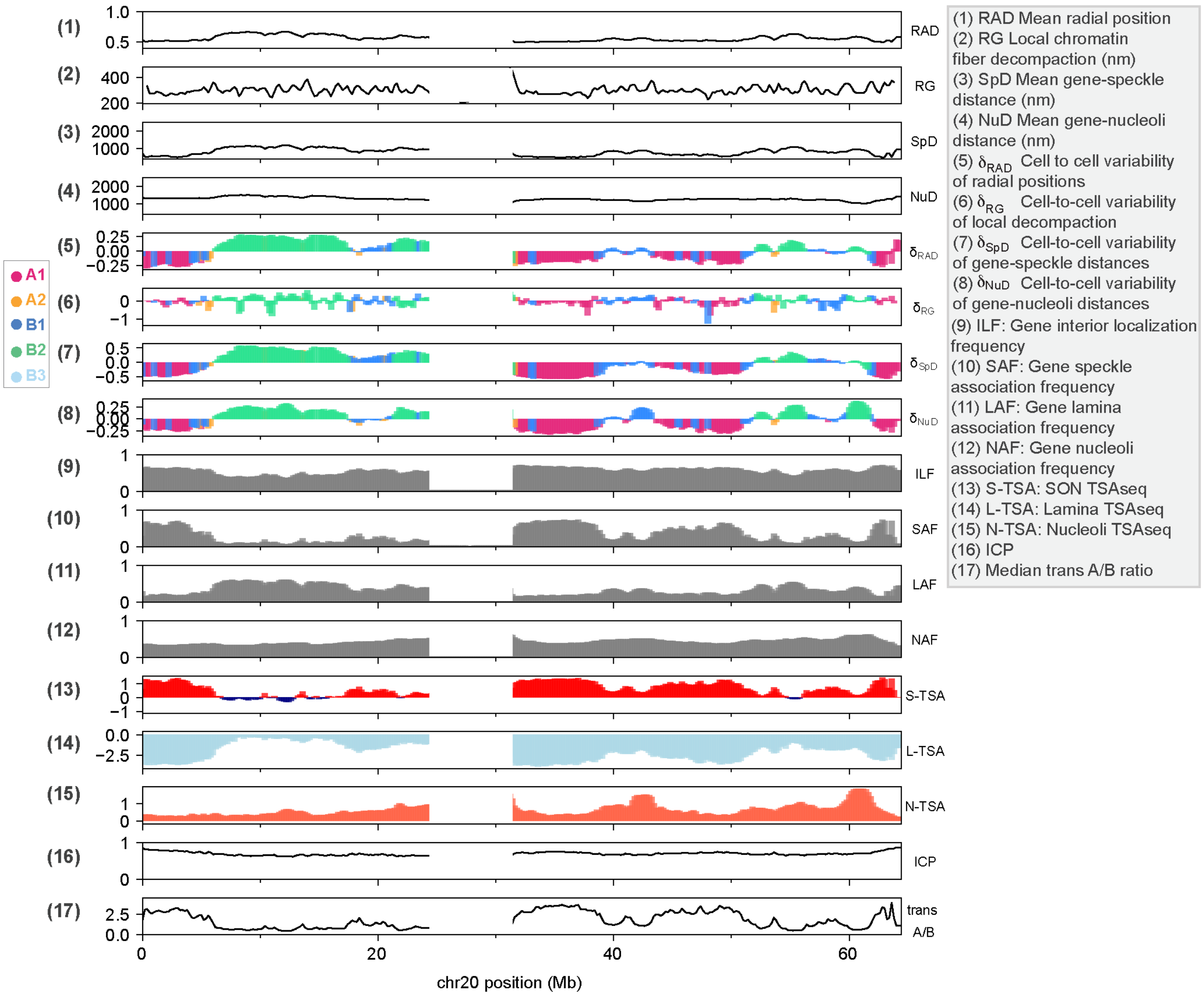


**Fig. S23.** Structure feature profiles for chromosome 20.

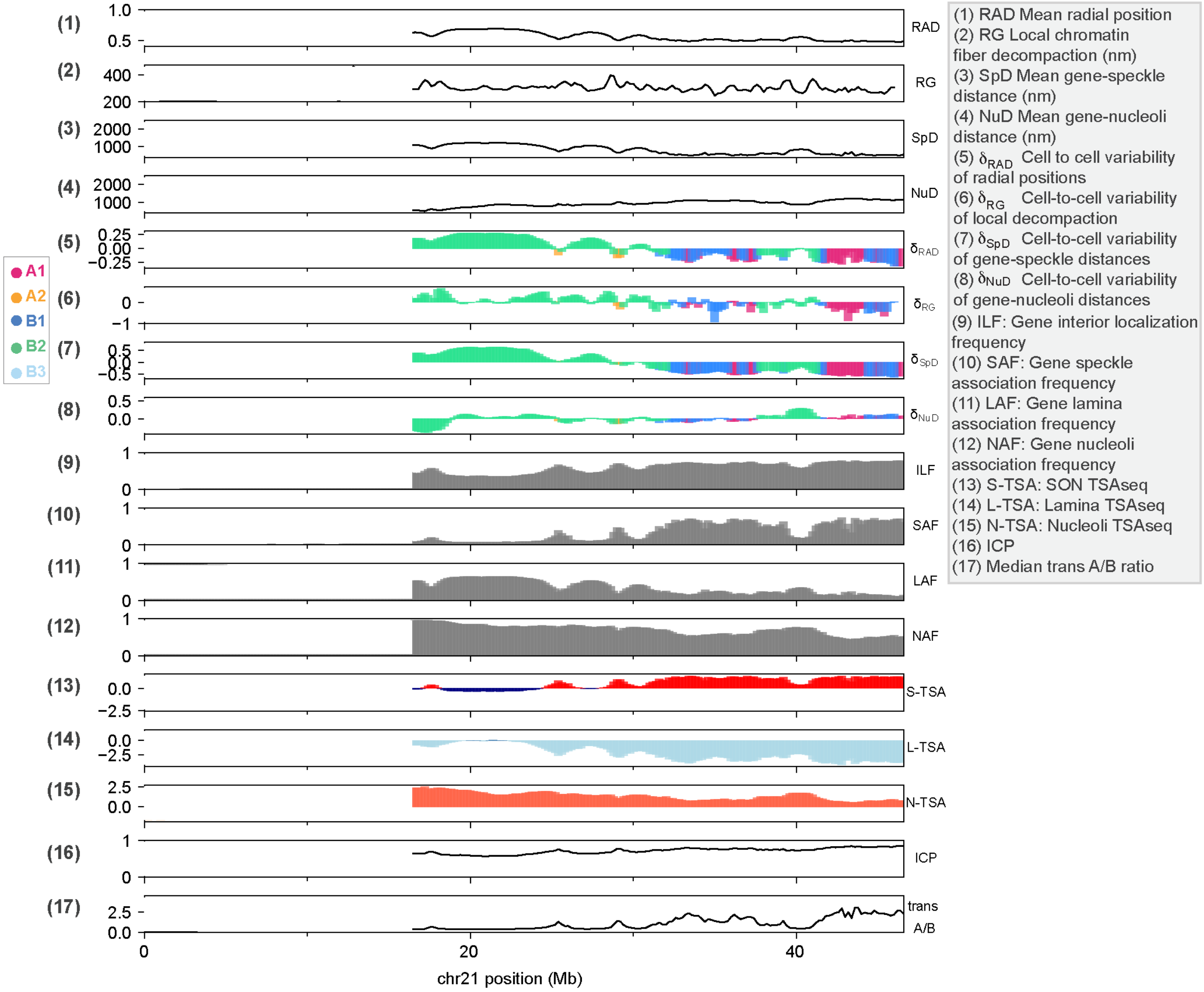


**Fig. S24.** Structure feature profiles for chromosome 21.

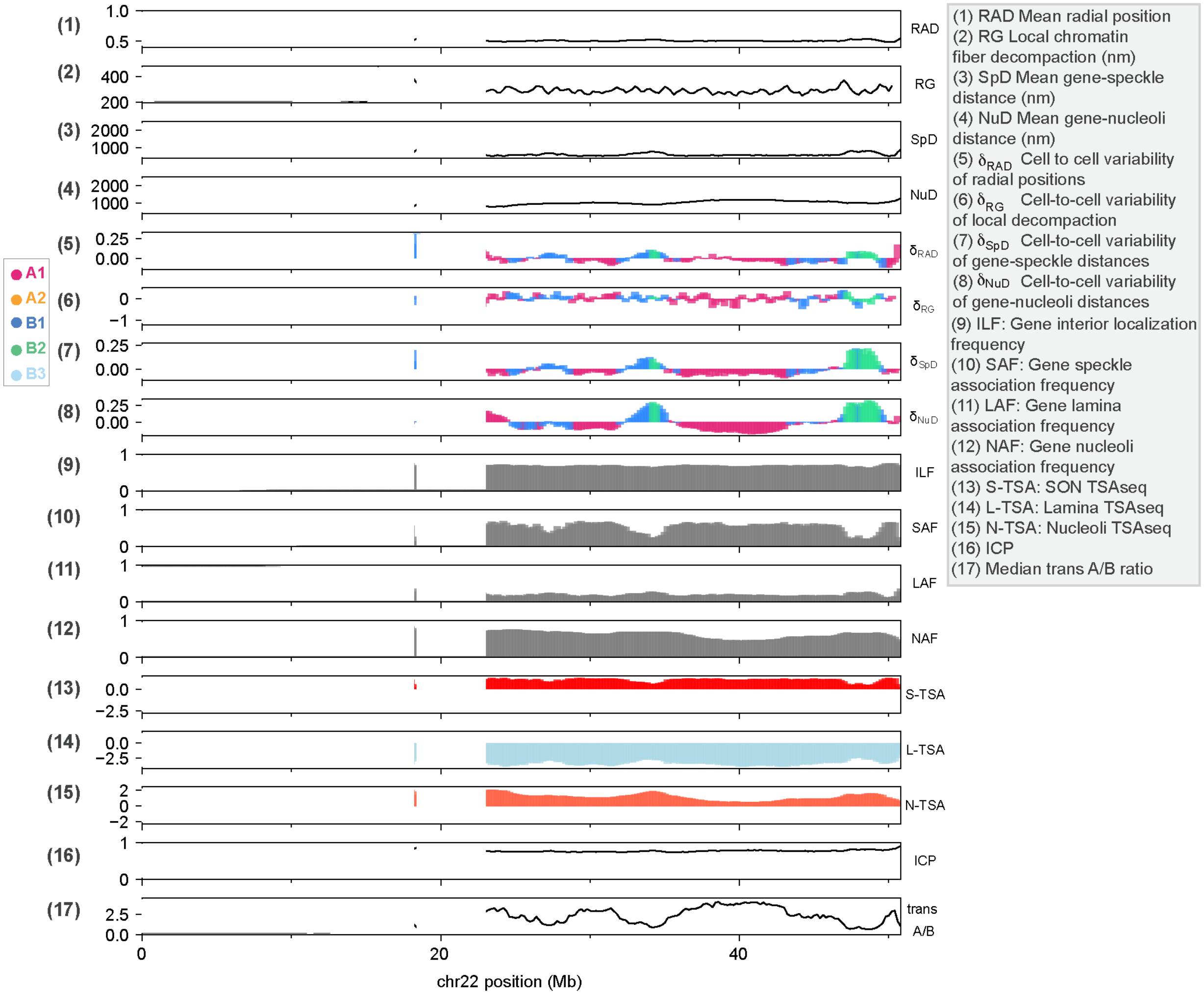


**Fig. S25.** Structure feature profiles for chromosome 22.

**Table S1.** Pearson and stratum adjusted correlation coefficients (SCC)[1] between the input and output Hi-C matrices for each chromosome. For SCC calculation, the smoothing parameter and the upper bound of the genomic distance for interacting loci were set to 0 and 50 Mb, respectively.

| Chromosome | Pearson's R | SCC |
|---|---|---|
| chr1 | 0.98 | 0.83 |
| chr2 | 0.99 | 0.90 |
| chr3 | 0.99 | 0.91 |
| chr4 | 1.00 | 0.92 |
| chr5 | 0.99 | 0.90 |
| chr6 | 0.99 | 0.88 |
| chr7 | 0.99 | 0.90 |
| chr8 | 1.00 | 0.90 |
| chr9 | 0.98 | 0.84 |
| chr10 | 0.99 | 0.86 |
| chr11 | 0.99 | 0.87 |
| chr12 | 0.99 | 0.85 |
| chr13 | 0.99 | 0.94 |
| chr14 | 0.98 | 0.88 |
| chr15 | 0.97 | 0.84 |
| chr16 | 0.98 | 0.81 |
| chr17 | 0.96 | 0.78 |
| chr18 | 0.99 | 0.87 |
| chr19 | 0.98 | 0.81 |
| chr20 | 0.97 | 0.79 |
| chr21 | 0.98 | 0.88 |
| chr22 | 0.97 | 0.88 |
| chrX | 1.00 | 0.91 |

**Table S2.** Experimental data used in our analyses.

| Data | Accession Code |
|---|---|
| Hi-C[2] | GEO: GSE63525 |
| SON TSA-seq[3] | GEO: GSE81553 |
| LaminB1 TSA-seq[3] | GEO: GSE81553 |
| Single cell Lamina DamID[4] | GEO: GSE56465 |
| LaminB1 pA-DamID[5] | 4DN: 4DNFIGL8MCSJ |
| GRO-seq[6] | GEO: GSM1480326 |
| GP-seq[7] | GEO: GSE135882 |
| Repli-seq[8] | GEO: GSM923451 |
| ChIP-seq (Histone modif.)[9,10] | ENCODE: ENCFF313LYI, ENCFF171MDW, ENCFF776DPQ, ENCFF309OEW, ENCFF028KBY, ENCFF601YET, ENCFF831ZHL, ENCFF039HDL, ENCFF340JIF, ENCFF803DJF, ENCFF683HCZ |
| scRNA-seq[11] | GEO: GSM3596321 |
| Superresolution imaging[12] | https://zenodo.org/record/3928890 |
| Subcompartments[2] | GEO: GSE63525 |
| Compartments[2] | 4DN: 4DNFILYQ1PAY |
| LADs | obtained from refs.[4,7] |
| Enhancers/Superenhancers | obtained from ref.[13] |

### Preprocessing Hi-C data

We used *in situ* Hi-C datasets from human lymphoblastoid cell line GM12878 (reference genome hg38)[2]. For this dataset, the raw contact map was downloaded from Gene Expression Omnibus (GEO) under the accession number GSE63525. Similar to the protocol by ref.[14], low bin sequence coverage 3% regions were discarded during the normalization process. For data normalization, we adopted the same KR normalization method used in[2], leading to a normalized contact frequency matrix $\boldsymbol{F} = (f_{ij})_{K\times K}$ at 20 kb resolution with $K = 151{,}561$ bins. We then generated a probability matrix at 200 kb level as our input for our algorithm using the following approach:

We converted the contact frequencies in the 20-kb matrix to contact probabilities by scaling the frequencies by a normalization factor, $f^{max}$, which is chosen to represent the contact frequency value at which two domains have a 100% probability to form a contact. The 20-kb contact probability matrix $\boldsymbol{P} = (p_{ij})_{K\times K}$ was calculated as $p_{ij} = \min\left(\frac{f_{ij}}{f^{max}}, 1\right)$, where $p_{ij}$ and $f_{ij}$ are the contact probability and frequency values, respectively. We set the value of $f^{max}$ so that the average contact probability sum of a region is ~24, which, based on our experience, is the average number of contacts a domain has at saturation (where no more contact restraints can be satisfied, given the contact distance cutoff)[15].

We then defined a mapping $b(i)$ as the set of all 20-kb bins in matrix $\boldsymbol{P}$ that belongs to 200-kb region $i$. Then the domain-level matrix $A = (a_{ij})_{N\times N}$ was calculated as:

$$a_{ij} = mean(\ top10\% < \{p_{\alpha\beta}:\ \alpha \in b(i), \beta \in b(j)\} > )$$

In the case that some contacts were drastically higher than the surrounding contacts, these contacts were identified as outliers by $\{p: p > \mu + 1.5IQR\ \}$, where $p \in \{p_{\alpha\beta}: \alpha \in b(i), \beta \in b(j)\}$ and $\mu = mean\{p_{\alpha\beta}: \alpha \in b(i), \beta \in b(j)\}$. The IQR refers to the interquartile range of $\{p_{\alpha\beta}\}$. These outliers were excluded from the calculation.

After obtaining the contact probability matrix at 200-kb resolution, we identified bins that have spurious inter-chromosomal interaction probabilities (higher than 0.2), and removed the corresponding bins in the 20-kb raw matrix, and repeated the KR normalization, and regenerated the 200-kb contact probability matrix where no inter-chromosomal contact probability is higher than 0.2 following the same procedure explained above. Finally, we set contact probabilities between the consecutive domains as well as between domains up to 1 Mb distance in the gap regions to 1 in order to maintain the chain integrity.

### Iterative refinement of restraint assignment

During the optimization procedure, it is possible that excess contacts can lead to compact structures that cannot be further relaxed easily. A heuristic method has been put in place to compensate for such an effect, by allowing us to assess the actual portion of expected contacts that are available for allocation, which then affects the way the activation distance $d_{act}^{IJ}$ is computed in the Hi-C assignment steps (A). The empirical procedure relies on the predicament that when a population expresses more contacts than it should, we reduce the assignment probability; a lower probability (with fewer restraints) should be equally effective.

Assume the expected number of contacts $Sp_{ij}^{input}$ is the sum of a number of effective contacts that are actually imposed, $N_{eff}$, and a number of incidental contacts (that are also expressed but not imposed), $N_{inc}$:

$$Sp_{ij}^{input} = N_{eff} + N_{inc} = N_{eff} + \eta(S - N_{eff})$$

The number of incidental contacts is expressed as a fraction of the number of non-applied (non-enforced) contacts. The latter term can originate from cooperative effects, which automatically bring loci closer without an explicit bonding term operating. We can solve for probability $p_0 = N_{eff}/S$: $p_0 = \frac{p^{input} - \eta}{1-\eta}$. This is an effective probability which controls the number of restraints to be enforced. The estimate for the scalar $\eta$ is calculated as follows.

Let us compare the factual number of contacts in the population (expressed by the tensor $A_{ij}^{X} = \sum_{s=1}^{S} \boldsymbol{W}_{ijs}^{(k-1),\mathrm{X}}$) with the predicted number of contacts from the previous assignment step ($A_{ij}^{assig} = \sum_{s=1}^{S} \boldsymbol{W}_{ijs}^{(k-1),\mathrm{assig}}$):

$$\sum_{s=1}^{S} \boldsymbol{W}_{ijs}^{(k-1),\mathrm{X}} = \sum_{s=1}^{S} \boldsymbol{W}_{ijs}^{(k-1),\mathrm{assig}} + \left(S - \sum_{s-1}^{S} \boldsymbol{W}_{ijs}^{(k-1),\mathrm{assig}}\right)\eta_{ij}$$

We can solve for $\eta_{ij}$:

$$\eta_{ij} = \frac{\sum_{s=1}^{S} \boldsymbol{W}_{ijs}^{(k-1),\mathrm{X}} - \sum_{s=1}^{S} \boldsymbol{W}_{ijs}^{(k-1),\mathrm{X}}}{S - \sum_{s=1}^{S} \boldsymbol{W}_{ijs}^{(k-1),\mathrm{assig}}} = \frac{A_{ij}^{X} - A_{ij}^{assig}}{1 - A_{ij}^{assig}},$$

which can then be plugged into the equation 1 to find a corrected assignment probability $p_0 = \frac{p^{input} - \eta}{1-\eta}$, which is then used to update the activation distance $d_{ij}^{act}$. Please note that the correction is only implemented if there is a contact excess.

### Mapping experimental data to 200kb models

The list of experimental data used in our analyses is reported in Table S2. Each of the experimental data is in a different resolution; therefore, we performed several mapping methods to bring the data to the models' resolution (200-kb). We also lifted any hg19 data to hg38 by using the liftOver tool from UCSC[16]. The mapping methods for different data are listed below.

- **scRNA-seq**

Each gene in the scRNA-seq data set[11] was mapped to the 200-kb chromatin region in our models with the largest sequence overlap. Each gene was mapped to a single chromatin region, and if multiple genes were mapped to a single chromatin region, the gene with the highest total expression level was selected. After the mapping process, a total of 8,920 genes (7,090 of them with non-zero total expression levels) were mapped to 8,920 200-kb chromatin regions in our models. In our analyses, scRNA-seq data for the GM12878 cells in G1 phase were used.

- **SON and LaminB1 TSA-seq**

Each region in the TSA-seq data[3] was mapped to 200-kb regions with an overlap of 50% or higher. After mapping, each 200-kb region had multiple TSA-seq regions; therefore the signals mapped to each 200-kb region were averaged (we first took the inverse log2 of the signals, then averaged and took the log2 of the average value.). Transcription hot zones were also downloaded from ref[3].

- **LaminB1 scDamID and pA-DamID**

Each region in the pA-DamID data[5] was mapped to 200-kb regions with an overlap of 50% or higher. After mapping, each 200-kb region had multiple DamID regions; therefore the signals mapped to each 200-kb region were averaged (we first took the inverse log2 of the signals, then averaged and took the log2 of the average value.).

100-kb single-cell LaminaDamID lamina contact frequencies[4] were mapped to 200kb and averaged over 200-kb regions.

- **GP-seq**

First, 100-kb rescaled GP-seq scores were averaged over 4 replicate experiments (2 HindII, 2 MboI experiments) for each 100-kb region[7]. The values were then mapped to 200-kb and averaged over 200-kb regions.

- **GRO-seq**

GRO-seq read counts[6] at plus and minus strands were summed up for each 200-kb region.

- **Repli-seq**

We first calculated the average percentage normalized signals (percentage of normalized tag densities over all phases) in each 200-kb region for each phase (G1b, S1, S2, S3, S4, G2)[8]. We then assigned each 200-kb region with the cell phase with the highest average percentage. For example, if a 200-kb region showed the average normalized percentage of tag densities for G1b, S1, S2, S3, S4, and G2 as 10%, 45%, 20%, 10%, 5%, and 0%, then the region was assigned to S1 phase.

- **Superresolution imaging**

For the recently publised superresolution imaging data[12], we used the provided datasets at https://zenodo.org/record/3928890. 1041 imaged loci were first mapped to 200-kb resolution. All loci were either mapped to one or two 200-kb regions depending on their overlaps. If a locus was mapped to two regions, the features calculated from the models were averaged over these two regions in the analyses.

- **ChIP-seq (Histone marks)**

The signals and peaks for H3K36me3, H3K27me3, H3K9ac, H3K9me3, H3K27ac, H3K4me1, H3K4me2, H3K4me3, H3K79me2, H4K20me1, and H2AFZ histone modifications were downloaded from ENCODE[9,10]. Each region in the ChIP-seq data was mapped to 200-kb regions with an overlap of 50% or higher. After mapping, each 200-kb region had multiple ChIP-seq regions; therefore the signals mapped to each 200-kb region were averaged (we first took the inverse log2 of the signals, then averaged and took the log2 of the average value.), and the number of peaks mapped to each region were counted.

- **Subcompartments**

The definition of subcompartment states was retrieved from ref[2] which are at 100-kb resolution. For each 200-kb region of chromatin, we first mapped the subcompartment states and calculated their proportion. The state of the 200-kb region was then set to the majority of its constituent state (50% or more). If there was no majority constituent state, we then assigned the region as NA and discarded them from further subcompartment-related analyses.

- **Compartments**

The definition of compartment states were retrieved from the 4DN data portal (Table S2) and ref[2] which are at 250-kb resolution. We mapped each compartment state to a 200-kb region with the maximum overlap. If there was no mapped state to a 200-kb region, then we assigned it as NA and discarded them from further compartment-related analyses.

- **LADs**

The definitions of LADs (constitutive and facultative LADs and inter-LADs) were retrieved from refs.[4,7]. LAD assignments were mapped to overlapping 200-kb regions. After mapping, if a 200-kb region had only one mapped LAD state, we assigned that state to the region. However, if the region had multiple mapped LAD states, we assigned the region as NA and discarded them from further LAD-related analysis. This mapping procedure resulted in 1304, 495, 1010, 1904 regions with cLAD, fLAD, ciLAD, and fiLAD assignments, respectively.

- **Enhancers/Superenhancers**

The definitions of enhancers (EN) and superenhancers (SEN) were retrieved from ref[13]. Each EN and SEN region was mapped to 200-kb regions with an overlap of 50% or higher. After mapping, we assigned regions as EN/SEN if they overlapped with one or more EN/SEN peaks. We assigned regions as NA if they did not overlap with any EN/SEN peaks and discarded them from further EN/SEN-related analysis.

### 3D DNA FISH *Experiments*

We carried out a set of 3-color FISH experiment where all probes were on chromosome 6: RP11-945M14 (306,712 – 532,406), RP11-1076L22 (130,076,074 – 130,287,007), and RP11-111J1 (87,193,201 – 87,351,481). For any particular chromosome domains (regions), multiple BAC clones were chosen and synthesized by Empire Genomics and tested individually for their specificity. The experiment was performed following the previous protocols[17,18]. GM12878 cells were cultured in a DMEM medium supplemented with 15% FBS, glutamine and penicillin/streptomycin as suggested by ENCODE. Two days before the experiment, 22mm x 22mm coverslips were cleaned and coated with L-poly-lysine (1mg/ml) at room temperature for 1-2 hours, and dried in a tissue culture hood after a brief rinse with sterile MilliQ water. On the day of the experiment, 10 million GM12878 cells were harvested by centrifugation at 100g for 10 minutes, resuspended in fresh culture medium ($3x10^6$ cells/ml), and seeded evenly on the coverslip in a 6-well tissue culture plate. After incubating at 37°C for one hour and briefly washing with PBS, the cells (on coverslips) were fixed with 4% freshly made paraformaldehyde (in 0.4x PBS) at room temperature for 10 minutes. The cell membrane was permeabilized firstly with 0.5% triton X100/1xPBS at room temperature for 20 minutes, and then through 4-5 freeze-and-thaw cycles (by dipping in liquid nitrogen and then thawing in room temperature) in the next day after pretreatment overnight with 20% glycerol/1xPBS and also before each dip. To facilitate access of the FISH probe to the chromatin DNA, the samples were each washed twice with 0.05% triton X100/1xPBS for five minutes, and then treated with 0.1N HCl at room temperature for 5-10 minutes to remove basic nuclear proteins. The HCl was removed from the sample followed by two washes with 0.05% triton X100/PBS and one wash with 2x SSC (diluted from 20xSSC: 3M NaCl, 0.3M sodium citrate, pH 7.0) for 5-10 minutes each wash. The coverslips were then stored in 50% formamide/2x SSC at 4°C and were ready for the next step (good for two days to two months). The denaturation and hybridization steps were performed according to the protocols suggested by the manufacturer. (https://www.empiregenomics.com/files/store/products/FISH_probes/FISH_Protocol.pdf). The coverslips were brought to room temperature for 24 hours in advance before denaturing. On the day of experiment, fresh 70% formamide/2x SSC was prepared and pre-warmed at 73°C for 30 minutes. Cells on the coverslips were denatured in this solution (73°C) for five minutes, and dried through sequentially dipping into 70%, 85% and 100% ethanol one minute each at room temperature, and finally through evaporation at 45°C for 20 minutes. FISH probes were denatured similarly in 70% formamide/2x SSC for 5 minutes at 73°C and then quickly cooled down on ice. After incubating at 37°C for 10-20 minutes, three probes (150 ng each) for either targeted regions or for the three control regions were mixed thoroughly with 18 μl hybridization buffer (provided by the manufacturer), and applied evenly with the sample on a microscope slide. Hybridization of FISH probes with the samples occurred in a humidified chamber containing a paper towel soaked with 50% fomamide/ 2x SSC in dark at 37°C for 18-20 hours. Unbound FISH probes were removed by a series of washes, three times with 0.3% NP-40/0.4x SSC at 73°C for two minutes, each followed by a wash with 0.1% NP40/2x SSC at room temperature for one minute. After air-drying for five minutes in dark, the coverslips were mounted on microscope slide with 10 μl DAPI mounting solution and kept in dark at 4°C (ready for imaging).

The FISH images were acquired with Zeiss Laser Scanning Confocal microscope (LSC780) with 63x magnification oil immersion objective lenses. Cells are randomly chosen (each vision field contains 6-15 cells). Signals from four different fluorophores were obtained with two alternative frame scans for best separation: the first scan with two laser beams of 488 nm and 594 nm, followed by the second scan of 405 nm (for DAPI) and 532 nm laser beams (for the yellow probe used in targeted group) or 405 nm and 555 nm laser beams (for the orange probe used in control group). The minimal laser power was used in combination with appropriate filter settings (MBS 488/594 and MBS 458/514/561/633) to greatly reduce the signal bleed through between channels. Images of cells with optical Z sections from the bottom to the top with 0.25 μm or 0.3 μm intervals were acquired one section after another (frame scanning) with the software Zen provided by the manufacturer. Signals of each probe were stored in separate channels (four channels for three chromosomal regions plus DAPI staining of the whole chromosomal DNA).

The nucleus detection and distance measurements between probes were performed using the Nemo software for FISH image analyses[19]. Automated nucleus detection mode was used as the standard procedure, and additional manual selection followed when needed. Each of the cells was subject to manual inspection and validated for containing at least a set of the closest three probes.
